# Trait-space disparity in fish communities spanning 380 million years from the Late Devonian to present

**DOI:** 10.1101/2024.08.29.610219

**Authors:** John Llewelyn, John A. Long, Richard Cloutier, Alice M. Clement, Giovanni Strona, Frédérik Saltré, Michael S. Y. Lee, Brian Choo, Kate Trinajstic, Olivia Vanhaesebroucke, Austin Fitzpatrick, Corey J. A. Bradshaw

## Abstract

The diversity and distribution of species’ traits in an ecological community determine how it functions. While modern fish communities conserve trait space across similar habitats, little is known about trait-space variation through deep time or across different habitats. We examined how fish trait diversity varies through space and time by comparing three Late Devonian fish communities — a tropical reef (Gogo, Australia), a tropical estuary (Miguasha, Canada), and a temperate freshwater system (Canowindra, Australia) — with six modern communities from diverse habitats. Trait-space metrics reflecting within-community diversity (functional richness) and species similarity (functional nearest-neighbour distance) indicated Late Devonian communities had scores similar to modern communities. However, they were less functionally rich than their closest modern analogues, and their species tended to be more functionally distinct from one another. Metrics describing location in trait space (centroid distances and hypervolume overlap) showed modern communities were similar to each other, Gogo and Miguasha were similar but distinct from modern communities, and Canowindra was distinct from all others. This pattern suggests period-associated differentiation and substantial heterogeneity among some Late Devonian communities. In addition to temporal changes, we found consistent differences associated with habitat type and climate zone. Reef and tropical communities were the most functionally rich, whereas functional nearest-neighbour scores were highest in estuarine and temperate communities. These results indicate fish community trait space varies with time, habitat and climate, suggesting (*i*) lability in fish trait space and (*ii*) that evolutionary history, environmental filtering, and stochasticity influence community assembly.

## 1. Introduction

The combination of a species’ traits mediates its interactions with its environment (Kiørboe et al., 2018; McGill et al., 2006), so the range and distribution of traits among interacting species is important for determining function and resilience of ecological communities (Cadotte, 2017; McLean et al., 2019; Micheli et al., 2014). Further, the diversity and distribution of traits in a community — also known as community ‘trait space’ — can provide insights into the rules of community assembly, including what roles evolution, species filtering, and niche differentiation play in controlling how communities form (Mouillot et al., 2021; Schöb et al., 2012; Weiher et al., 1998). Indeed, recent research on the trait space of contemporary communities has addressed how communities develop and adjust to their ever-shifting environments (Frimpong and Angermeier, 2010; Vogel et al., 2019). However, it is still unclear whether patterns observed within and among modern communities also apply to ecological communities over deep time. Consistent spatial and temporal patterns in community trait space would suggest that community assembly rules are constant, whereas variation could indicate phylogenetic constraints (Kohli and Rowe, 2019), stochasticity in evolutionary pathways (e.g., random mutations providing trait variation and the stochastic extinction of clades; Chase and Myers, 2011; Zhou and Ning, 2017), and/or shifts in the environmental parameters that determine community composition (Powell et al., 2015).

A range of metrics are commonly used to describe and investigate community trait space (Mammola et al., 2021). Such trait-diversity metrics often indicate variation among communities in different habitats and climates (Ingram and Shurin, 2009; Pease et al., 2012). For example, trait diversity in fish (non-tetrapod vertebrate) communities differs depending on substratum (e.g., reef *versus* seagrass *versus* sand) (Henseler et al., 2019; Pecuchet et al., 2016; Rincón-Díaz et al., 2018) and habitat complexity (Quirino et al., 2021; Sgarlatta et al., 2023). Despite this variation, there is also evidence for the conservation of trait space across similar habitats, even when there are substantial phylogenetic and taxonomic differences among the resident communities (McLean et al., 2021). That evolutionarily distinct communities occupying similar habitats have similar trait spaces suggests convergence (through evolution and/or niche filtering) and highlights the importance of deterministic processes in shaping community trait space (i.e., environment determining trait space) (Triantis et al., 2022; Vellend, 2010).

Although trait space varies among habitats and is conserved within habitat types across locations in contemporary communities, it remains unclear how it varies through deep time. Few studies have examined changes in functional diversity and trait space over millions of years (but see Reeves et al., 2021 for impacts of extinction events). By testing for patterns across time as well as across contemporaneous habitats, it is possible to evaluate whether similar habitats select for similar community trait space irrespective of phylogeny and evolutionary history.

The Devonian Period (419.2 to 358.9 million years ago [Ma]) is popularly termed the “Age of Fishes”. During that time, diverse fish communities established and radiated across the planet, giving rise to the first tetrapods (Klug et al., 2010; Long, 2010). Since then, multiple mass extinction events have occurred, which along with the passing of time and millions of ‘background’ extinction and speciation events, have caused massive phylogenetic turnover (McGhee et al., 2013; Sallan and Coates, 2010). In addition, Devonian and modern fish communities differ in terms of the non-fish species and nutrient inputs that make up the wider ecological community (Beerling et al., 1998; Brett and Walker, 2002). For example, Devonian reefs were primarily built of stromatoporoid sponges, calcareous algae, and microbial communities rather than hexacorallian corals (but also included some rugose and tabulate corals) (Copper, 2011; Trinajstic et al., 2022), and ancient estuaries generally had low invertebrate and algal diversity compared to modern estuaries (although taphonomic biases might partly explain this lower observed diversity) (Cloutier, 2013; Gess and Whitfield, 2020). Although the environments Devonian fish communities experienced were physically similar to their modern counterparts, their vertebrate and non-vertebrate biota therefore differed greatly. By comparing these ancient assemblages to modern communities, it is possible to test if the physical environment is the primary determinant of fish community trait diversity across space and time, or whether biological, phylogenetic, and evolutionary differences associated with the passing of hundreds of millions of years have a dominant effect on trait space. Such knowledge will improve our understanding of how communities have responded to change in the past, and how they could change in response to anthropogenic and non-anthropogenic disturbances in the future.

We compared the trait space of three of the best-described Late Devonian fish communities — a tropical reef (Gogo Formation, north-western Australia), a tropical estuary (Miguasha, eastern Canada), and a temperate freshwater billabong (river backwater) system (Canowindra fish beds, south-eastern Australia) — to that of six modern fish communities from different habitats and climate zones. Our aim was to identify how trait diversity varies through space and time and thereby shed light on the processes of community assembly. We compared these communities in terms of two trait-space metrics that reflect diversity and distribution of traits within each community (functional richness [FRic] and functional nearest-neighbour distance [FNND]), and two that quantify each community’s position in trait space relative to other communities (distance between hypervolume centroids and overlap of hypervolumes) (Mouchet et al., 2010). We hypothesise that (*i*) if community trait space is primarily determined by habitat, there will be larger differences between communities from different habitats (i.e., reef *versus* estuary *versus* freshwater) than between Devonian and modern communities from the same habitat (e.g., Devonian reef community similar to modern reef community) or between tropical and temperate/subtropical communities from the same habitat (e.g., tropical reef similar to temperate reef). (*ii*) If community trait space is primarily determined by climate, there will be larger differences between tropical *versus* temperate/subtropical communities than between communities from different periods of time or different habitat types within the same climate zone. (*iii*) If community trait space is primarily determined by evolutionary history and phylogenetic turnover through time (includes stochasticity such as mutations generating trait variation and random extinction of phylogenetic clades), there will be larger differences between Devonian and modern communities compared to between communities from different habitat types or different climate zones within the same period. (*iv*) If habitat, climate and time all substantially affect community trait space, there will be consistent groupings/patterns across all these variables (e.g., community metric scores will group together based on time, habitat, and climate); and (*v*) If community trait space is independent of habitat, climate and time, trait space metrics will not be grouped by habitat, climate zone or time period (Devonian *versus* modern). By identifying which of these five patterns are observed in trait space metrics, we provide insight into the factors shaping community trait space (i.e., community assembly) in the past, present, and future.

## 2. Methods

### 2.1 Sites and species lists

We compared three Late Devonian (Frasnian to Famennian) and six modern fish communities. The Late Devonian communities of Miguasha (Escuminac Formation) in north-eastern Canada, the Gogo Formation in north-western Australia, and Canowindra fish beds (Mandagery Formation) in south-eastern Australia are recognised fish Lagerstätten from this epoch (Supplementary Fig. S1) (Cloutier and Lelièvre, 1998). We selected these communities based on (1) their fossil records capturing most of the *in situ* fish diversity, (2) the availability of trait data for a range of traits and species that make up these communities (Supplementary Table S1), and (3) the different habitat types they represent.

Miguasha (Escuminac Formation; modern: 48° 06′ N, 66° 21′ W) was a tropical estuary in Laurussia 379 to 375 million years ago (middle Frasnian; Chevrinais et al., 2017; Cloutier et al., 2011, 1996). Thousands of fish specimens have been recovered from Miguasha, representing at least 19 species (Cloutier et al., 2011). The Gogo Formation (modern: 18° 26′ S, 125° 55′ E) represents the basinal and channel facies adjacent to a tropical, stromatoporoid sponge-dominated reef that was on the edge of the Gondwanan landmass from the Givetian to the end of the Famennian (Playford et al., 2009). Although reef building continued throughout this interval, the fish-bearing nodules are known only from the Frasnian (Late Devonian) horizons and are estimated to be 384 to 382 million years old (Trinajstic et al. 2022). These fish were deposited and preserved under anoxic marine conditions and came from the adjacent reef (Trinajstic et al., 2022). Fifty-three fish species have been identified from Gogo (including 47 described and 6 undescribed species) (Supplementary Tables S1 and S2; Mory and Hocking, 2011; Long and Trinajstic, 2010; Long and Trinajstic, 2017, Trinajstic, et al., 2022). The Canowindra fish beds (Mandagery Formation; modern: 33° 35′ 94" S, 148° 33′99" E) are thought to have been deposited in a freshwater billabong (river backwater or oxbow lake) in a temperate, arid environment on Gondwana approximately 363 million years ago (Famennian stage) (Australian Heritage Council, 2012; Retallack, 2024). Over 3,000 fish specimens belonging to eight species have been recovered from these fish beds that are thought to have been created when the billabong dried up (Retallack, 2024). Although the Kellwasser Event (Frasnian-Famennian boundary, ∼ 372 million years ago) separates Canowindra from Miguasha and Gogo, that extinction event primarily affected marine communities — freshwater communities appear to have been less affected, with evidence suggesting continuity of diversity and composition across the event (Sallan and Coates, 2010).

Discovery curves suggest all (preserved) fish species from Miguasha have been described (Cloutier, 2013), and that most (> 95%) fish species from Gogo have been identified (Supplementary Fig. S2 and Table S2). Specimens were only collected from Canowindra twice (in 1956 and 1993) (Long, 2013) and so it is not appropriate to fit species discovery curves to these data. However, the Chao1 estimator extrapolated a species richness of 8.5 (5.5–11.5 95% confidence interval) for Canowindra, suggesting that all or most species have already been discovered for this community (Llewelyn et al., 2024). However, because these fish beds are believed to have been deposited in a single drying event, it is possible that this fossil assemblage does not represent the entire fish diversity in that community.

We selected six modern fish communities from a diversity of environments (different habitat types and climate zones) for comparison to the Devonian communities. Our selection of modern communities was further guided by the availability of species lists for the assemblage and the presence of information on trophic interactions (allowing for later development of trophic-network models of those communities). We included reef, estuarine, and freshwater communities, with tropical and temperate or subtropical representatives of each (i.e., we included each habitat-type/climate-zone combination). The modern communities were: Caribbean reefs (tropical; 18° 3′ N, 65° 28′ W) (Bascompte et al., 2005); Chilean reefs (temperate/subtropical; 31° 30′ S, 71° 35′ W) (Pérez-Matus et al., 2017); Santa Cruz Channel, north-eastern Brazil (tropical estuary; 7° 46′ S, 34° 53′ W) (Ferreira et al., 2019; Lira et al., 2022); Ythan Estuary, Scotland (temperate; 57° 19′ N, 1° 59′ W) (Cohen et al., 2009; Hall and Raffaelli, 1991; Huxham et al., 1996); Braço Morto Acima and Abaixo, central Brazil (tropical freshwater oxbow lakes; 19° 41′ S, 56° 59′ W) (Angelini et al., 2013; Costa-Pereira et al., 2011; de Resende, 2000; de Resende and Pereira, 2000a, 2000b, 1998; Ferreira et al., 2017; Pereira and de Resende, 1998; Severo-Neto et al., 2015); and the Nepean River, New South Wales, Australia (temperate/subtropical freshwater river section; 33° 47′ S, 150° 38′ E; ala.org.au; Supplementary Table S3) (Growns et al., 2003).

### 2.2 Traits

We collected information on 11 traits from the 496 species in our study (416 extant species and 80 Devonian species; Supplementary Table S4). We chose traits based on their ecological relevance and their availability for both living and extinct species. To avoid introducing bias into trait-space patterns due to trait selection, we excluded traits that were specific to one time period (e.g., we did not include bony armour or protrusible jaws), focusing instead on general traits applicable across deep time. The traits we chose were: sagittal body shape (i.e., lateral profile), transverse body shape, maximum total body length (tip of snout to tip of caudal fin), head length, eye diameter, pre-orbital length, body depth, position of mouth, position of eyes, presence of spiracle, and caudal fin shape (Supplementary Fig. S3 and Table S4). We used traits for adult or large specimens of each species because although intraspecific trait variation can influence trait space (Palacio et al., 2025), collecting such data on the many extant and extinct species in our study was not logistically feasible. Therefore, we focused on adult or large specimens and took the mean value when multiple measurements were available. We expressed all morphometric variables (except maximum total body length) as a proportion of maximum total body length as a size standardisation and to reduce correlations among these traits.

We collected trait data for Devonian species from the literature, specimens, and photographs of specimens. However, it was not possible to collect all traits for all species from Gogo and Canowindra because some of these species are only known from incomplete or disarticulated body parts (Long and Trinajstic, 2010). We addressed these gaps by estimating traits using: (1) expert opinion in cases where traits could be confidently inferred (inferences made by J. Long, A. Clement, B. Choo, and K. Trinajstic) or (2) multiple imputation using species’ traits, coarse taxonomy, and the missForestR package (proportion of trait data imputed: < 20% and < 16% for Gogo and Canowindra, respectively) (Llewelyn et al., 2024; Stekhoven, 2022). Evaluating multiple imputation performance indicated out-of-bag error estimates of 0.14 (normalised root-mean-square error) and 0.09 (proportion of falsely classified) for continuous and categorical variables, respectively. For modern species, we extracted trait data from *Fishbase* (using the rFishbase package) (Boettiger et al., 2012), photographs, and scientific literature (Llewelyn et al., 2024). We visually inspected the distribution of continuous traits, identifying two as right-skewed (total length and pre-orbital length); we log_e_-transformed those traits to normalise them.

### 2.3 Analyses

We built a Gower dissimilarity matrix (Gower and Legendre, 1986) to quantify trait differences among species using the gawdis package in R (de Bello et al., 2021). Gower dissimilarity can handle mixed data types (our trait data included continuous and categorical variables) and can normalise the contribution of variables (Gower and Legendre, 1986; Mouillot et al., 2021). In addition to including all traits when calculating Gower distances, we did sensitivity analyses using (*i*) only the most accurately imputed traits (numeric traits with a normalised root mean-squared error < 0.3, and categorical traits with piecewise constant fitting < 0.25), and (*ii*) using only morphometric traits. Observed patterns in functional diversity metrics were consistent across these trait sets (Supplementary Fig. S4 and S5). We therefore only present results from the ‘all traits’ Gower distance trait set in the main text.

We applied principal coordinate analysis (PCoA) to the Gower distances to ordinate species within a lower-dimensional space and calculate functional diversity metrics using the mFD R package (Magneville et al., 2022). The first seven principal coordinate axes explained 84% of the total variation; we identified that seven was the optimal number of axes based on mean absolute deviation and root mean-squared error (minimising the deviation between trait-based distances and functional space distances) (Llewelyn et al., 2024). This was also the maximum number of axes we could use because the smallest community consisted of only eight species (the number of axes must be less than the number of species). We therefore used these seven axes to calculate each community’s observed functional richness (FRic) (Mouillot et al., 2013) and observed functional nearest neighbour (FNND) (Magneville et al., 2022) scores. Functional richness, defined as the proportion of functional space filled by a community, quantifies the functional diversity of a community. In contrast, functional nearest neighbour is the average distance in trait space to each species’ nearest neighbour within the community, quantifying functional similarity (redundancy) among species in a community.

These metrics, calculated from PCoA coordinates, are unitless ‘scores’ that can be used to compare communities. Additionally, we calculated functional evenness, functional divergence, and functional specialisation, which indicate (respectively) how regularly species are distributed in trait space, how distinct species are from the centroid of the community’s trait space, and how distinct species in a community are from the global centroid. We present these three metrics in the Supplementary Material (Supplementary Fig. S6).

Species diversity varied substantially among the included communities, ranging from 8 species in Canowindra to 208 species in Caribbean reefs (Supplementary Tables S1 and S3). Functional diversity metrics can be correlated with species diversity (Mammola et al., 2021; Ricklefs and Miles, 1994). We therefore calculated standardised effect sizes for both metrics, in addition to reporting the observed (raw) scores. This method controls for species richness, and we hereafter refer to these adjusted scores as ’standardised functional richness’ and ’standardised functional nearest-neighbour’ (Hurtado-Materon and Murillo-García, 2023). To calculate these scores, we generated 1000 null models by randomly shuffling species among the communities while maintaining the species richness of each community. We then calculated the mean and standard deviation of the functional diversity metrics for each community in the null models. We subtracted null mean values from observed functional diversity metrics to calculate the effect sizes, which we divided by the null standard deviations to standardise across scales. Controlling for species diversity in this way allowed us to (*i*) separate patterns due to species diversity from those due to functional diversity, (*ii*) detect patterns in functional diversity that were masked by species diversity, and (*iii*) limit the effects of species with extreme traits. We plotted both the ‘observed’ and ‘standardised’ (species-diversity controlled) metrics, allowing us to evaluate visually whether period, habitat type, or climate zone grouped communities in either of these metrics.

We also used the principal coordinates to build trait-space hypervolumes for each community using Gaussian kernel density estimation and the hypervolume R package (Blonder et al., 2023). We set the requested quantile to 0.95 and estimated the bandwidth for each axis in each community separately. We then used the mean estimated bandwidth for each axis across the nine communities as fixed bandwidths to recalculate hypervolumes (i.e., ensuring the same bandwidths for each community). We used the resulting hypervolumes to quantify distances between community centroids (geometric centre) and overlap in multidimensional trait space. We measured overlap using the Jaccard index, and we further decomposed non-overlap into its turnover and nestedness components (Baselga, 2012). We made these comparisons between all pairwise community combinations to test whether communities were grouped/clustered according to their period, habitat, or climate zone (see hypotheses listed in the Introduction). Because distances between centroids suggested the fish community trait spaces were segregated, we identified the specific traits responsible by determining along which hypervolume axes the centroids differed, and which traits were correlated with these axes. We then investigated how these traits were distributed among the fish communities. All data and code we used in the analyses are available online (Llewelyn et al., 2024).

## 3. Results

Devonian communities had lower observed functional richness than their closest modern counterparts (Fig. 1a; Gogo *vs.* Caribbean: 7.5 × 10^-3^ *vs.* 0.29; Miguasha *vs.* Santa Cruz Estuary: 2.3 × 10^-3^ *vs.* 4.7 × 10^-2^; Canowindra *vs.* Nepean: 7.9 × 10^-6^ *vs.* 1.7 × 10^-4^). However, standardised functional richness (controlling for species richness) of Devonian communities was similar to that of their modern counterparts (Fig. 1b; Gogo *vs.* Caribbean: -2.7 *vs.* -2.2; Miguasha *vs.* Santa Cruz Estuary -0.8 *vs.*-2.3; Canowindra *vs.* Nepean River: -0.4 *vs.* -1.2).

**Figure 1.**
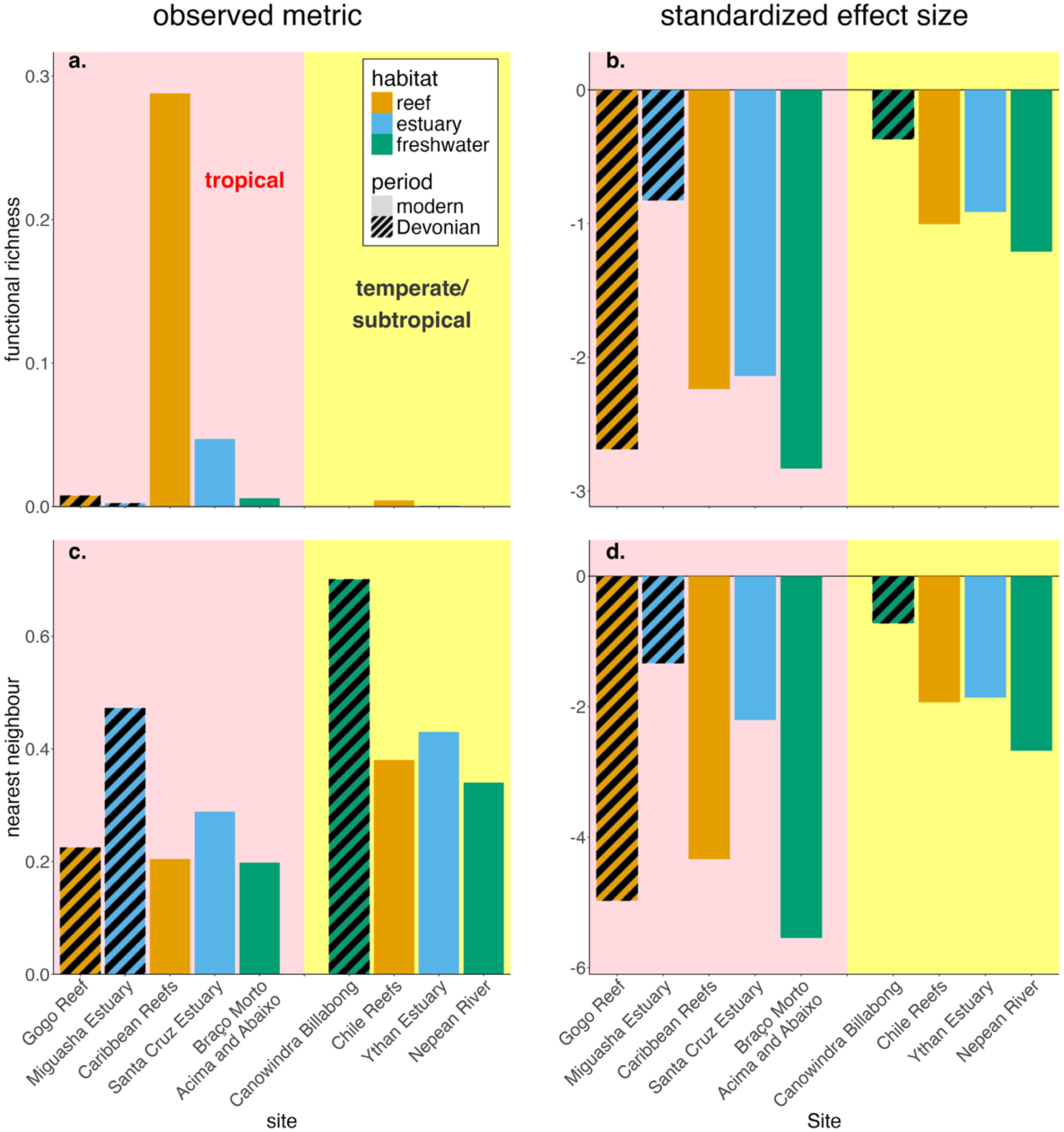
Functional diversity metrics for three Devonian and six modern fish communities. Two functional diversity metrics are shown: functional richness (a. and b.) and functional nearest neighbour (c. and d.). Panels in the left column show the observed (raw) metrics (not controlling for species diversity), whereas panels in the right column show standardised effect sizes (controlling for species diversity). Pink background indicates the five tropical communities (ancient and modern), yellow background indicates the four temperate/subtropical communities (ancient and modern). Devonian communities are indicated with black diagonal lines.

Functional richness varied among habitats, although the pattern differed depending on whether we controlled for species diversity (Fig. 1a,b). Within each time period and climate zone, reef communities had the highest observed functional richness followed by estuarine communities (Fig. 1a). In contrast, for standardised functional richness, the Devonian freshwater community (Canowindra) had the highest score, the modern freshwater communities the lowest, and estuarine communities had higher scores than the reef communities (Fig. 1b). This switching between reefs and estuaries being the more functionally rich when controlling for species diversity emerged among both modern and Devonian communities (Fig. 1a,b). Similarly, functional richness varied with climate, and the direction of this pattern depended on whether we controlled for species diversity. Tropical communities had higher observed functional richness than their temperate/subtropical counterparts (Fig. 1a; mean richness tropical *vs.* temperate/subtropical: 0.11 *vs.* 0.0006), whereas temperate and subtropical communities had higher standardised functional richness (Fig. 1b; mean richness tropical *vs.* temperate/subtropical: -2.1 *vs.* -0.9).

Observed nearest-neighbour scores, reflecting the distance in functional space of each species to the nearest species in its community, tended to be higher for Devonian than modern communities (mean nearest neighbour Devonian *vs.* modern: 0.47 *vs*. 0.31; Fig. 1c,d). Canowindra and Miguasha had higher observed scores than the other communities, and Gogo’s score — although similar to that of modern communities — was slightly higher than its modern counterpart (Fig. 1c; Canowindra = 0.7, Miguasha = 0.47, modern assemblages = 0.20 to 0.43; Gogo *vs.* Caribbean Reefs: 0.22 *vs.* 0.20). Standardised nearest-neighbour scores for Canowindra and Miguasha remained higher than other communities, whereas Gogo’s score was slightly lower than its modern counterpart (Fig. 1d; Miguasha = -1.3, Canowindra = -0.7; modern assemblages = -5.5 to -1.9; Gogo *vs.* Caribbean Reefs: -5.0 *vs.* -4.3).

There were differences among habitats in nearest-neighbour scores within periods and climate zones. Among modern assemblages, estuarine communities consistently (for both observed and standardised scores) had the highest nearest-neighbour scores followed by reef communities (Fig. 1c,d). Similarly, the Devonian estuarine community had a higher score than the Devonian reef community (Fig. 1c,d). However, unlike the modern assemblages, the Devonian freshwater community (Canowindra) had the highest nearest neighbour score of the communities in that period (its score was also higher than the modern communities; Fig. 1c,d). In addition to these habitat-linked differences, there was a consistent latitudinal pattern among communities — species in temperate and subtropical communities were farther from their nearest neighbour than were species in tropical communities (Fig. 1c,d; mean nearest neighbour tropical *vs.* temperate/subtropical: 0.28 *vs.* 0.46 [observed] and -3.6 *vs.* -1.8 [standardised]).

The distances between hypervolume centroids of Devonian and modern communities tended to be greater than those between centroids of communities from the same period (Fig. 2a,b; distance between Devonian and modern: 0.16 ± 0.03, distance between communities from the same period: 0.07 ± 0.03; reported values are means ± standard deviations unless otherwise stated). This suggests that community trait spaces were centred in different locations depending on time period. Although Canowindra was closer to the two other Devonian communities than to the modern communities, its centroid distances to Gogo and Miguasha (0.15 and 0.12, respectively) were still higher than those observed in other same-period comparisons (0.03–0.09; Fig. 2). Furthermore, distances between Canowindra and modern communities (0.171–0.206) were higher than those of other inter-period comparisons (0.126–0.165). Centroids of communities from the same habitat type or the same climate zone were not closer together than centroids from different habitats or climate zones (Fig. 2c,d).

**Figure 2.**
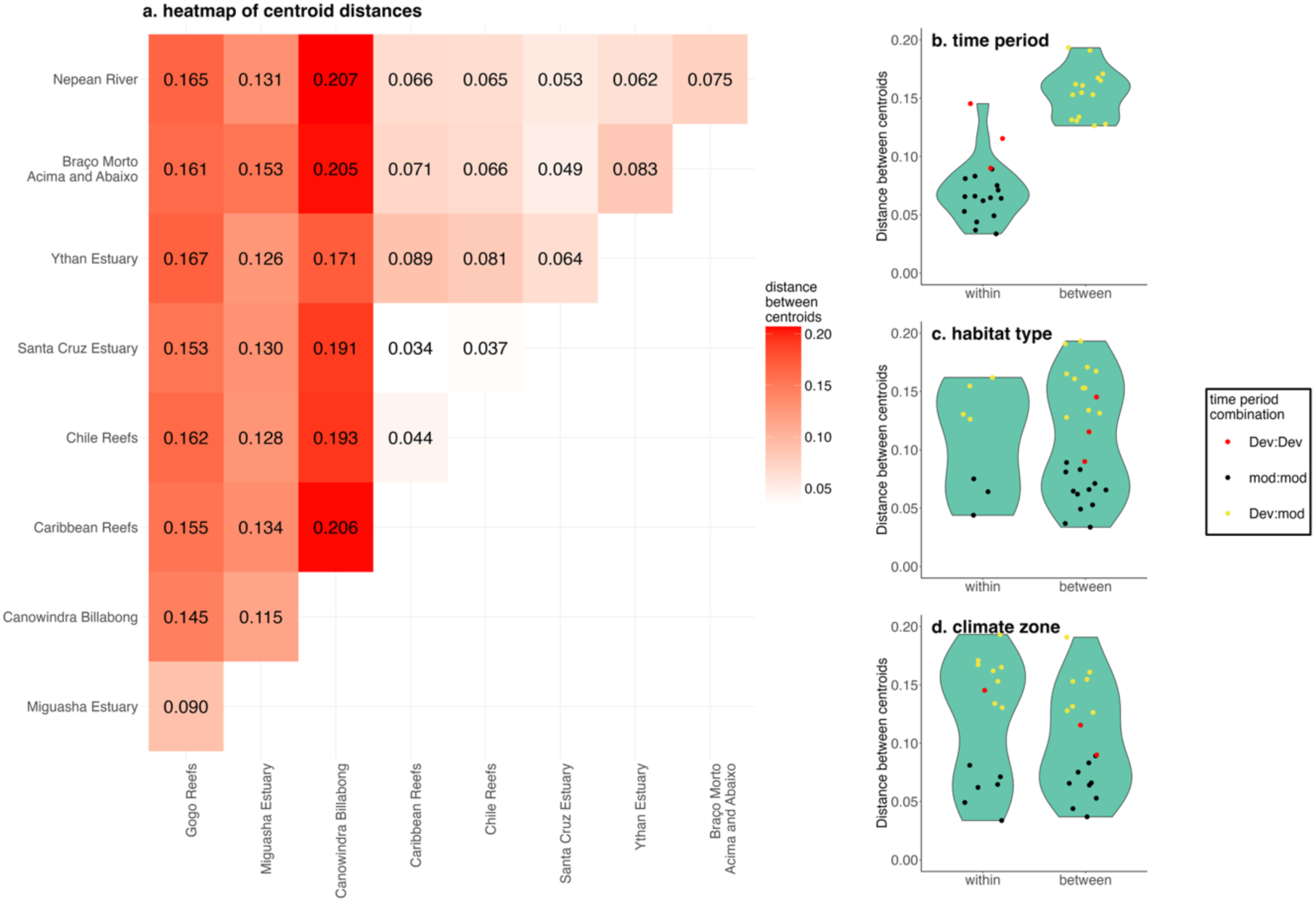
Distances between centroids of functional trait space hypervolumes for fish communities. We compared three Devonian (Gogo, Miguasha, and Canowindra) and six modern fish communities from different habitats and climates. Plot a. (left) is a heatmap showing distances between the centroids of each community. Plots b., c., and d. (right) are three violin plots showing distances between communities from either the same (within) or different (between): b. periods, c. habitat types, and d. climate zones. Points in the violin plots represent individual comparisons between communities, jittered on the *x* axis for clarity.

The hypervolume Jaccard index measuring trait-space overlap also indicated less similarity between Devonian and modern communities compared to between communities from the same period (Fig. 3a,b; overlap between Devonian and modern: 0.07 ± 0.04, overlap between communities from the same period: 0.25 ± 0.11). Canowindra’s overlap with other communities followed a similar pattern to that observed with centroid distances: although it had low overlap with all other communities, it tended to overlap more with the other Devonian communities — Miguasha and Gogo (0.035 and 0.03, respectively) — than with modern communities (Fig. 3). Canowindra overlapped with only one modern community as much as it overlapped with the Devonian communities (Canowindra *vs.* Chile Reefs: 0.035; Fig. 3). There was no evidence of greater overlap between hypervolumes of communities from the same *versus* different habitat types or climate zones (Fig. 3c,d). Partitioning the Jaccard index into components of dissimilarity due to turnover *versus* nestedness indicated that turnover largely explained differences among communities, i.e., these communities occupied different trait spaces rather than subsets of each other’s trait spaces (dissimilarity due to turnover: 0.35–0.99; dissimilarity due to nestedness: 0–0.47; Supplementary Figs. S7 and S8).

**Figure 3.**
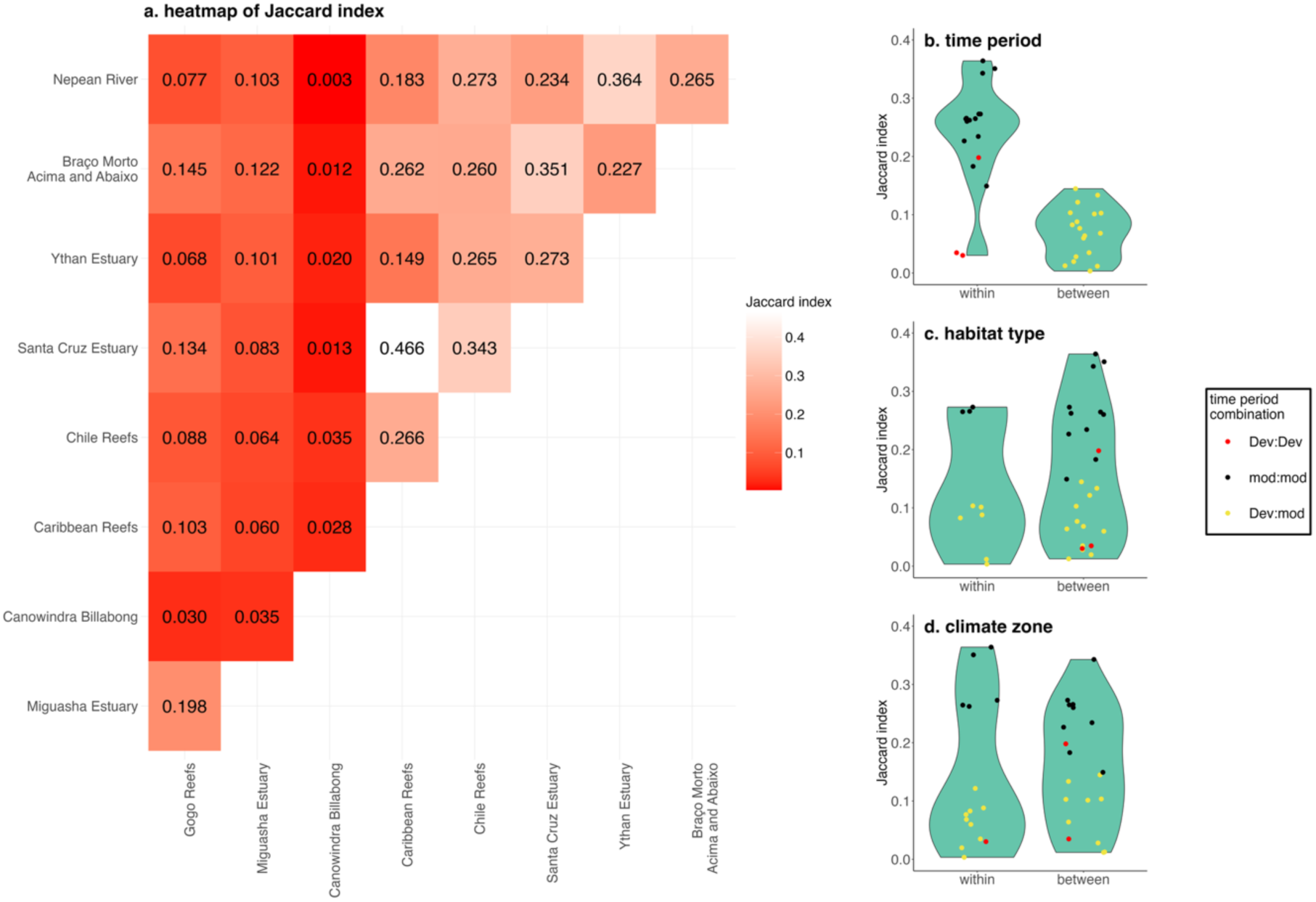
Jaccard Index comparing overlap in functional trait space hypervolumes for fish communities. We compared three Devonian (Gogo, Miguasha, and Canowindra) and six modern fish communities from different habitats and climates. Plot a. (left) is a heatmap showing Jaccard indices for all community comparisons. Plots b., c., and d. (right) are violin plots showing Jaccard indices between communities from either the same (within) or different (between): b. periods, c. habitat types, and d. climate zones. Points in the violin plots represent individual comparisons between communities, jittered on the *x* axis for clarity. See supplementary Figures S7 and S8 for the turnover and nestedness components of the Jaccard Index.

The centroids of all three Devonian communities were outside the range observed in modern communities on axis 1 (Devonian ≤ -0.079, modern ≥ -0.043) and axis 7 (Devonian ≤ -0.026, modern ≥ 0.004; Supplementary Table S5 and Fig. S9). Although the principal coordinate axes are based on Gower distances between species and cannot be directly linked to specific traits, correlations between species coordinates along these axes and traits suggest which traits they reflect. Axis 1 was most strongly correlated with eye diameter (correlation: *η*^2^ = 0.622, p < 0.0001), body depth (*η*^2^ = 0.523, p < 0.0001), transverse body shape (*η*^2^ = 0.359, p < 0.0001), and caudal fin shape (*η*^2^ = 0.349, p < 0.0001; Supplementary Table S6). Transverse body shape was also correlated with axis 7 (*η*^2^ = 0.262, p < 0.0001), as was sagittal body shape (*η*^2^ = 0.348, p < 0.0001) and presence/absence of spiracles (*η*^2^ = 0.255, p < 0.0001). Accordingly, Devonian communities were distinct from modern communities in these traits. Fish in the Devonian communities tended to have smaller eye diameters, their bodies were not as deep, and they displayed less variation in these traits compared to fish in the modern communities (eye diameter relative to total body length: 0.037 ± 0.017 *vs.* 0.052 ± 0.023; body depth relative to total body length: 0.190 ± 0.052 *vs.* 0.257 ± 0.113, for Devonian and modern communities, respectively; Fig. 4c,e). The proportion of fish with a compressed transverse body shape was lower in the Devonian than modern communities (23% *vs.* 55%; Fig. 5c), whereas a circular transverse body shape was more common in the Devonian (43% *vs.* 12%; Fig. 5c). Homocercal tails were more common in modern than Devonian fish (92% *vs.* 3.8%; Fig. 5d), whereas heterocercal tails were more common in the Devonian communities (80% *vs.* 3%; Fig. 5d). Spiracles were also more prevalent among the Devonian fish (spiracles: 58% *vs.* 2%; Fig. 5b). Similarly, fusiform sagittal body shapes were more common among Devonian fish (60% *vs.* 46%; Fig. 5a), while ‘short and/or deep’ body shapes were more prevalent in the modern communities (23% *vs.* 14%; Fig. 5a).

**Figure 4.**
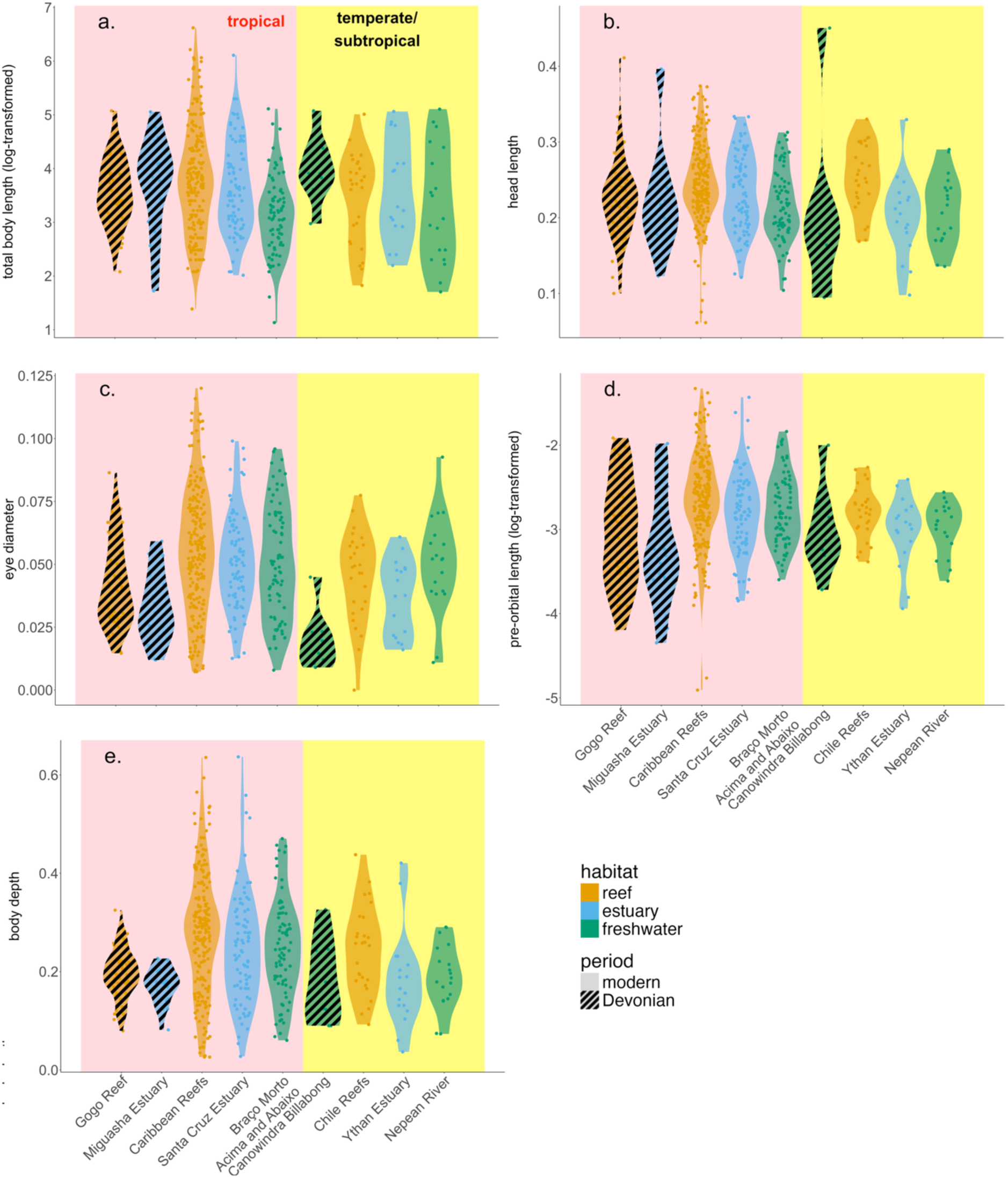
Violin plots of five continuous fish traits from three Devonian and six modern communities. We transformed total body length (cm) using the natural logarithm (log*_e_*) to normalise. We expressed all other traits as a proportion of total body length (log*_e_*-transformed for pre-orbital length). Pink background indicates the five tropical communities (ancient and modern), yellow background indicates the four temperate/subtropical communities (ancient and modern). Devonian communities are indicated with black diagonal lines.

**Figure 5.**
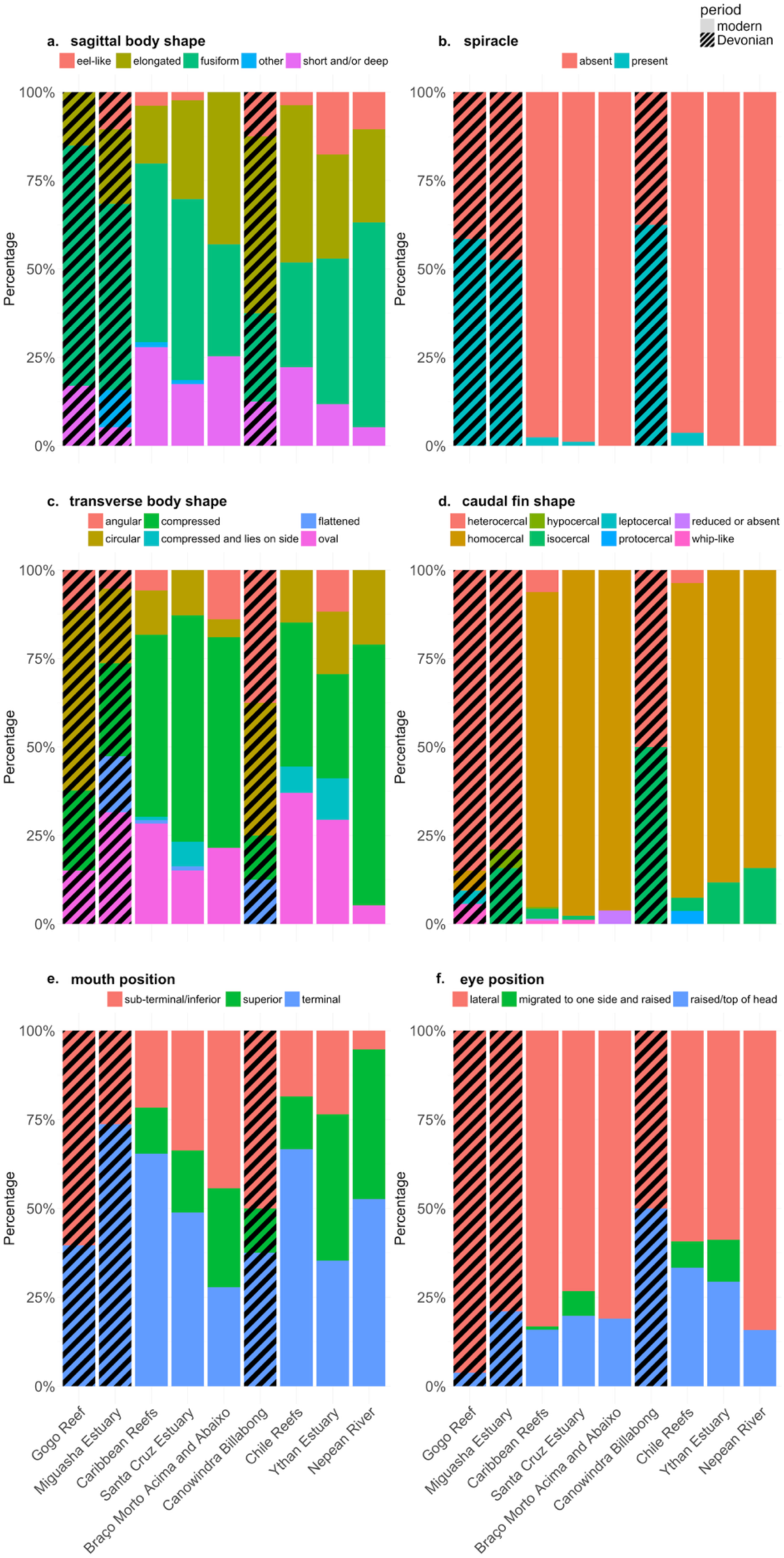
Stacked bar charts of six categorical fish traits from three Devonian and six modern communities: a. sagittal body shape, b. presence/absence of spiracles, c. transverse body shape, d. caudal fin shape, e. mouth position, and f. eye position. The *y* axis indicates the percentage of species in each community with each category of the trait. Devonian communities are indicated with black diagonal lines.

Miguasha and Gogo were outside the range observed in modern communities on axis 2 (Miguasha and Gogo ≥ 0.051, modern ≤ 0.007) and axis 5 (Miguasha and Gogo ≤ -0.001, modern ≥ 0.006), and Miguasha’s centroid was extreme on axis 4 (Miguasha = -0.048, modern ≥ -0.002). Axis 2 was associated with position of mouth (*η*^2^ = 0.479, p < 0.0001) and eye position (*η*^2^ = 0.376, p < 0.0001). Position of mouth also strongly correlated with axis 4 (*η*^2^ = 0.479, p < 0.0001), as did sagittal body shape (*η*^2^ = 0.271, p < 0.0001), while axis 5 most strongly correlated with sagittal body shape (*η*^2^ = 0.571, p < 0.0001). Reflecting these associations, Gogo and Miguasha were distinct from other communities in these traits.Neither Gogo nor Miguasha had fish with superior mouth position, whereas all other communities did (Fig. 5e). None of the Devonian fish had eyes migrated to one side of the body (as in flatfish; Fig. 5f). Gogo lacked eel-shaped fish, while Miguasha had a higher occurrence of fish with unconventional (‘other’) body shapes, which included the tadpole-shaped *Escuminaspis laticeps* and *Levesquaspis patteni* (Fig. 5a). Although centroid positions did not suggest consistent differences in relative head lengths between Devonian and modern communities, each Devonian community included at least one species with a longer relative head length than found in the modern communities (Fig. 5b).

## 4. Discussion

Our analysis of modern and ancient fish communities revealed patterns in community trait space associated with time period, habitat type, and climate zone. These associations imply that both evolutionary history and environmental factors are important in determining community composition. However, the importance of time, habitat type, and climate zone appear to have differed among trait-space metrics, indicating that different factors are important for different trait-space attributes. The two metrics that reflect diversity and distribution of traits within a community — functional richness and functional nearest neighbour distance — suggest: (1) broad similarity between Late Devonian and modern assemblages, but with Late Devonian communities generally exhibiting lower functional richness and higher functional nearest-neighbour scores than their closest modern counterparts when not accounting for species diversity, (2) an association with habitat type, and (3) an association with climate zone (Fig. 1; consistent with hypothesis *iv* — that time, habitat, and climate all substantially affect community trait space). In contrast, time period was the only variable that differentiated fish assemblages in the two metrics that reflect a community’s position in trait space relative to other communities (distance between hypervolume centroids and Jaccard index); we observed no associations between these metrics and habitat type or climate zone (Figs. 2 and 3; consistent with hypothesis *iii* — that trait space is primarily determined by evolutionary history). The trait-space positions of Late Devonian communities differed from modern communities on several axes associated with nine of the eleven traits we included in our analyses (Supplementary Tables S5 and S6); such trait differences could indicate disparity between Devonian and modern communities in terms of their structure and function.

### 4.1 Period

If fish community trait space has changed through time, we would expect to see communities from different periods segregated in terms of their trait-space metrics (i.e., modern communities similar to each other and different to Devonian communities). This pattern is displayed in the four metrics we calculated: Devonian communities differ from modern communities in terms of functional richness, functional nearest neighbour distance, centroid location, and hypervolume overlap (Figs. 1a, 1c, 2, 3).

Functional richness of Devonian communities was similar to that of modern communities, but the observed scores (not controlling for species diversity) of Devonian communities in this metric were substantially lower than those in modern communities from the same habitat type and climate zone (i.e., Gogo Reef *versus* Caribbean Reef, Miguasha *versus* Santa Cruz Estuary, Canowindra *versus* Nepean River; Fig. 1a). Despite the Devonian period being popularly named the ‘Age of Fishes’, Devonian fish communities tended to have lower species diversity compared to modern fish communities (although still higher than in earlier periods) (Friedman and Sallan, 2012). Our results suggest that this lower species diversity was associated with lower trait diversity compared to their modern counterparts (Fig. 1a *vs.* b). Discovery curves and the Choa1 estimator confirm that most species in the Devonian communities we examined have already been described (Supplementary Table S2 and Fig. S2) (Cloutier, 2013; Llewelyn et al., 2024) and therefore, the relatively low observed functional and species richness in Devonian communities compared to their modern analogues are probably real features instead of preservational artefacts. When we controlled for the effect of species diversity on functional richness using standardised effect sizes, Devonian communities had functional richness scores that were similar or higher than their modern counterparts (Fig. 1b). This result emphasises that changes in trait diversity between time periods are associated with changes in species diversity, although it remains unclear whether higher trait diversity is a cause or effect (or both) of higher species’ diversity.

Lower functional richness in Devonian fish assemblages does not necessarily mean that more trait space was vacant in these communities. It is possible that non-fish taxa filled some of the space occupied by fish in modern communities. For example, conodonts (early, non-fish, but fish-like, jawless vertebrates) might have occupied the trait space of modern lampreys, hagfish, and small eels (Aldridge and Donoghue, 1998; Aldridge and Purnell, 1996), while invertebrates such as eurypterids (sea scorpions) and cephalopods might have filled trait and trophic spaced occupied by predatory fishes in modern communities (McCoy et al., 2015; Greif et al. 2022). Similarly, non-fish taxa, such as sea snakes, turtles, and marine mammals, make unique contributions to the trait space in modern aquatic communities (Pimiento et al., 2020).

Miguasha and Canowindra consistently had the highest observed and standardised functional nearest-neighbour scores (Fig. 1c,d), suggesting greater trait differentiation and lower functional redundancy among fish in these Devonian communities than in modern communities. Gogo had a similar (but slightly higher) observed nearest-neighbour score than its closest modern counterpart (Fig. 1c). That Miguasha and Canowindra had substantially higher scores and Gogo only differed marginally suggest a possible interaction between habitat and time period. If such an interaction is real, one plausible explanation for the distinctiveness (high nearest-neighbour scores) of fish within Late Devonian estuarine and freshwater communities is that these environments had only recently been colonised by vertebrates — with the earliest evidence of fish in brackish and freshwater environments dating to the Middle and Late Silurian, respectively (Halstead and Lawson, 1985; Jiang and Dineley, 1988). Fish might have still been diversifying and filling available trait space in these habitats, whereas Late Devonian marine communities were already more saturated.

Centroid distances and the Jaccard index (trait-space overlap) are metrics that reflect the location of a community’s trait space relative to other communities. These metrics show that modern communities are similar to each other, but distinct from the Devonian communities, i.e., they occupy different areas of trait space (Fig. 2 and 3). Although Miguasha and Gogo are similar to each other, Canowindra is distinct from all communities — especially from modern ones (Fig. 2 and 3). This pattern suggests that Devonian communities, and particularly Canowindra, exhibited greater inter-community variation in trait space than modern communities, which were more clustered and similar to each other. The observation that Devonian and modern communities occupy different areas of trait space indicates variation in trait values (continuous traits), trait status (categorical traits), and/or trait combinations.

By examining which hypervolume axes separated modern and Devonian communities, and which traits were correlated with these axes, we were able to identify the traits involved in differentiating the trait spaces of these communities. Nine of the eleven traits we included in our analysis appear to contribute to this differentiation, including eye diameter, body depth, head length, presence of large spiracles, transverse body shape, caudal fin shape, sagittal body shape, mouth position, and eye position (Fig. 4 and 5). Of these traits, eye diameter, body depth and eye position had lower diversity in Devonian compared to modern communities, whereas the only trait that showed greater diversity among Devonian fish was head length (Fig. 4 and 5). Thus, differences between modern and Devonian communities as indicated by centroid distances and the Jaccard index reflect, at least in part, trait space that is absent in the Devonian fish, which is consistent with the observed differences in functional richness between communities from these periods (Fig. 1a,b). However, decomposition of the Jaccard index dissimilarity suggested differences between Devonian and modern fish community trait spaces were primarily due to turnover rather than nestedness (Supplementary Figs. S7 and S8). In other words, Devonian communities occupied different trait spaces to modern communities rather than subsets of those trait spaces. This result is somewhat counterintuitive because the modern communities largely encompass the trait variation found in the Devonian communities when traits are considered individually (Fig. 4 and 5). However, some trait categories were unique to Devonian communities (e.g., hypocercal and leptocercal caudal fins; Fig. 5), and some Devonian species had longer heads than any found in the modern communities (Fig. 4b). Thus, the dominance of turnover in explaining trait space dissimilarity likely reflects both unique trait combinations among Devonian fish and some unique trait values.

Community trait space is linked to community structure and function (Le Bagousse-Pinguet et al., 2021; Schleuning et al., 2020). Our trait space results therefore suggest substantial ecological differences between Devonian and modern fish communities. For example, the dearth of Devonian fish with a superior mouth position — an adaptation to feeding on items higher in the water column from below (Brind’Amour et al., 2011; Moyle and Cech, 2004) — indicates differences in trophic interactions. While superior mouth position was observed in one Devonian fish in the three communities we studied (and has been documented in other Devonian species) (Janvier, 1996; Jobbins et al., 2024), the rarity of this trait in the Devonian contrasts with its widespread occurrence in modern communities (Fig. 5). Greater diversity of mouth position does not only indicate a greater variety of feeding strategies used today, but also suggests that species in modern communities experience different predation pressures to those present in the Devonian (e.g., in terms of predator archetypes) (Ehlman et al., 2019), which could lead to the evolution of distinct antipredator strategies. Similarly, the lack of fish with deep, laterally compressed bodies in the Devonian communities (i.e., fish from these communities had body depths relative to their total length < 33%, and only 23% of species were laterally compressed; Figs. 4 and 5) likely indicates ecological differences associated with microhabitat use and how a species interacts with conspecifics and other species (Kelley et al., 2013; Schakmann and Korsmeyer, 2023). Deep-bodied, laterally compressed jawless fishes, like some thelodonts, have been documented in older communities (Late Silurian and Early Devonian) (Wilson and Caldwell, 1993), but they were less widespread compared to deep-bodied fish in modern communities, and they appear to have gone extinct before the Late Devonian (i.e., the epoch we examined). Thus, the relatively low variation of traits in Late Devonian communities suggests they were less ecologically/functionally diverse overall than modern fish communities. However, the rapid increase in species richness of nektonic fish during the Devonian could have been a precursor to trait diversification, despite later diversity losses during the Late Devonian mass extinctions (Klug et al. 2010; Friedman and Sallan, 2012).

The greater (raw) trait variation we observed in modern *versus* Devonian fish communities could result from (*i*) greater species diversity (though this could be a cause or effect of trait diversity), (*ii*) increased habitat/environmental complexity through time (Girard and Renaud, 2012), (*iii*) the longer time modern fish assemblages have had to evolve and diversify (Friedman and Sallan, 2012), and/or (*iv*) the morphological flexibility of teleost fish, a group that did not evolve until the Triassic (> 100 million years after the Devonian; although greater variation in actinopterygian traits started to emerge immediately following the Hangenberg extinction event at the end of the Devonian) (Henderson et al., 2023). In modern fish communities there is a positive correlation between habitat complexity and functional richness (Quirino et al., 2021; Richardson et al., 2017). Thus, given that Devonian habitats appear to have been less complex than modern habitats (e.g., even the most complex Devonian reefs tended to be less structurally complex than modern coral reefs) (Majchrzyk et al., 2024) and diversity of benthic/habitat-forming organisms was lower (Benton and Emerson, 2007)), this factor might contribute to the lower trait diversity in Devonian fish. Conversely, teleosts have several traits that foster morphological flexibility. For example, the prehensile mouth parts of teleosts give great flexibility in mouth-linked traits, leading to the evolution of different mouth positions and head shapes (Hill et al., 2018) — although relative head lengths tended to vary more among Devonian than modern fish in the communities we studied (Fig. 4b). Other attributes that promote morphological flexibility in teleosts and other osteichthyans — fish that are more prevalent in modern communities compared to Devonian ones — include the presence of a swim bladder for buoyancy control and bony endoskeletons, facilitating diversification in movement patterns and body shape (He et al., 2023; Witten and Hall, 2015).

While our results suggest higher trait diversity in modern fish communities, we acknowledge that the choice of traits and the groupings we used for categorical variables could influence conclusions regarding which taxa or communities are most diverse (Mouillot et al., 2021). For example, if we had focused on bony armour plates and lobe-paired fin structure, Devonian communities would be classified as more trait-diverse than modern communities (Long et al., 2018). However, the observed pattern of lower functional richness in Devonian communities persisted even when we restricted analyses to morphometric traits (i.e., basic body-dimension traits; Supplementary Fig. S5), suggesting the pattern is not an artefact of which traits we chose. Regardless, our main conclusion — that overall trait space of modern and Devonian communities differ — should hold true for any broad sample of traits across the entire phenotype.

### 4.2 Habitat

In addition to showing differences between periods, functional richness and nearest-neighbour scores displayed variation with habitat type. Nearest-neighbour scores were consistently higher for estuarine communities than reef communities, modern freshwater communities had the lowest scores, and the Devonian freshwater community (Canowindra) had the highest scores (Fig. 1c,d). Similarly, functional richness varied with habitat type, but the functional richness-habitat pattern depended on whether we controlled for species diversity (Fig. 1a,b). Across climate zones, reef communities had higher observed functional richness than estuarine communities, whereas estuarine communities had higher standardised (controlling for species diversity) functional richness than reef communities (Fig. 1a,b).

Freshwater communities had the lowest observed functional richness (Fig. 1c). Modern freshwater communities still had the lowest functional richness in the standardised scores, whereas the Devonian freshwater community had the highest (Fig. 1d).

The low functional richness and low nearest-neighbour scores in the modern freshwater communities we examined (i.e., indicating low trait diversity and high similarity among species; Fig. 1a,b,c,d) is likely the result of strong niche filtering, corroborating previous conclusions that habitat filters determine species (and therefore trait) composition in freshwater communities (Grossman et al., 1998; Peres-Neto, 2004). The low observed functional richness in the Devonian freshwater community is consistent with this conclusion. However, Canowindra’s high functional richness after controlling for species diversity (Fig. 1b) and its high nearest-neighbour scores (Fig. 1c,d) suggest that species within this community were more functionally distinct from each other than were species within the other communities examined. In other word, Canowindra was functionally diverse for a community with low species richness. The reasons for this higher-than-expected functional diversity and distinctiveness remain unclear but might reflect Devonian freshwater species using niche space that is not available to fish today — perhaps because tetrapods occupy that niche (e.g., crocodiles, turtles, beavers, and platypus) or make it unfeasible for fish (Gess and Whitfield, 2020).

Reef communities in both periods have high observed functional richness (Fig. 1a), and low-to-intermediate nearest-neighbour scores (Fig. 1c,d). This combination suggests that many, densely concentrated and diverse niches are facilitated by complex and productive reef habitats (corroborating research on modern communities) (Gratwicke and Speight, 2005). Estuarine communities have higher nearest-neighbour scores than reef communities in their climate zone and time period (Fig. 1c,d), indicating that their species are more distinct from each other in trait space. The trait variation among fish in estuarine communities might reflect that these assemblages are an admixture of species from different origins — they include marine, freshwater, and diadromous fish, as well as species that complete their life cycle in estuaries — and therefore include species that have experienced different niche/environmental filters (Passos et al., 2016; Potter et al., 2015). The higher species richness in reef compared to estuarine communities (e.g., 208 *versus* 86 species for tropical communities, and 27 *versus* 17 species for temperate/subtropical communities), combined with their relatively low observed nearest-neighbour scores, potentially explains why standardising for species richness switches which habitat type had the higher functional richness (Fig. 1 a,b). The difference might result from saturation of trait space in species-rich reef communities (i.e., the expansion of a community’s trait space associated with the addition of species diminishes as more species are added to the community), a hypothesis supported by the low nearest-neighbour scores for the reef compared to estuarine communities (Fig. 1c,d).

### 4.3 Climate

Observed functional richness of tropical communities was higher than that of temperate and subtropical communities, consistent with the previously reported pattern of increasing functional richness with decreasing latitude in fish, other vertebrates, and invertebrates (Berke et al., 2014; Cardoso et al., 2011; Jarzyna et al., 2021; Lamanna et al., 2014; Mouillot et al., 2021, 2014; Myers et al., 2021; Pigot et al., 2016; Schumm et al., 2019; Stuart-Smith et al., 2013). This pattern was reversed when we standardised for species diversity (Fig. 1b), indicating that high trait diversity in tropical fish communities is associated with high species richness.

Our results also revealed a latitudinal/climate pattern among communities in terms of their nearest-neighbour scores, with tropical communities having lower scores than their temperate/subtropical counterparts (Fig. 1c,d). Together, functional richness and nearest-neighbour scores indicate that fish communities in the tropics have greater overall variation in traits but their constituent species are packed more closely together in trait space, compared to temperate and subtropical communities (where species richness also tends to be lower) (Hillebrand, 2004; Stuart-Smith et al., 2013). This suggests stronger niche differentiation and reduced functional redundancy at higher latitudes, potentially indicating that competition is more limiting in temperate than tropical zones (Ford and Roberts, 2018). We were unable to assess whether latitudinal patterns occurred within habitat types among the Devonian fish communities because we could not include replicate communities from each habitat type for this period. Thus, future research could test whether latitude (climate) has been important in determining community trait space within habitat types through time, or whether this pattern developed recently.

### 4.4 Limitations

Although we detected patterns in trait space associated with time period, habitat, and climate, there are several important limitations to acknowledge. First, we included three Devonian and six modern communities, providing a snapshot of trait diversity present in both periods. Including more communities — particularly more Late Devonian communities — would improve the generality of our conclusions. Second, comparisons between ancient and modern communities are complicated by taphonomic biases. To address this issue, we (*i*) restricted Devonian communities to those in which most or all preserved fish species were estimated to have been discovered (based on discovery curves or the Chao1 estimator, although the Canowindra fish beds represent only a narrow temporal snapshot, raising uncertainty about whether the full range of fish diversity was captured), and (*ii*) controlled for species richness using standardised effect sizes for functional richness and functional nearest-neighbour scores. Nevertheless, taphonomic biases could still influence trait-space patterns. Future trait-space research could attempt to correct for such biases — for example, by accounting for preservation potential (Mitchell, 2015). Third, intraspecific variation influences trait space (Moran et al., 2016), but it was prohibitively difficult to include in our study due to the large number of species and the challenge of quantifying intraspecific variation in extinct species, some of which are known only from a single specimen. However, it might be possible to infer intraspecific variation and incorporate it in palaeo *versus* modern trait-space comparisons as modelling techniques advance. Finally, the temporal and spatial sampling of each community varied. For example, Canowindra represents a single, rapid drying event whereas the Miguasha fossil record spans 1.6 to 2.5 million years (Cloutier et al., 2011) — sampling variation that could affect trait space results. Future studies that include more communities could test and correct for association between trait space and spatial or temporal scope.

### 4.5 Conclusion

The differences we detected between modern and Late Devonian fish communities suggest that community trait space has changed through time. Although the Devonian period is known for its diverse and abundant fish fauna (especially the Late Devonian) (Friedman and Sallan, 2012), our results imply that modern fish communities are more trait-diverse than their Late Devonian counterparts, but that this distinction disappears after correcting for species richness. Modern communities also have greater functional redundancy, with fish closer in functional traits space to their nearest neighbour, and the trait spaces of Late Devonian and modern communities are centred in different locations in trait space. These differences could indicate several phenomena, including the longer time modern communities have had to develop and diversify (Friedman and Sallan, 2012), greater variation in modern habitats (Benton and Emerson, 2007; Villéger et al., 2011), phylogenetic constraints on the functional traits in different fish lineages (McKitrick, 1993), and/or stochasticity in evolutionary pathways (Champagnat et al., 2006). Irrespective of the root cause(s), the observed differences in trait space suggest Late Devonian communities were structured and functioned differently to modern communities. Further research on how community structure and function has changed through time, combining information from ancient and modern communities (Fritz et al., 2013), could provide important insights into community-assembly rules and the ecological and evolutionary responses of communities to environmental disturbances in the past and the future.

## Acknowledgements

We thank S. Libralato, R. Angelini, A. Pérez-Matus, S. Villéger, C. Brauer for help accessing species lists and trait data for modern fish communities. We thank T. Llewelyn for providing helpful feedback. We also acknowledge the traditional owners of the lands on which this research was done, including the Gooniyandi, Kaurna, Wiradjuri, and Mi’kmaq peoples. Funding: This work was supported by the Australian Research Council [DP220100825].

## Data Availability

All data and code are available on Github (github.com/JohnLlewelyn/Trait-space-disparity).

## Declaration of Generative AI and AI-assisted technologies in the writing process

During the preparation of this work the authors used ChatGPT to improve the readability of some sections of text. After using this tool, the authors reviewed and edited the content as needed and take full responsibility for the content of the publication.

## Notes

https://github.com/JohnLlewelyn/Trait-space-disparity

### Competing Interest Statement

The authors have declared no competing interest.

### Summary of Updates

We have included an additional Devonian community, updated some text, and rerun analyses with the additional community.

