## Supplementary for "Trait-space disparity in fish communities spanning 380 million years from the Late Devonian to present"

| **Figures** | |
|---|---|
| Fig. S1 | Map showing geographic location of Devonian and modern communities |
| Fig. S2 | Discovery curve for the Gogo Fm |
| Fig. S3 | Diagram of fish traits used |
| Fig. S4 | Function diversity metrics calculated using a subset of traits for which multiple imputation was highly accurate |
| Fig. S5 | Function diversity metrics calculated using morphometric traits only |
| Fig. S6 | Additional functional diversity metrics calculated using all 11 traits |
| Fig. S7 | Dissimilarity in Jaccard index due to turnover |
| Fig. S8 | Dissimilarity in Jaccard index due to nestedness |
| Fig. S9 | Pair plots of each community's trait space *versus* the global (all nine communities') trait space. |
| **Tables** | |
| Table S1 | Devonian species traits |
| Table S2 | Gogo Fm discovery curve data |
| Table S3 | Modern community details |
| Table S4 | Observed and inferred traits (all study species) |
| Table S5 | Community centroid positions on axes |
| Table S6 | Trait-axes correlations |

All data and code are available on GitHub (github.com/JohnLlewelyn/Trait-space-disparity).

**Figure S1.** Map showing the present-day geographic locations of the Devonian fossil sites and modern communities used in this study.

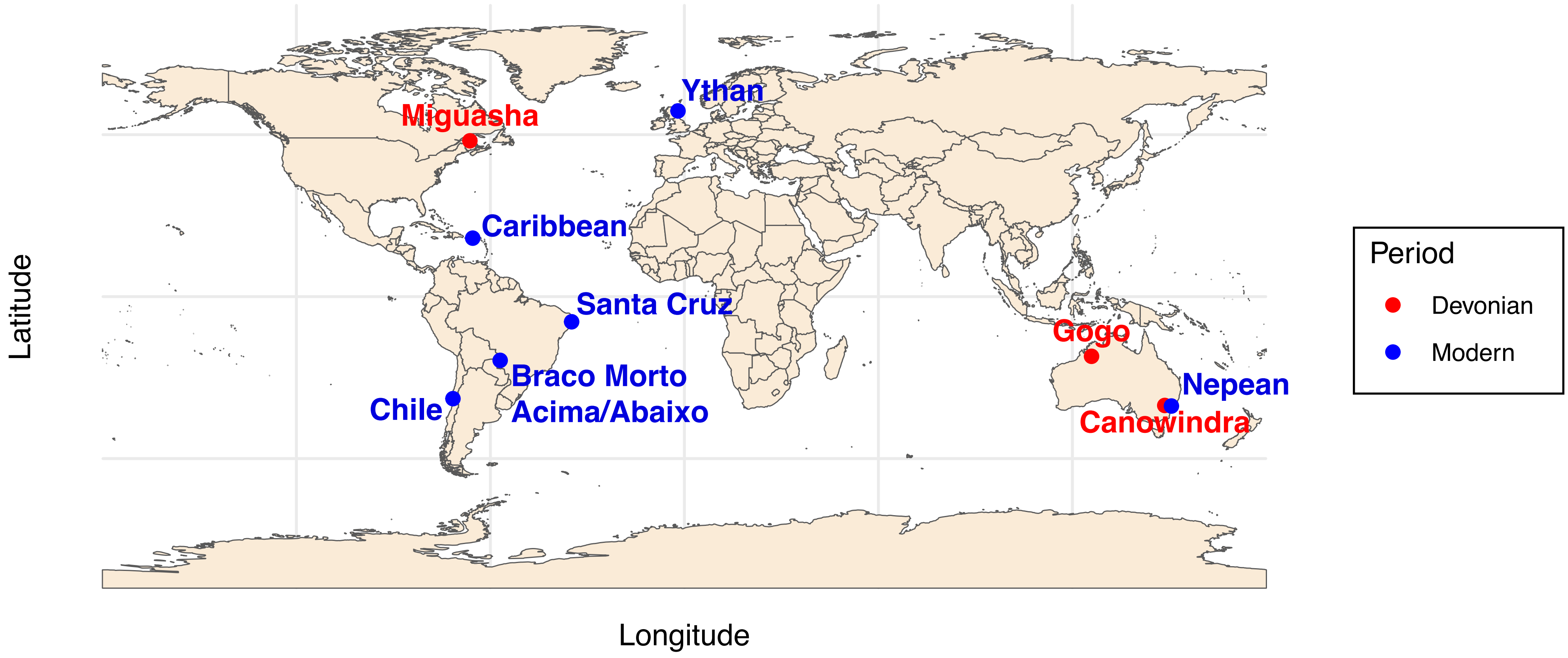

**Figure S2.** Discovery curve for the Gogo Formation fish assemblage. Blue dots indicate years when new species were named/described (x axis) and the cumulative number of described species (y axis). Solid red line shows an asymptotic regression model fit to the species description data (i.e., blue dot data), and dashed red line indicates the model's asymptote — an estimate of species diversity. The model estimates that 54 species inhabited the reefs adjacent to Gogo. We included 53 species for which we had trait data in our analyses.

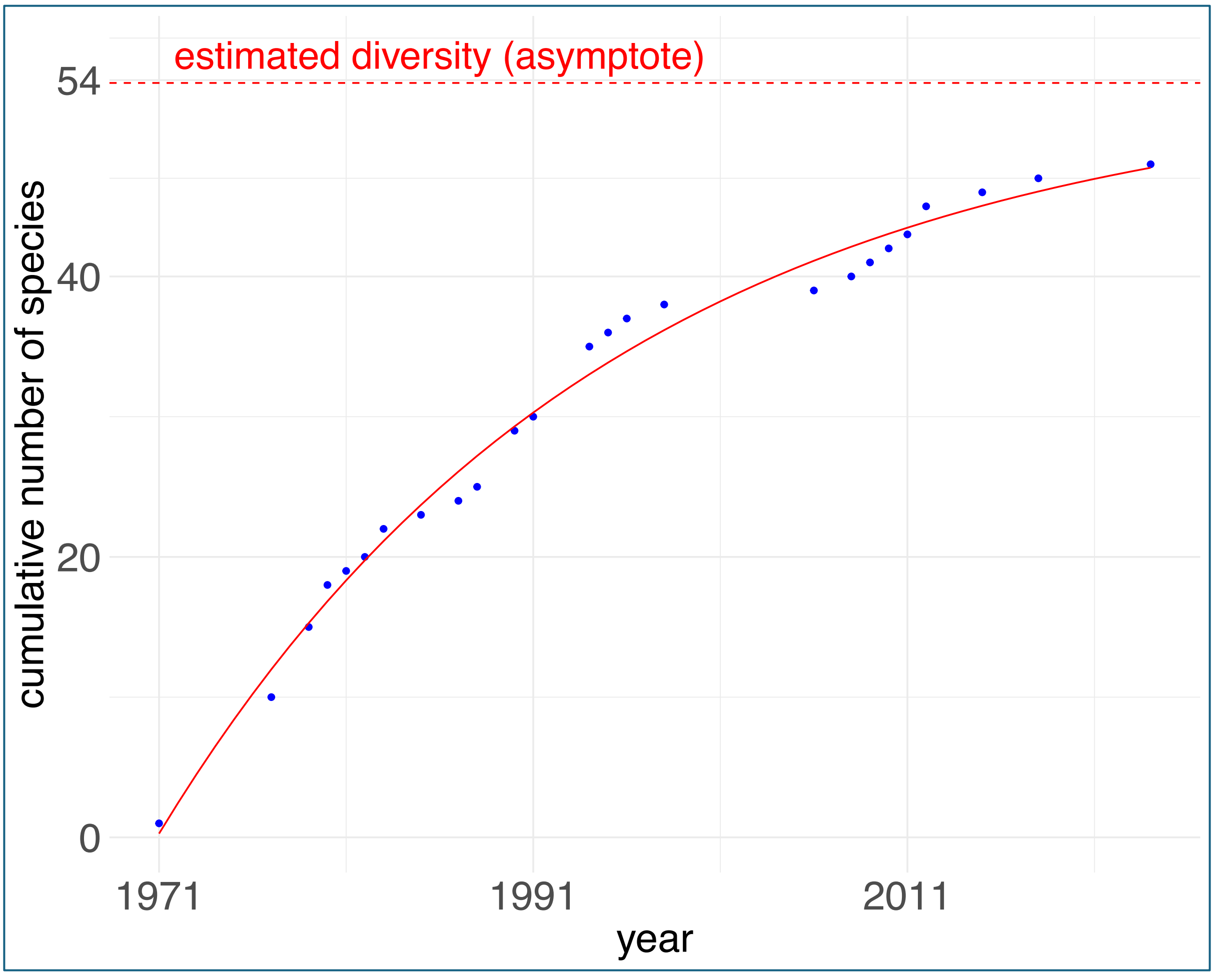

**Figure S3.** Fish traits used in the trait space analysis. An example from each category is shown for the categorical traits, whereas trait extremes are shown for continuous traits. Sagittal body shapes include (a) short and deep, (b) elongated, (c) eel-like, (d) fusiform, and (e & f) other. Transverse body shapes include (g) circular, (h) oval, (i) compressed, (j) flattened, and (k) compressed and lies on side. Eye positions included (l & m) lateral, (n) raised/top of head, and (o) migrated to one side and raised. Large spiracles could be present (p) or absent. Caudal fin shapes included (q) heterocercal, (r) homocercal, (s) hypocercal, (t) isocercal, (u) protocercal, (v) leptocercal, (w) reduced or absent, and (x) whip-like. Mouth positions included (y) terminal, (z) superior, and (aa) subterminal/inferior.

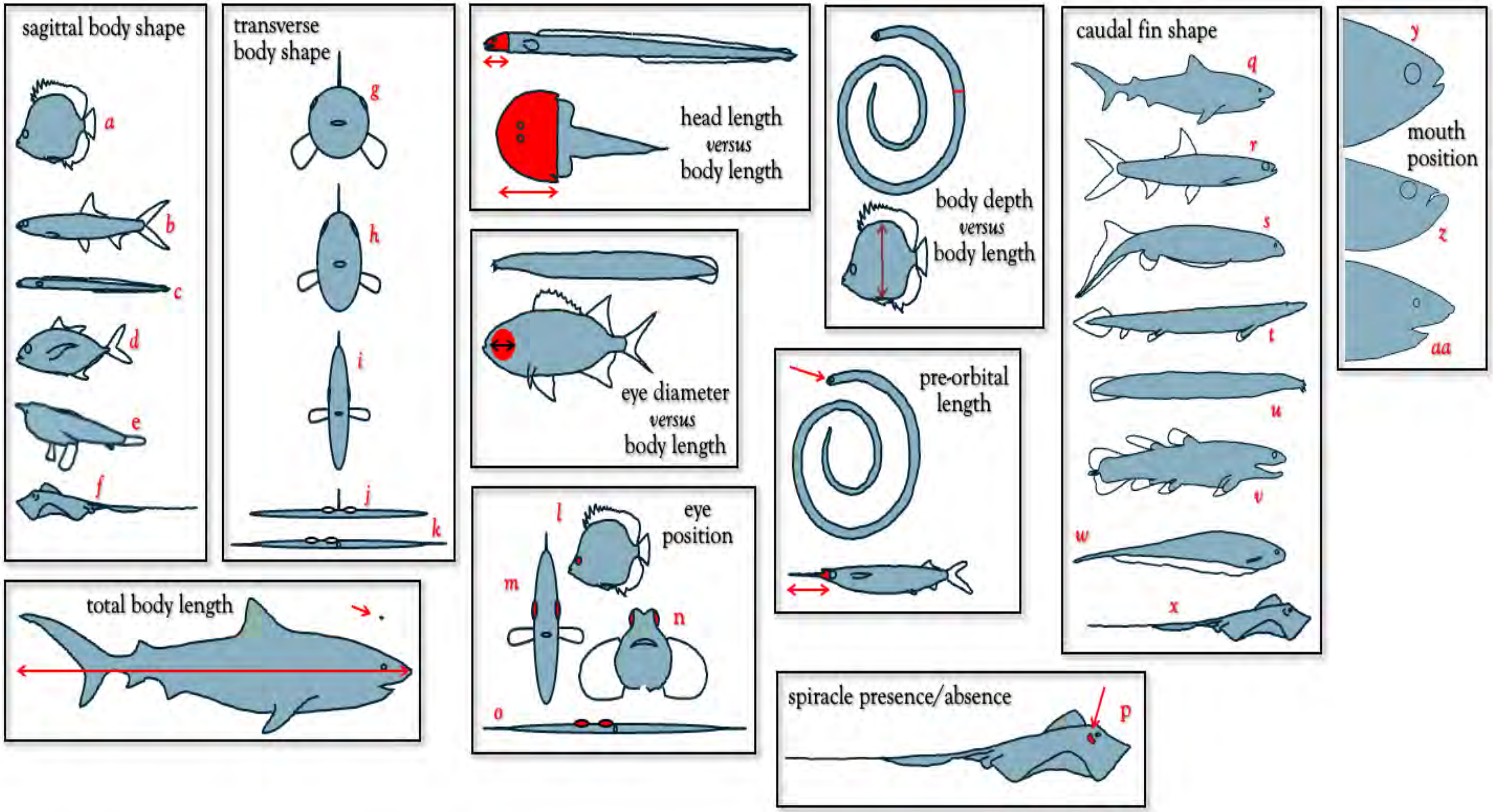

**Figure S4.** Five functional diversity metrics calculated using the subset of traits for which multiple imputation was highly accurate (numeric traits with a normalised root mean-squared error < 0.3, and categorical traits with piecewise constant fitting < 0.25).

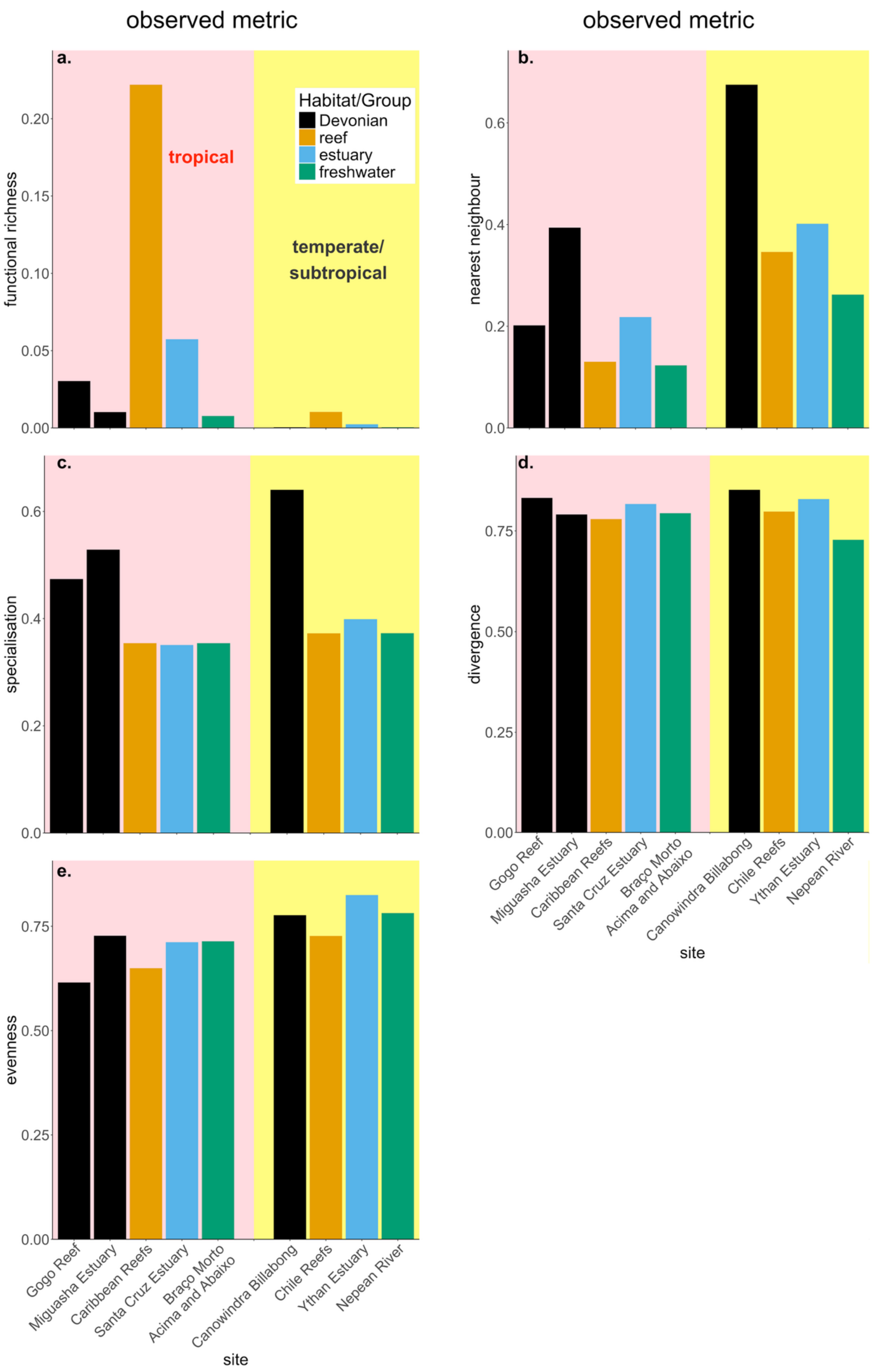

**Figure S5.** Five functional diversity metrics calculated using morphometric traits only.

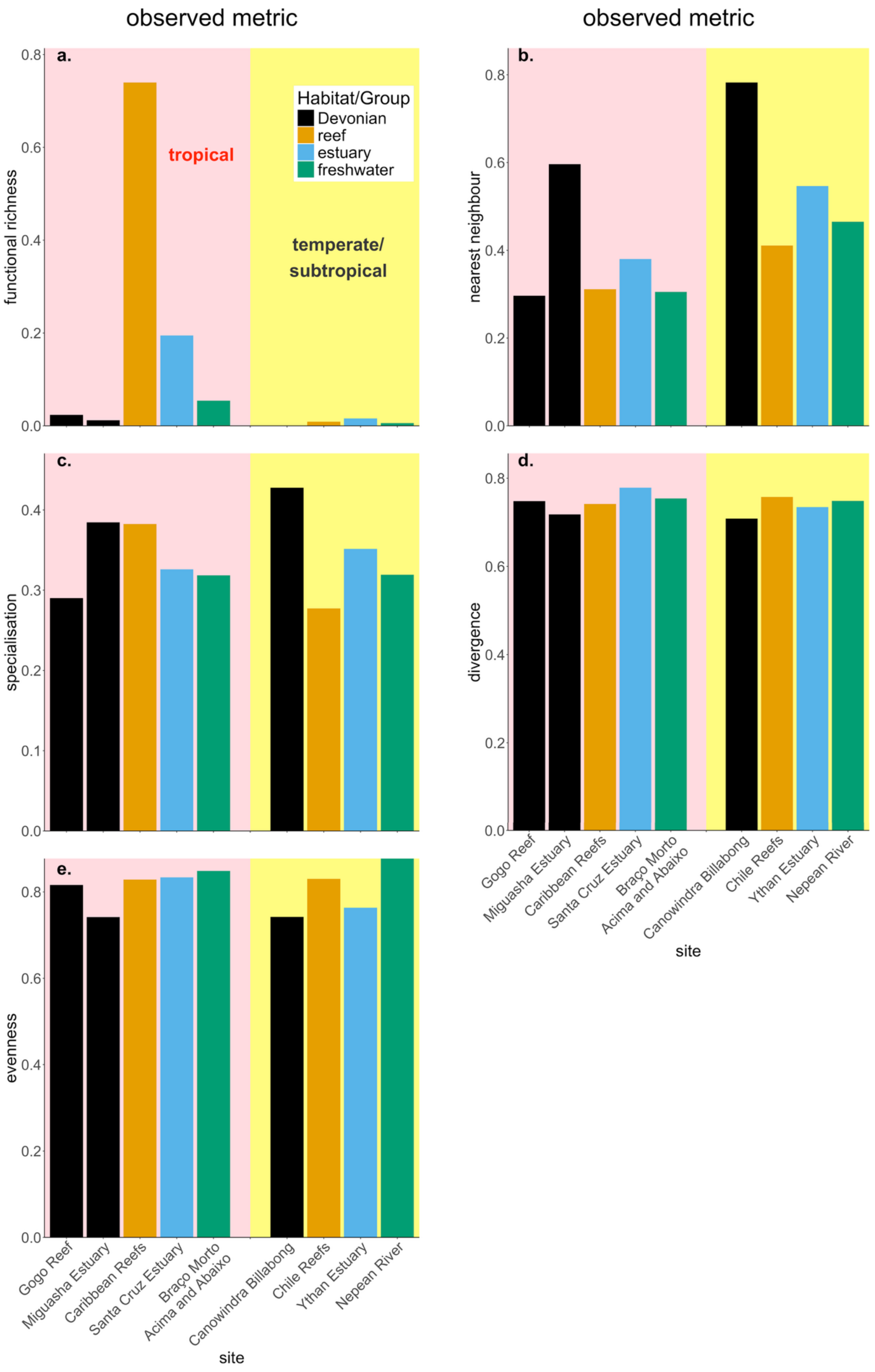

**Figure S6.** Additional functional diversity metrics calculated using all 11 traits. The metrics include functional evenness, functional divergence, and functional specialization (left column shows observed scores; right column shows standardised effect sizes).

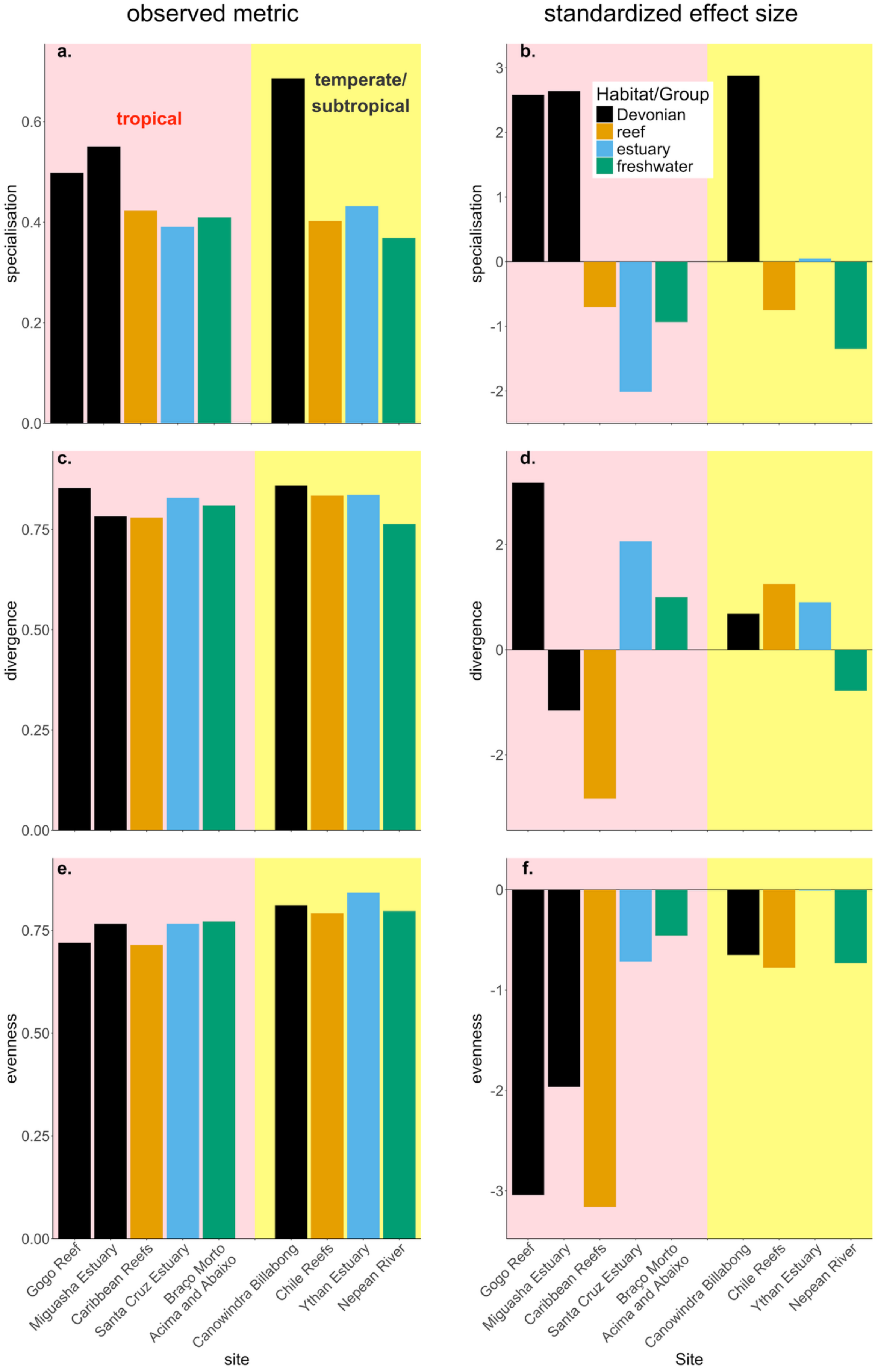

**Figure S7.** Component of Jaccard index dissimilarity due to turnover.

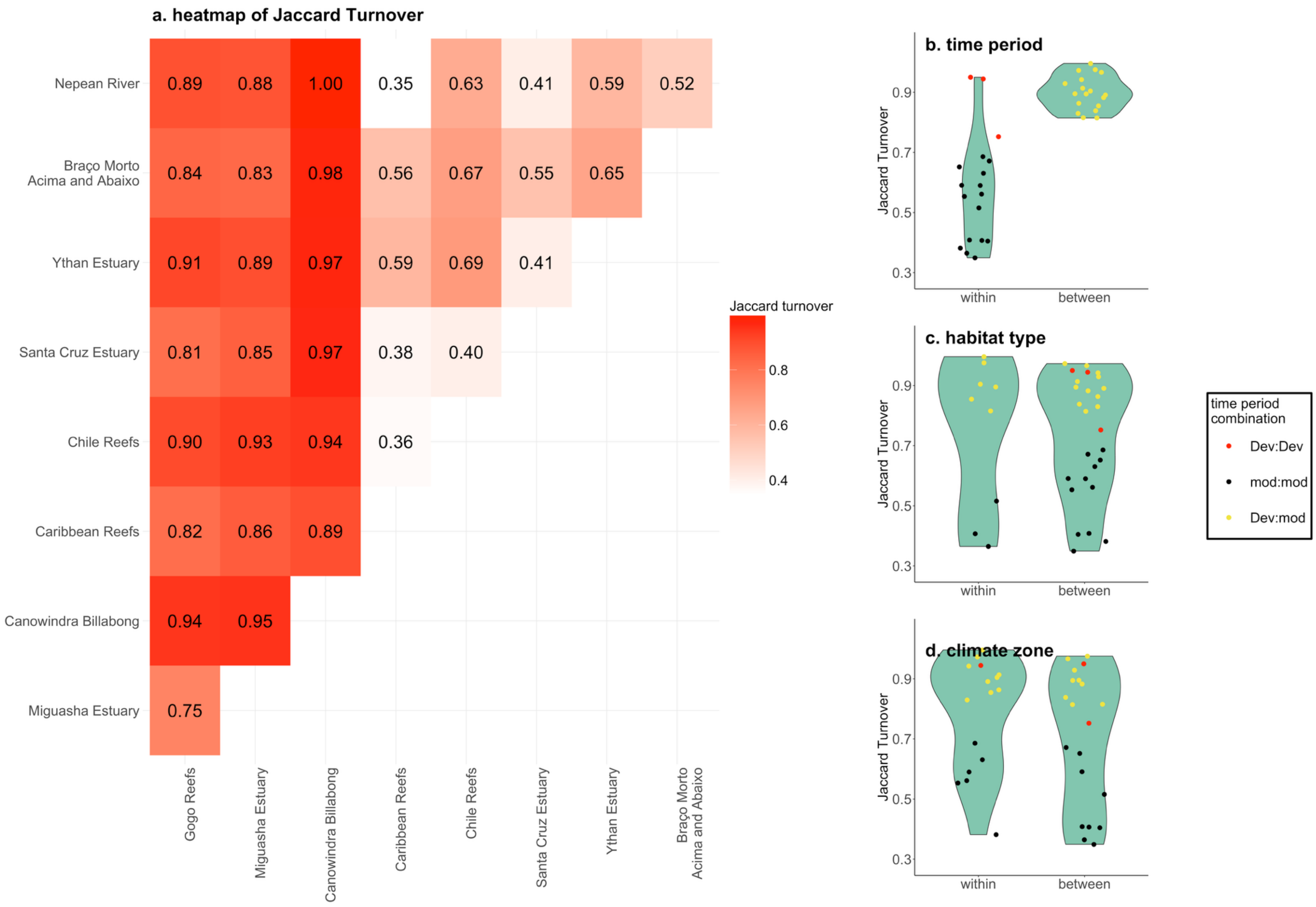

**Figure S8.** Component of Jaccard index dissimilarity due to nestedness.

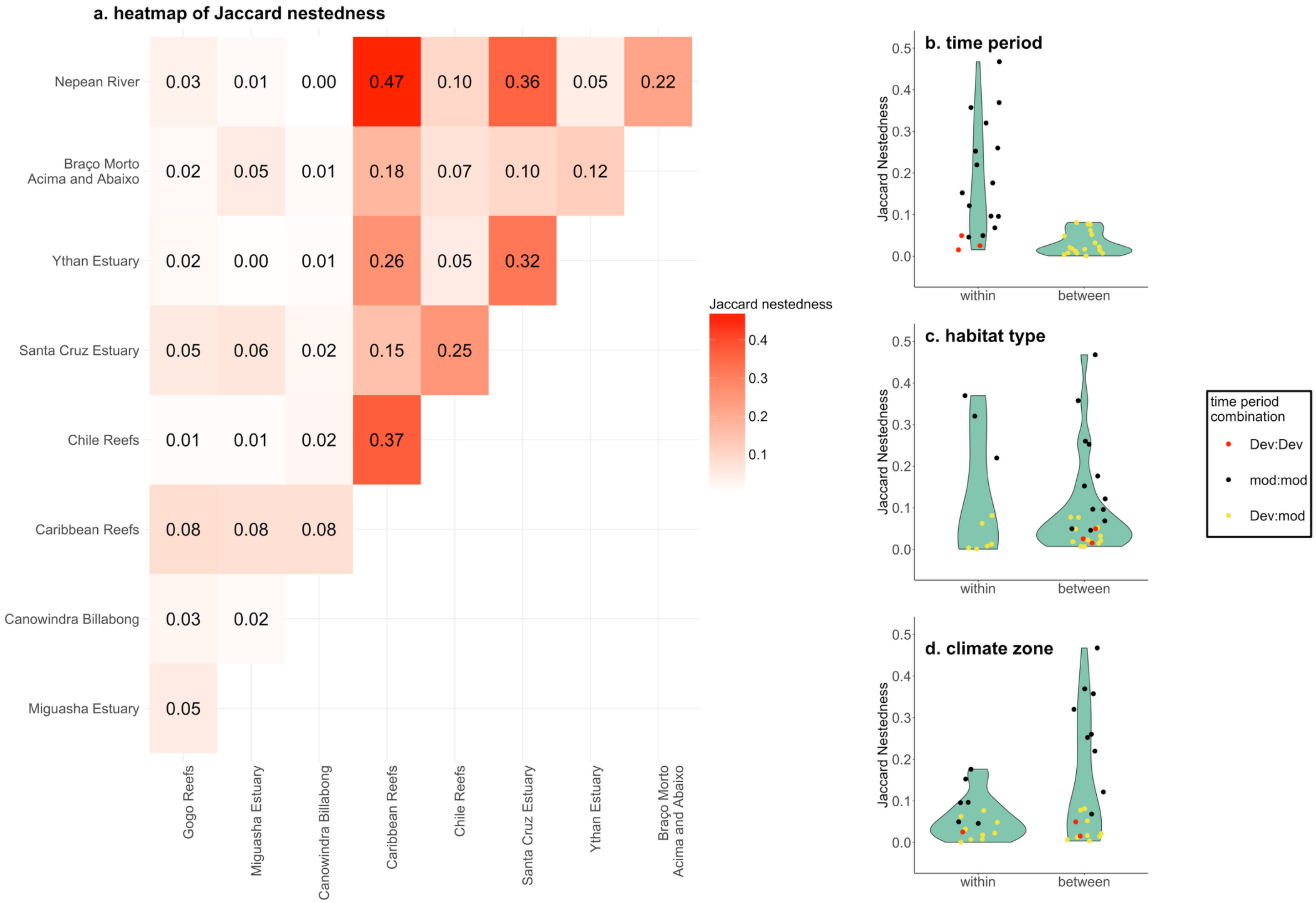

**Figure S9.** Pair plots comparing each community's trait space to the global (i.e., nine communities we examined in this study) trait space across seven axes. Trait spaces are shown by plotting 400,000 random points in the two hypervolumes being compared. The two large points with white margins in each plot indicate the centroids of the focal (red) and global (black) trait spaces.

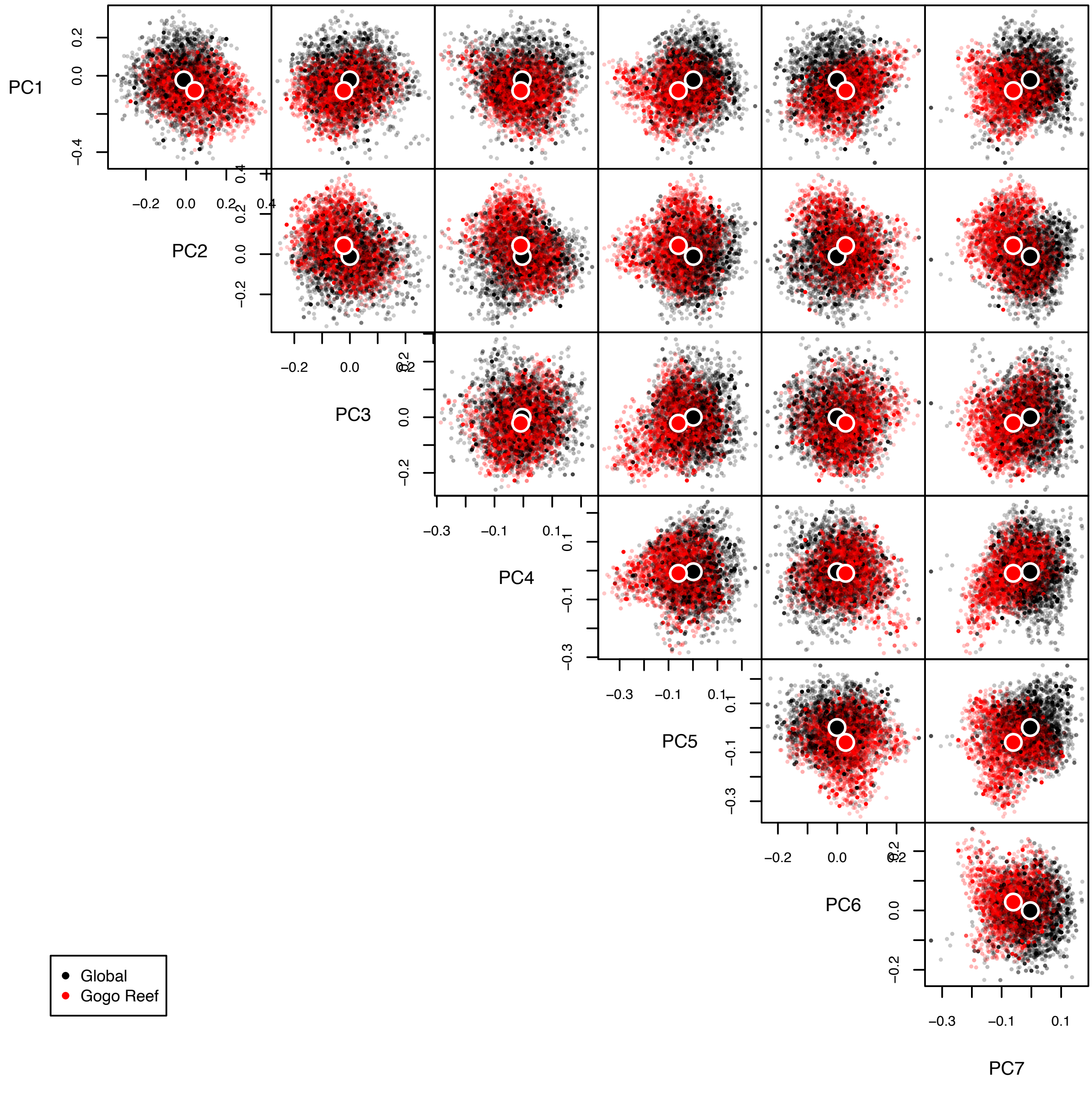

PC1
PC2
PC3
PC4
PC5
PC6
PC7
Global
Gogo Reef

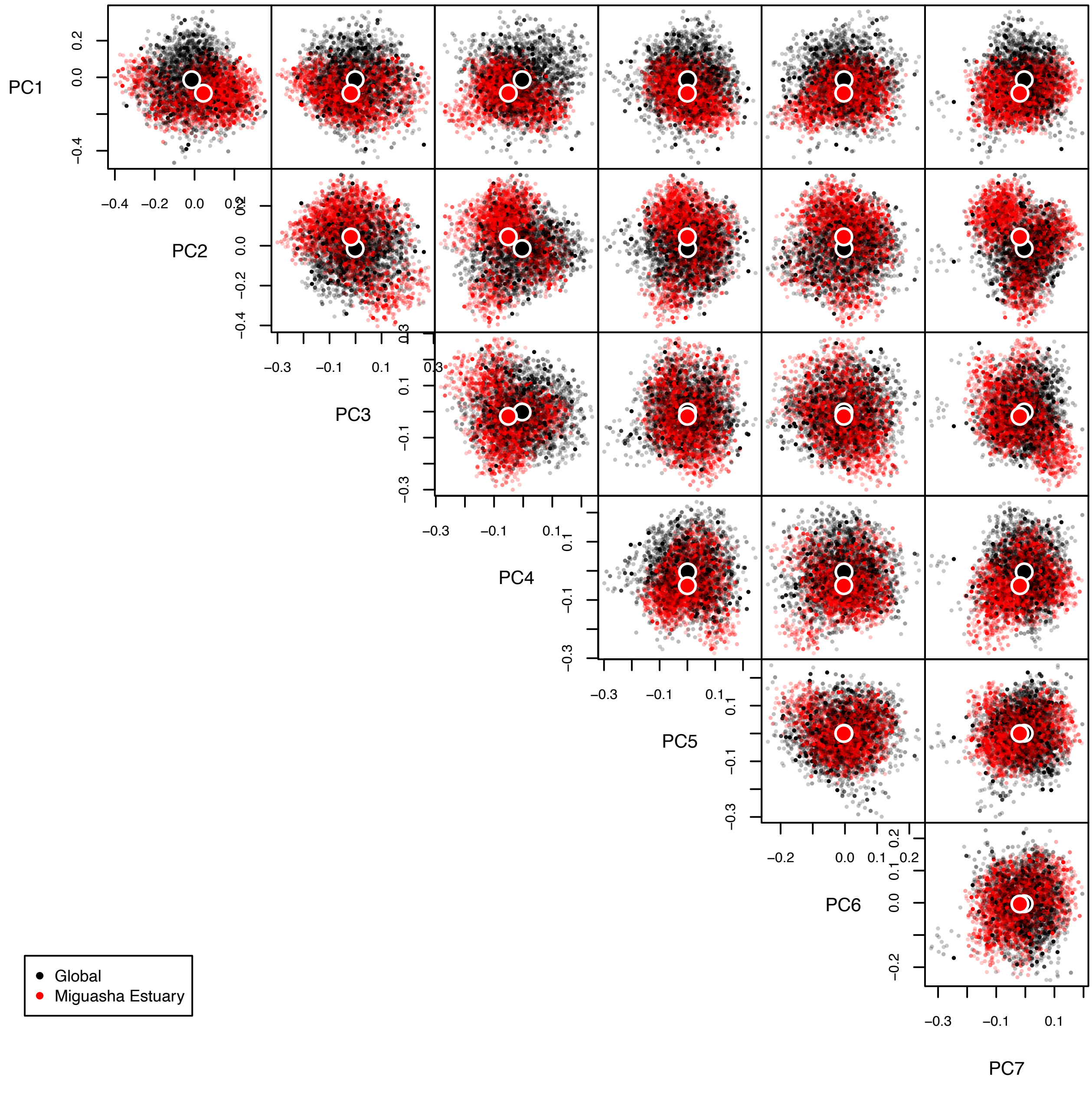
PC1
PC2
PC3
PC4
PC5
PC6
PC7
Global
Miguasha Estuary

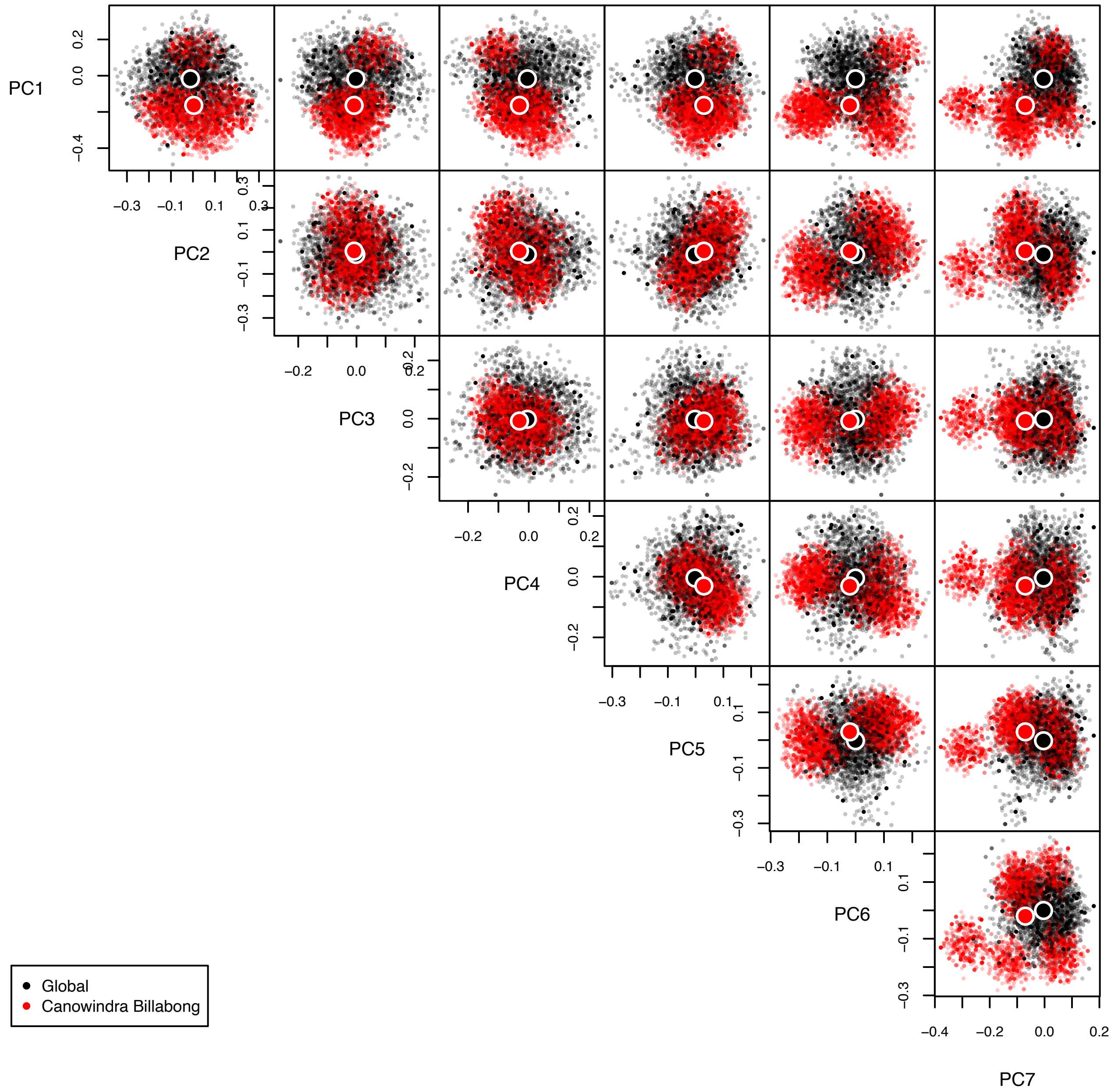
PC1
PC2
PC3
PC4
PC5
PC6
PC7
Global
Canowindra Billabong

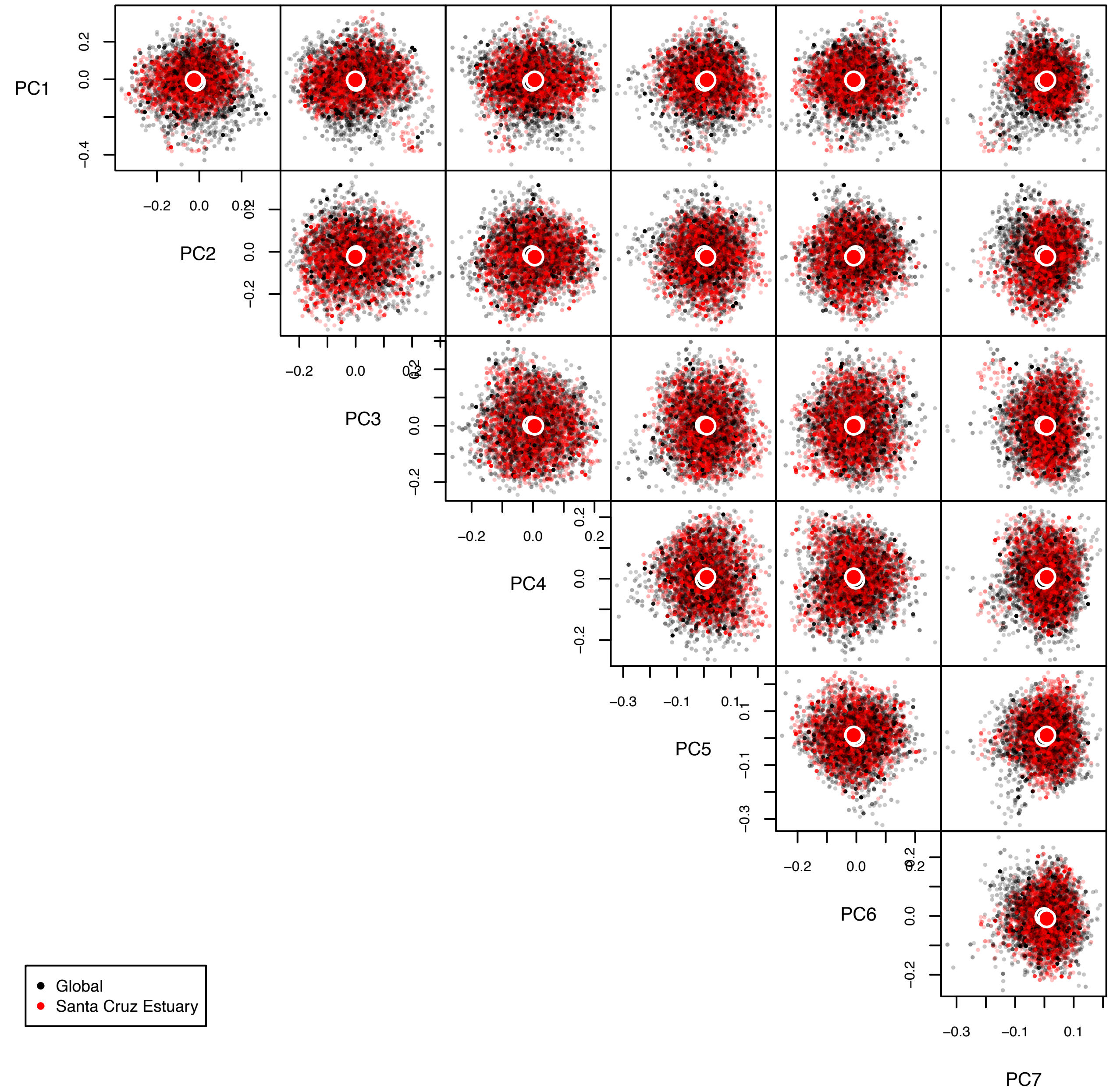
PC1
PC2
PC3
PC4
PC5
PC6
PC7
Global
Santa Cruz Estuary

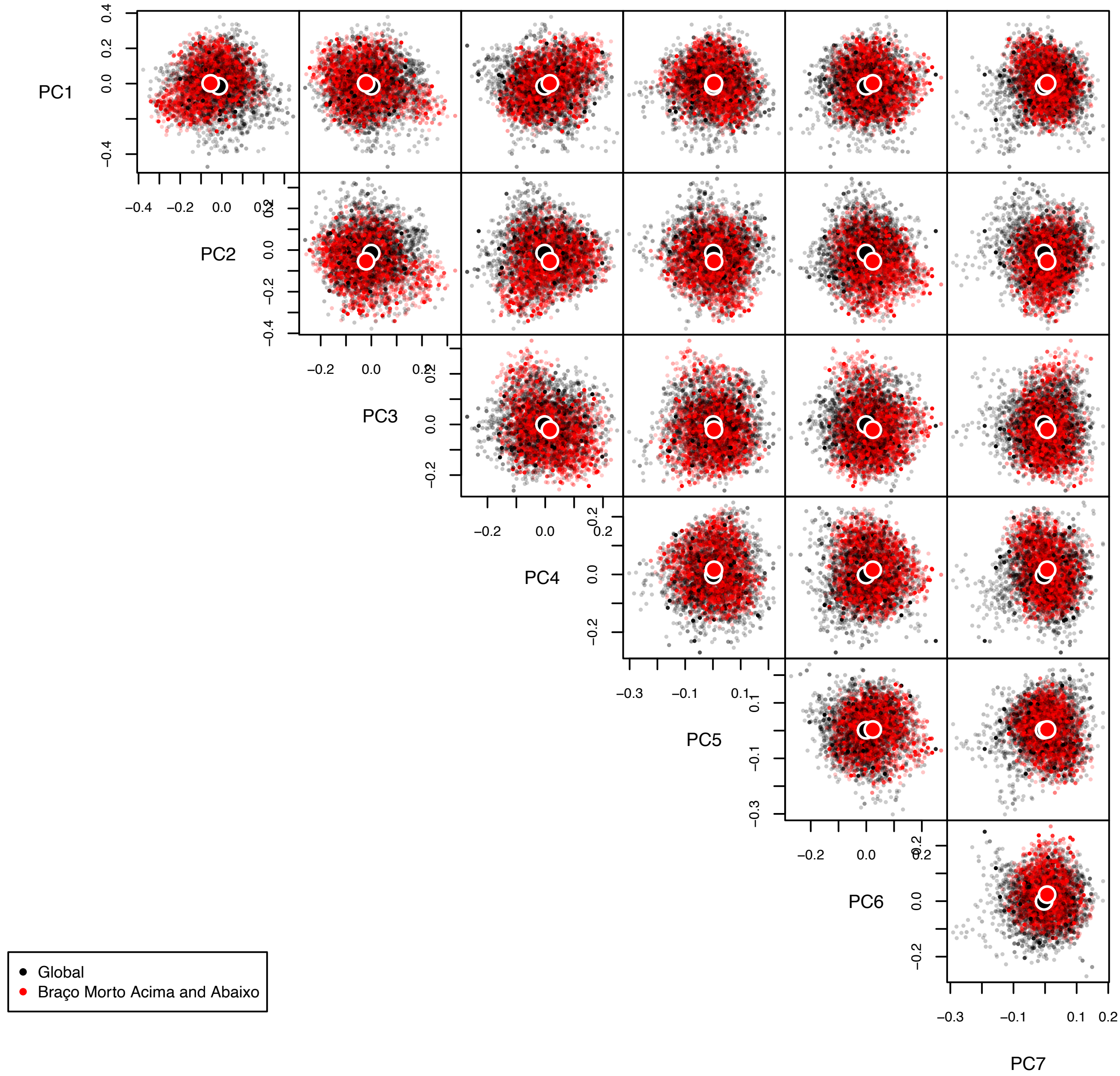
PC1
PC2
PC3
PC4
PC5
PC6
PC7
Global
Braço Morto Acima and Abaixo

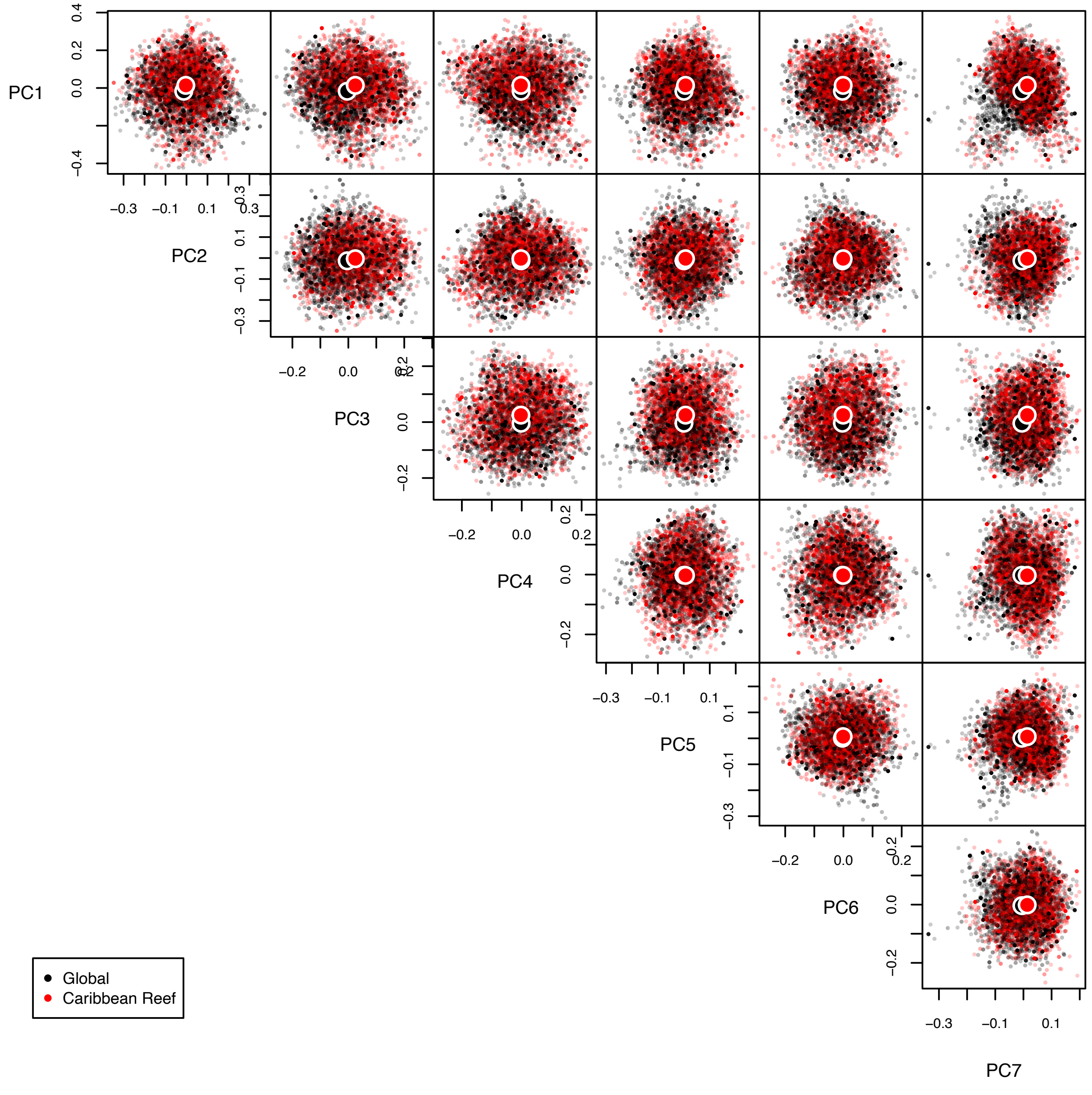
PC1
PC2
PC3
PC4
PC5
PC6
PC7
Global
Caribbean Reef

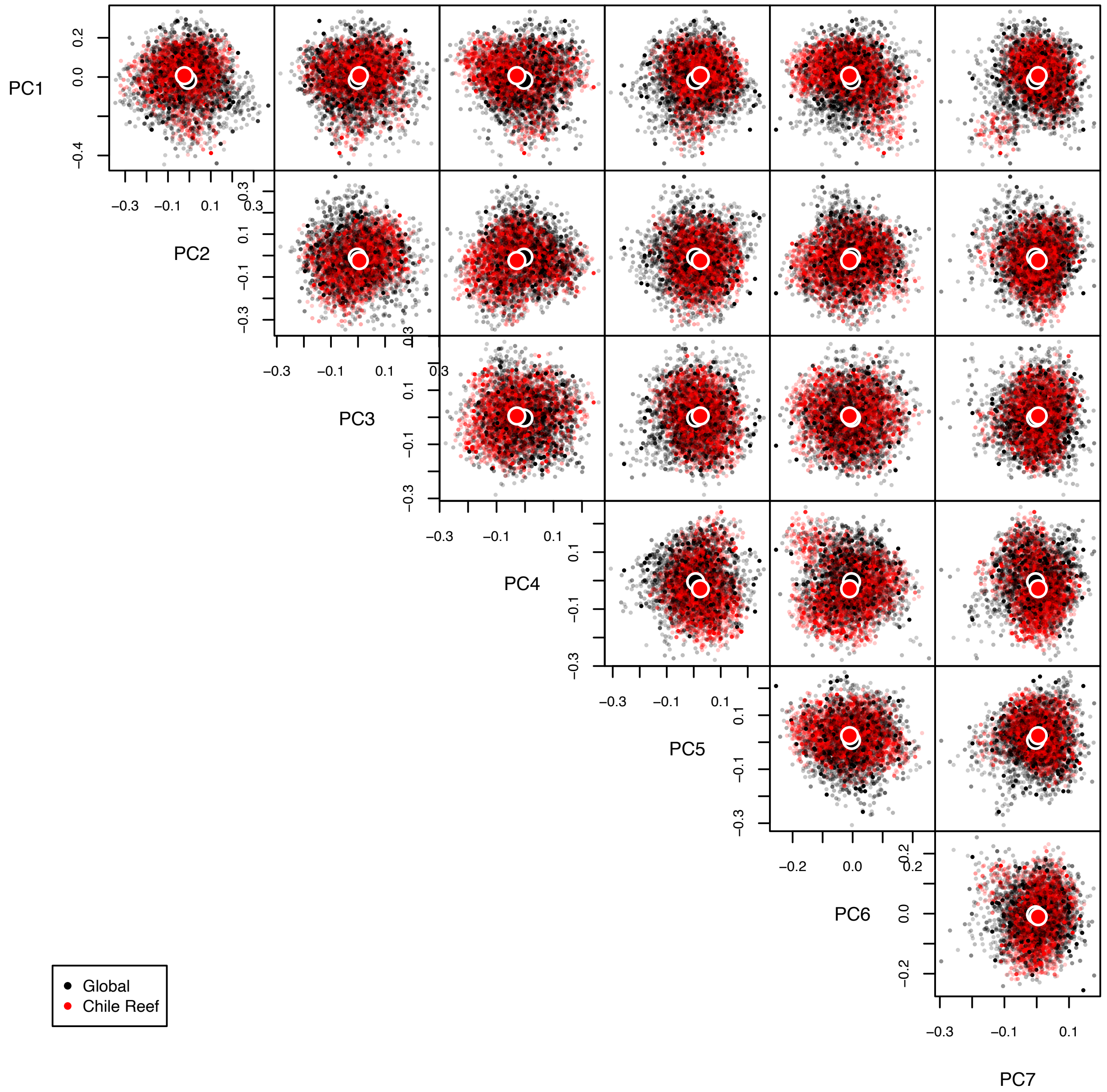
PC1
PC2
PC3
PC4
PC5
PC6
PC7
Global
Chile Reef

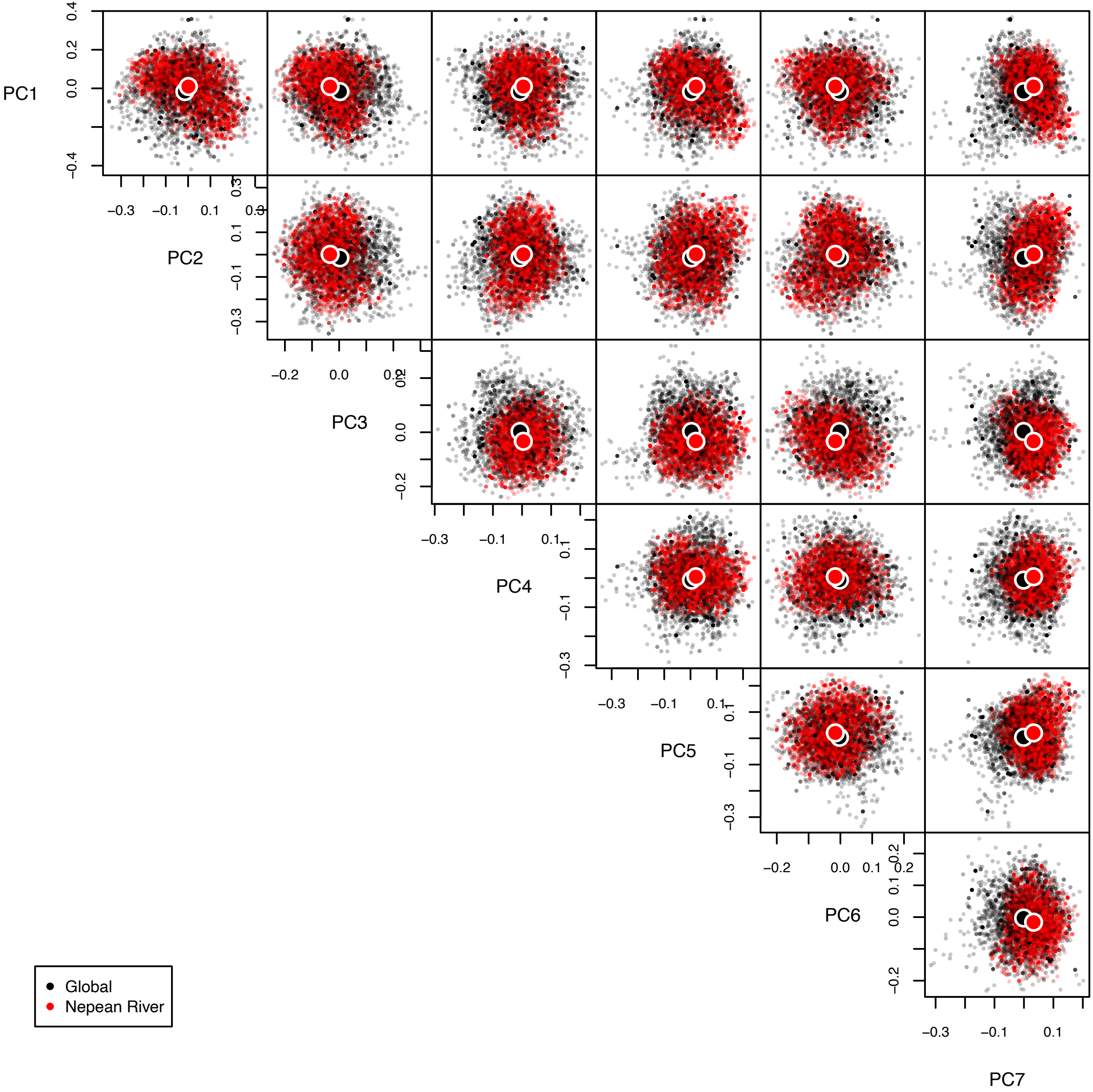
PC1
PC2
PC3
PC4
PC5
PC6
PC7
Global
Nepean River

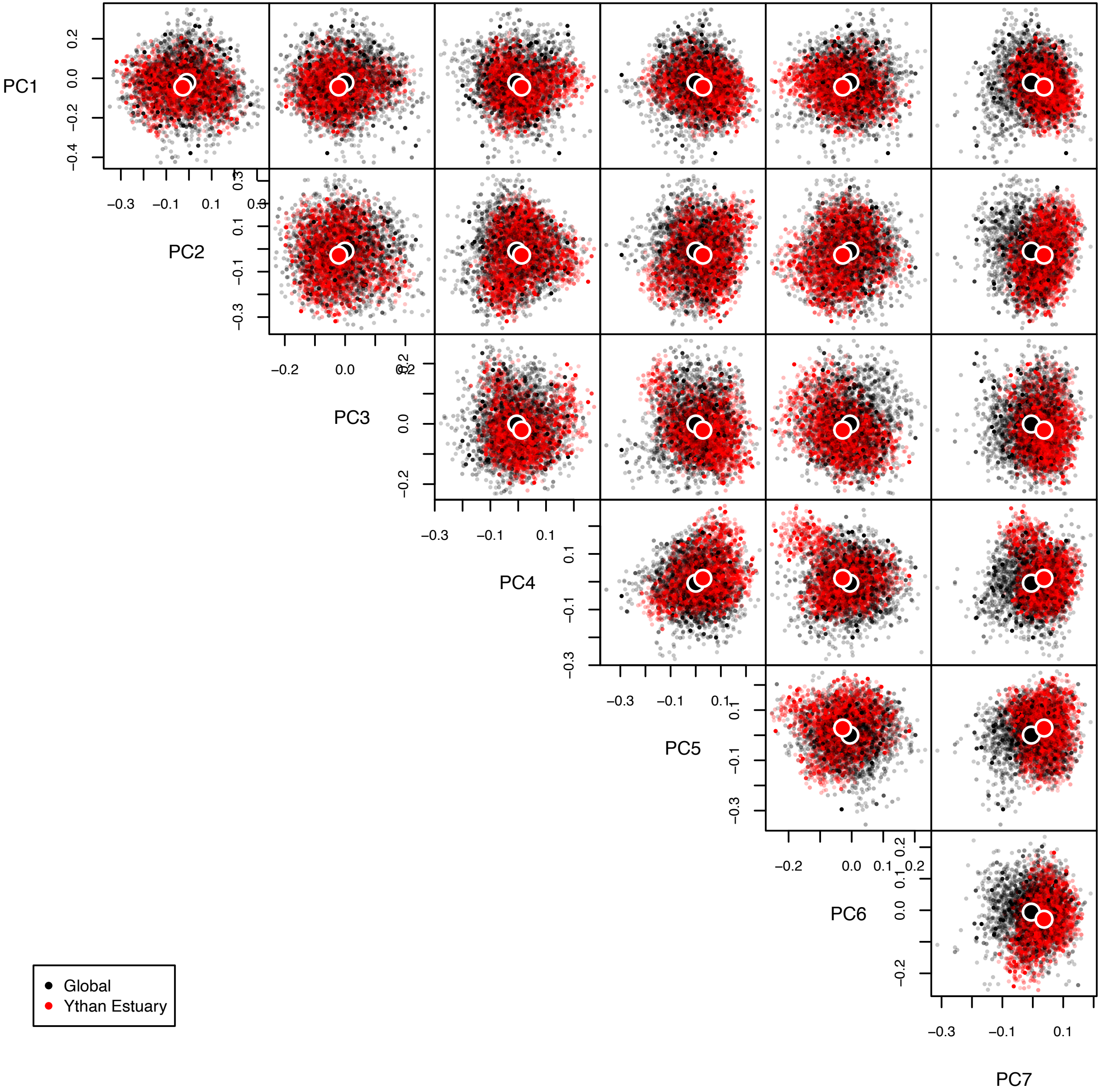
PC1
PC2
PC3
PC4
PC5
PC6
PC7
Global
Ythan Estuary

**Table S1.** Traits of fish from three Late Devonian communities: Gogo, Miguasha, and Canowindra (not including traits inferred using multiple imputation).

| taxon | site | total length | standard length | head length | pre-orbital length (snout length) | body depth (body height) | eye size (diameter) | body shape I (saggital plane) | body shape II (transverse plane) | mandible | mouth position | eye position | spiracular | caudal fin shape |
|---|---|---|---|---|---|---|---|---|---|---|---|---|---|---|
| *Adololopas moyasmithae* | Gogo | NA | NA | 72.8 | 19.1 | NA | 8.8 | NA | NA | present | sub-terminal/inferior | lateral | absent | NA |
| *Asthenorhynchus meemannae* | Gogo | NA | NA | 130 | 45 | NA | 15 | NA | NA | present | sub-terminal/inferior | lateral | absent | NA |
| *Austroptyctodus gardineri* | Gogo | 200 | 200 | 20 | NA | NA | NA | elongated | compressed | present | terminal | lateral | present | whip-like |
| *Bothriolepis canadensis* | Miguasha | 912 | 912 | 112 | 28 | 146 | 23 | short and/or deep | angular | present | sub-terminal/inferior | front-facing | absent | heterocercal |
| *Bothriolepis sp.* | Gogo | 330 | NA | 40 | 20 | 50 | 9 | short and/or deep | angular | present | sub-terminal/inferior | raised/top of head | NA | heterocercal |
| *Bothriolepis yeungae* | Canowindra | 500 | NA | 50 | 20 | 45 | 6 | fusiform/normal | angular | present | sub-terminal/inferior | raised/top of head | NA | heterocercal |
| *Bruntonichthys multidens* | Gogo | 440 | 440 | 80 | 10 | NA | 22 | NA | NA | present | terminal | lateral | NA | heterocercal |
| *Bullerichthys fascidens* | Gogo | 493 | 493 | NA | NA | NA | 15 | NA | NA | present | terminal | lateral | NA | heterocercal |
| *Cabonnichthys burnsi* | Canowindra | 750 | 646 | 123 | 27.4 | 92.3 | 11 | elongated | circular | present | terminal | lateral | NA | isocercal |
| *Callistiopterus clappi* | Miguasha | 78 | 76 | 19 | 5 | 11 | 2 | fusiform/normal | circular | present | terminal | lateral | present | isocercal |
| *Campbellodus decipiens* | Gogo | 500 | 500 | NA | NA | NA | NA | elongated | compressed | present | terminal | lateral | NA | whip-like |
| *Camuropiscis concinnus* | Gogo | 290 | 290 | 62 | 17 | 36.5 | 12.5 | fusiform/normal | circular | present | sub-terminal/inferior | lateral | absent | heterocercal |
| *Camuropiscis laidlawi* | Gogo | 220 | 220 | 55 | 25 | NA | 9 | NA | NA | present | sub-terminal/inferior | lateral | NA | heterocercal |
| *Canowindra grossi* | Canowindra | 500 | 410 | 106 | 35 | 45 | 4.5 | elongated | circular | present | terminal | lateral | NA | NA |
| *Cheirolepis canadensis* | Miguasha | 530 | 499 | 127 | 15 | 96 | 15 | fusiform/normal | oval | present | terminal | lateral | present | heterocercal |
| *Chirodipterus australis* | Gogo | 286.5 | 286.5 | 59.3 | 18.2 | 59.3 | 8 | short and/or deep | circular | present | sub-terminal/inferior | lateral | absent | heterocercal |
| *Compagopiscis croucheri* | Gogo | 300 | 300 | 64.3 | 6 | 61.3 | 16.4 | fusiform/normal | circular | present | sub-terminal/inferior | lateral | NA | heterocercal |
| *Diplacanthus ellsi* | Miguasha | 130 | 130 | 22 | 2 | 22 | 4 | fusiform/normal | oval | present | terminal | lateral | absent | heterocercal |
| *Diplacanthus horridus* | Miguasha | 203 | 203 | 45 | 6 | 46 | 11 | fusiform/normal | oval | present | terminal | lateral | absent | heterocercal |

| | | | | | | | | | | | | | | |
|---|---|---|---|---|---|---|---|---|---|---|---|---|---|---|
| *Eastmanosteus calliaspis* | Gogo | 750 | 750 | 133.7 | 22 | 148.2 | 33.4 | fusiform/normal | angular | present | terminal | lateral | present | heterocercal |
| *Eastmanosteus sp. A* | Gogo | 800 | 800 | 140 | 24 | 175 | 23 | fusiform/normal | angular | present | terminal | lateral | present | heterocercal |
| *Eastmanosteus sp. B* | Gogo | 1400 | 1400 | 272 | 28 | NA | 30 | fusiform/normal | angular | present | terminal | lateral | present | heterocercal |
| *Elpistostege watsoni* | Miguasha | 1570 | 1486 | 341 | 130 | 128 | 19 | eel-like | flattened | present | terminal | raised/top of head | present | isocercal |
| *Escuminaspis laticeps* | Miguasha | 435 | 420 | 170 | 60 | 54 | 15 | other | flattened | absent | sub-terminal/inferior | raised/top of head | absent | heterocercal |
| *Euphanerops longaevus* | Miguasha | 381 | 360 | 64 | 15 | 55 | 16 | eel-like | circular | absent | sub-terminal/inferior | lateral | absent | hypocercal |
| *Eusthenopteron foordi* | Miguasha | 1180 | 1136 | 256 | 43 | 210 | 21 | fusiform/normal | circular | present | terminal | lateral | present | isocercal |
| *Fallacosteus turnerae* | Gogo | 300 | 300 | 67.4 | 22 | 38.5 | 16 | fusiform/normal | circular | present | sub-terminal/inferior | lateral | absent | heterocercal |
| *Fleurantia denticulata* | Miguasha | 591 | 551 | 156 | 66 | 129 | 27 | fusiform/normal | compressed | present | terminal | lateral | present | heterocercal |
| *Glyptolepis sp.* | Miguasha | 586 | 577 | 120 | 16 | 105 | 7 | fusiform/normal | compressed | present | terminal | lateral | present | heterocercal |
| *Gogodipterus paddyensis* | Gogo | NA | NA | 107 | NA | NA | NA | NA | circular | present | sub-terminal/inferior | lateral | absent | heterocercal |
| *Gogonasus andrewsae* | Gogo | NA | NA | 60 | 7.3 | NA | 5.3 | fusiform/normal | circular | present | sub-terminal/inferior | lateral | present | heterocercal |
| *Gogosardina coatesi* | Gogo | 135 | 135 | 30 | 4 | 23 | 9 | fusiform/normal | oval | present | sub-terminal/inferior | lateral | present | heterocercal |
| *Gogoselachus lynnbeazleyae* | Gogo | 300 | NA | NA | NA | NA | NA | elongated | circular | present | NA | lateral | present | heterocercal |
| *Gooloogongia loomesi* | Canowindra | 800 | NA | 142.7 | 45.9 | NA | 18.6 | NA | NA | present | superior | NA | NA | NA |
| *Griphognathus whitei* | Gogo | 665 | 665 | 190 | 98 | 192 | 18 | fusiform/normal | compressed | present | sub-terminal/inferior | lateral | absent | heterocercal |
| *Groenlandaspis sp.* | Canowindra | 460 | NA | 100 | 20 | 150 | 9 | short and/or deep | angular | present | sub-terminal/inferior | raised/top of head | NA | heterocercal |
| *Halimacanthodes ahlbergi* | Gogo | 240 | NA | 60 | 30 | NA | NA | elongated | compressed | present | terminal | lateral | present | heterocercal |
| *Harrytoombsia elegans* | Gogo | 280 | 280 | 60 | 13 | 72.1 | 17.6 | fusiform/normal | circular | present | terminal | lateral | present | heterocercal |
| *Holodipterus elderae* | Gogo | NA | NA | NA | NA | NA | NA | NA | NA | present | sub-terminal/inferior | lateral | absent | heterocercal |
| *Holodipterus gogoensis* | Gogo | NA | NA | 180 | NA | NA | NA | NA | NA | present | sub-terminal/inferior | lateral | absent | heterocercal |
| *Holonema westolli* | Gogo | 715 | 715 | 180 | 42 | 160 | 11 | fusiform/normal | compressed | present | sub-terminal/inferior | raised/top of head | present | heterocercal |
| *Holoptychius jarviki* | Miguasha | 470 | 455 | 94 | 13 | 83 | 10 | elongated | oval | present | terminal | lateral | present | heterocercal |

| | | | | | | | | | | | | | | |
|---|---|---|---|---|---|---|---|---|---|---|---|---|---|---|
| *Homalacanthus concinnus* | Miguasha | 309 | 309 | 46 | 4 | 63 | 9 | elongated | oval | present | terminal | lateral | absent | heterocercal |
| *Incisoscutum ritchiei* | Gogo | 290 | 290 | 57.5 | 11 | 49 | 17.3 | fusiform/normal | circular | present | sub-terminal/inferior | lateral | absent | heterocercal |
| *Incisoscutum sarahae* | Gogo | 500 | 500 | 90 | 17 | 78 | 18 | fusiform/normal | circular | present | sub-terminal/inferior | lateral | present | heterocercal |
| *Kapitany sarcopt* | Gogo | NA | NA | NA | NA | NA | NA | NA | NA | present | NA | NA | NA | NA |
| *Kendrickichthys cavernosus* | Gogo | 1040 | 1040 | 180 | 30 | NA | 30 | fusiform/normal | circular | present | terminal | lateral | present | heterocercal |
| *Kimberleyichthys bispicatus* | Gogo | NA | NA | 188 | 19 | NA | NA | fusiform/normal | circular | present | NA | lateral | NA | heterocercal |
| *Kimberleyichthys whybrowi* | Gogo | NA | NA | 125 | 11 | NA | 22.5 | fusiform/normal | circular | present | NA | lateral | NA | heterocercal |
| *Latocamurus coulthardi* | Gogo | 230 | 230 | 46 | 7 | 33 | 14.5 | fusiform/normal | circular | present | sub-terminal/inferior | lateral | absent | heterocercal |
| *Levesquaspis patteni* | Miguasha | 169 | 154 | 67 | 20 | 21 | 10 | other | flattened | absent | sub-terminal/inferior | raised/top of head | absent | heterocercal |
| *Mandageria fairfaxi* | Canowindra | 1600 | 1450 | 276 | 62.6 | 199.4 | 17 | elongated | circular | present | sub-terminal/inferior | lateral | NA | NA |
| *Materpiscis attenboroughi* | Gogo | 420 | 420 | NA | NA | NA | NA | elongated | compressed | present | terminal | lateral | NA | whip-like |
| *Mcnamaraspis kaprios* | Gogo | 354 | 354 | 56.3 | 8 | 63 | 14 | fusiform/normal | circular | present | sub-terminal/inferior | lateral | present | heterocercal |
| *Miguashaia bureaui* | Miguasha | 482 | 470 | 130 | 19 | 108 | 18 | fusiform/normal | compressed | present | terminal | lateral | present | heterocercal |
| *Mimipiscis bartrami* | Gogo | 125 | 125 | 30 | 4 | 32 | 9 | fusiform/normal | oval | present | sub-terminal/inferior | lateral | present | NA |
| *Mimipiscis toombsi (normal)* | Gogo | 80 | 80 | 25 | 2.7 | 20 | 6 | fusiform/normal | oval | present | sub-terminal/inferior | lateral | present | homocercal |
| *Mimipiscis toombsi (supersized)* | Gogo | 200 | 200 | 60 | 9 | 65 | 10 | fusiform/normal | oval | present | sub-terminal/inferior | lateral | present | homocercal |
| *Moythomasia durgaringa* | Gogo | 170 | 170 | 40 | 7 | 48 | 7 | fusiform/normal | oval | present | sub-terminal/inferior | lateral | present | heterocercal |
| *Namugawi wirngarri* | Gogo | 225 | NA | 32 | 6 | 25.6 | 5 | NA | NA | present | terminal | lateral | NA | leptocercal |
| *Onychodus jandemarrai* | Gogo | 1600 | 1581 | 320 | 50 | NA | 40 | fusiform/normal | compressed | present | terminal | lateral | present | leptocercal |
| *Pickeringius acanthophorus* | Gogo | 140 | 140 | 33 | 4.5 | 30 | 7 | fusiform/normal | oval | present | sub-terminal/inferior | lateral | present | heterocercal |
| *Pillararhynhcus longi* | Gogo | NA | NA | 60 | NA | NA | NA | NA | circular | present | sub-terminal/inferior | lateral | absent | heterocercal |
| *Pinguosteus thulborni* | Gogo | 200 | 200 | NA | NA | NA | NA | short and/or deep | circular | present | NA | lateral | NA | heterocercal |
| *Plourdosteus canadensis* | Miguasha | 966 | 966 | 187 | 34 | 172 | 22 | elongated | circular | present | sub-terminal/inferior | lateral | absent | heterocercal |
| *Quebecius quebecensis* | Miguasha | 600 | 590 | 124 | 20 | 106 | 15 | fusiform/normal | compressed | present | terminal | lateral | present | heterocercal |

| | | | | | | | | | | | | | | |
|---|---|---|---|---|---|---|---|---|---|---|---|---|---|---|
| *Remigolepis walkeri* | Canowindra | 370 | NA | 35 | 9 | 90 | 8.5 | fusiform/normal | angular | present | sub-terminal/inferior | raised/top of head | NA | heterocercal |
| *Rhinodipterus kimberleyensis* | Gogo | NA | NA | 75 | 30 | NA | 15.4 | NA | NA | present | sub-terminal/inferior | lateral | absent | heterocercal |
| *Robinsondipterus longi* | Gogo | NA | NA | 171 | 85.5 | NA | 20 | NA | NA | present | sub-terminal/inferior | lateral | absent | heterocercal |
| *Rolfosteus canningensis* | Gogo | 440 | 440 | 115.2 | 55 | 35 | 17.7 | fusiform/normal | circular | present | sub-terminal/inferior | lateral | absent | heterocercal |
| *Scaumenacia curta* | Miguasha | 685 | 637 | 122 | 39 | 137 | 28 | fusiform/normal | compressed | present | terminal | lateral | present | heterocercal |
| *Simosteus tuberculatus* | Gogo | 650 | NA | 125 | 22 | NA | 22 | fusiform/normal | oval | present | terminal | lateral | present | heterocercal |
| *Soederberghia simpsoni* | Canowindra | 196 | NA | 88.3 | 26.5 | 48.1 | 8.8 | elongated | compressed | present | terminal | lateral | NA | heterocercal |
| *Torosteus pulchellus* | Gogo | 300 | 320 | 67.6 | 6 | 80 | 13.6 | fusiform/normal | circular | present | terminal | lateral | present | heterocercal |
| *Torosteus sp. ?* | Gogo | 200 | 200 | NA | NA | NA | NA | fusiform/normal | circular | present | NA | lateral | NA | heterocercal |
| *Torosteus tuberculatus* | Gogo | 420 | 420 | 83.2 | 8 | 92 | 17 | fusiform/normal | circular | present | terminal | lateral | present | heterocercal |
| *Triazeugacanthus affinis* | Miguasha | 56 | 56 | 10 | 1 | 11 | 3 | elongated | oval | present | terminal | lateral | absent | heterocercal |
| *Tubonasus lennardensis* | Gogo | 210 | 210 | 70 | 28 | 39 | 14 | elongated | angular | present | sub-terminal/inferior | lateral | absent | heterocercal |
| *undescribed chondrich.* | Gogo | 150 | NA | NA | NA | NA | NA | elongated | circular | present | NA | lateral | present | heterocercal |
| *Xeradipterus hatcheri* | Gogo | NA | NA | 90 | 26 | 54 | 14 | NA | circular | present | sub-terminal/inferior | lateral | absent | NA |

**Table S2** Discovery curve data for Gogo Formation.

| Group | Taxon | Year Found | Described/named by | Year described | Subsequent synonymising etc? | Notes | Original Name |
|---|---|---|---|---|---|---|---|
| Acanthodian | *Halimacanthodes ahlbergi* | 2008 | Burrows et al. | 2012 | | | |
| Chondrichthyan | *Gogoselachus lynnbeazleyae* | 2005 | Long et al. | 2015 | | | |
| | undescribed chondrich. | 2005 | | ? | | | |
| Antiarch (placoderm) | *Bothriolepis* sp. | 1963 | | ? | | | |
| Ptyctodontid (placoderm) | *Austroptyctodus gardineri* | 1967 | Miles & Young | 1977 | Renamed by Long 1997 | | *Ctenurella gardineri* |
| | *Campbellodus decipiens* | 1967 | Mikes & Young | 1977 | | | |
| | *Materpiscis attenboroughi* | 2005 | Long et al | 2008 | | | |
| Arthrodire (placoderm) | *Bruntonichthys multidens* | 1967 | Dennis & Miles | 1980 | | | |
| | *Bullerichthys fascidens* | 1967 | Dennis & Miles | 1980 | | | |
| | *Camuropiscis concinnus* | 1967 | Dennis & Miles | 1979 | | | |
| | *Camuropiscis laidlawi* | 1967 | Dennis & Miles | 1979 | | | |
| | *Compagopiscis croucheri* | 1967 | Gardiner & Miles | 1994 | | | |
| | *Eastmanosteus calliaspis* | 1967 | Dennis-Bryant | 1987 | | currently being revised | |
| | *Eastmanosteus sp. undescribed sp. 1* | | | | | | |
| | *Eastmanosteus sp. undescribed sp. 2* | | | | | | |
| | *Fallacosteus turneri* | 1986 | Long | 1990 | | | |
| | *Gogopiscis gracilis* | 1967 | Gardiner & Miles | 1994 | *Compagopiscis croucheri* TrinajsSc& Hazelton 2007 | | |
| | *Harrytoombsia elegans* | 1967 | Miles & Dennis | 1979 | | | |
| | *Holonema westolli* | 1967 | Miles | 1971 | | | |
| | *Incisoscutum ritchiei* | 1967 | Dennis& Miles | 1981 | | | |
| | *Incisoscutum sarahae* | 1986 | Long | 1994 | Genus changed by TrinajsSc& Dennis Bryan 2009 | | *Gogosteus sarahae* |
| | *Kendrickichthys cavernosus* | 1967 | Dennis & Miles | 1980 | | | |
| | *Kimberleyichthys bispicatus* | 1967 | Dennis & Miles | 1983 | | | |
| | *Kimberleyichthys whybrowi* | 1967 | Dennis & Miles | 1983 | | | |
| | *Latocamurus coulthardi* | 1986 | Long | 1988 | | | |
| | *Mcnamaraspis kaprios* | 1986 | Long | 1995 | there was a specimen in the BMNH misideniWied | | |
| | *Pinguosteus thulborni* | 1986 | Long | 1990 | | | |
| | *Rolfosteus canningensis* | 1967 | Dennis & Miles | 1979 | | | |
| | *Torosteus sp. ?* | | | | | | |
| | *Torosteus tuberculatus* | 1967 | Gardiner & Miles | 1990 | | | |
| | *Torosteus pulchellus* | 1967 | Gardiner & Miles | 1990 | | | |
| | *Tubonasus lennardensis* | 1967 | Dennis & Miles | 1979 | | | |
| | *Simosteus* | unknown | Dennis & Miles | 1982 | found in collecSon of the Australian Museum | specimen destroyed durin transport for the NHM Lonond | |
| Onychodont (sarcopt) | *Onychodus jandemarrai* | 1967 | Andrews et al. | 2006 | | | |

| | | | | | | |
|---|---|---|---|---|---|---|
| Actinistia (sarcopt) | *Namugawi wirngarri* | 2008 | Clement et al. | 2024 | | Still under review/in press |
| Dipnoi (sarcopt) | *Adololopas moyasmithae* | 1986 | Campbell & Barwick | 1998 | | |
| | *Asthenorhynchus meemannae* | 2001 | Pridmore et al. | 1994 | Elevated to new genus Long 2010 | |
| | *Chirodipterus australis* | 1967 | Miles | 1977 | | |
| | *Gogodipterus paddyensis* | 1967 | Miles | 1977 | Elevated to new genus Long 1992 | |
| | *Griphognathus whitei* | 1967 | Miles | 1977 | | |
| | *Holodipterus elderae* | 1967 | Pridmore et al. | 1994 | | |
| | *Holodipterus gogoensis* | 1967 | Miles | 1977 | | |
| | *Pillararhynhcus longi* | 1986 | Barwick & Campbell | 1996 | | |
| | *Rhinodipterus kimberleyensis* | ?2008 | Clement | 2012 | | |
| | *Robinsondipterus longi* | 1986 | Campbell & Barwick | 1991 | Elevated to new genus Long 2010 | |
| | *Xeradipterus hatcheri* | 2005 | Clement & Long | 2010 | | |
| Tetrapodomorph (sarcopt) | *Gogonasus andrewsae* | 1967 | Long | 1985 | | |
| Other (sarcopt) | *Cainocara enigma* | | Campbell & Barwick | 2011 | Nomen nudum - non taxon | In Long & Trinajstic 2018 review |
| | Kapitany sarcopt | | ? | | | Still not formally described - include date of it being seized? |
| Actinopterygian | *Gogosardina coatesi* | 2001 | Choo et al. | 2009 | | |
| | *Mimipiscis bartrami* | 2005 | Gardiner & Bartram | 1977 | Mimia renamed to Mimipiscis (Choo 2011) | Collected in 60s |
| | *Mimipiscis toombsi* | 1967 | Gardiner & Bartram | 1977 | Mimia renamed to Mimipiscis (Choo 2011) | |
| | *Moythomasia durgaringa* | 1967 | Gardiner & Bartram | 1977 | Mimia renamed to Mimipiscis (Choo 2011) | |
| | *Pickeringius acanthophorus* | 1990 | Choo et al. | 2018 | | |

**Table S3** Details of the six modern fish communities.

| community name | habitat | climate zone | fish diversity | latitude |
|---|---|---|---|---|
| Ythan | estuary | temperate | 17 | 57 N |
| Canal de Santa Cruz | estuary | tropical | 86 | 7 S |
| Central Chile subtidal | marine reef | temperate/ subtropical | 27 | 30-33 S |
| Caribbean | marine reef | tropical | 208 | 18 N |
| Nepean River (Richmond to Warragamba River) | freshwater | temperate | 19 | 33.75 S |
| Braço Morto Acima and Braço Morto Abaixo, Miranda River, Brazil | freshwater | tropical | 79 | 16 S |

**Table S4.** Observed and imputed traits for all study species. Total length (mm) is $\log_e$-transformed. All other continuous traits are expressed as a proportion of total length, with pre-orbital length also being $\log_e$-transformed. Some traits for some species from Gogo and Canowindra were inferred using multiple imputation.

| Species | BodyShapeI (sagittal) | BodyShapeII (transverse) | log(total length) | head length (relative to TL) | eye diameter (relative to TL) | log(pre-orbital length relative to TL) | body depth (relative to TL) | mouth position | eye position | spiracle | caudal fin shape |
|---|---|---|---|---|---|---|---|---|---|---|---|
| *Achirus declivis* | shortAndOrDeep | compressed and lies on side | 2.93 | 0.21 | 0.03 | -2.84 | 0.51 | sub-terminal/inferior | migrated to one side and raised | absent | homocercal |
| *Achirus lineatus* | shortAndOrDeep | compressed and lies on side | 3.50 | 0.22 | 0.03 | -2.64 | 0.56 | sub-terminal/inferior | migrated to one side and raised | absent | homocercal |
| *Anchoa lyolepis* | elongated | oval | 2.48 | 0.21 | 0.06 | -3.25 | 0.17 | sub-terminal/inferior | lateral | absent | homocercal |
| *Anchoa marinii* | elongated | oval | 2.80 | 0.19 | 0.04 | -3.16 | 0.18 | sub-terminal/inferior | lateral | absent | homocercal |
| *Anchoa spinifer* | elongated | oval | 3.18 | 0.19 | 0.04 | -3.82 | 0.21 | sub-terminal/inferior | lateral | absent | homocercal |
| *Anchoa tricolor* | elongated | oval | 2.70 | 0.20 | 0.06 | -3.15 | 0.17 | sub-terminal/inferior | lateral | absent | homocercal |
| *Anchovia clupeoides* | fusiform | oval | 3.40 | 0.21 | 0.05 | -3.55 | 0.24 | sub-terminal/inferior | lateral | absent | homocercal |
| *Archosargus rhomboidalis* | shortAndOrDeep | compressed | 3.50 | 0.23 | 0.06 | -2.52 | 0.38 | terminal | lateral | absent | homocercal |
| *Bathygobius soporator* | elongated | compressed | 2.71 | 0.26 | 0.06 | -2.76 | 0.18 | terminal | raised/top of head | absent | homocercal |
| *Batrachoides surinamensis* | fusiform | circular | 4.04 | 0.27 | 0.02 | -3.03 | 0.17 | terminal | raised/top of head | absent | homocercal |
| *Caranx crysos* | fusiform | compressed | 4.25 | 0.21 | 0.05 | -2.80 | 0.29 | terminal | lateral | absent | homocercal |
| *Caranx hippos* | fusiform | compressed | 4.82 | 0.25 | 0.06 | -2.88 | 0.33 | terminal | lateral | absent | homocercal |
| *Caranx latus* | fusiform | compressed | 4.76 | 0.25 | 0.06 | -2.60 | 0.30 | terminal | lateral | absent | homocercal |
| *Centropomus parallelus* | fusiform | compressed | 4.28 | 0.28 | 0.07 | -2.42 | 0.22 | terminal | lateral | absent | homocercal |
| *Centropomus pectinatus* | fusiform | compressed | 4.03 | 0.26 | 0.05 | -2.53 | 0.23 | terminal | lateral | absent | homocercal |
| *Centropomus undecimalis* | fusiform | compressed | 4.94 | 0.31 | 0.04 | -2.42 | 0.19 | terminal | lateral | absent | homocercal |
| *Cetengraulis edentulus* | fusiform | compressed | 3.02 | 0.24 | 0.07 | -3.28 | 0.26 | sub-terminal/inferior | lateral | absent | homocercal |
| *Chirocentrodon bleekerianus* | elongated | compressed | 2.78 | 0.20 | 0.06 | -2.67 | 0.19 | terminal | lateral | absent | homocercal |
| *Chloroscombrus chrysurus* | fusiform | compressed | 4.17 | 0.20 | 0.05 | -2.79 | 0.34 | superior | lateral | absent | homocercal |
| *Citharichthys spilopterus* | shortAndOrDeep | compressed and lies on side | 3.04 | 0.22 | 0.04 | -3.10 | 0.37 | terminal | migrated to one side and raised | absent | homocercal |
| *Ctenogobius boleosoma* | elongated | compressed | 2.01 | 0.21 | 0.05 | -2.88 | 0.12 | sub-terminal/inferior | raised/top of head | absent | homocercal |
| *Ctenogobius shufeldti* | elongated | compressed | 2.08 | 0.21 | 0.04 | -2.79 | 0.13 | sub-terminal/inferior | raised/top of head | absent | homocercal |
| *Ctenogobius smaragdus* | elongated | compressed | 2.71 | 0.17 | 0.03 | -3.29 | 0.11 | sub-terminal/inferior | raised/top of head | absent | homocercal |
| *Ctenogobius stigmaticus* | elongated | compressed | 2.08 | 0.19 | 0.05 | -3.11 | 0.13 | sub-terminal/inferior | raised/top of head | absent | homocercal |
| *Diapterus auratus* | fusiform | compressed | 3.76 | 0.25 | 0.09 | -2.62 | 0.35 | terminal | lateral | absent | homocercal |
| *Diapterus rhombeus* | shortAndOrDeep | compressed | 3.69 | 0.26 | 0.08 | -2.71 | 0.40 | terminal | lateral | absent | homocercal |
| *Etropus crossotus* | shortAndOrDeep | compressed and lies on side | 3.06 | 0.16 | 0.04 | -3.60 | 0.44 | superior | migrated to one side and raised | absent | homocercal |
| *Etropus longimanus* | fusiform | compressed and lies on side | 2.74 | 0.18 | 0.05 | -3.53 | 0.34 | terminal | migrated to one side and raised | absent | homocercal |
| *Gobionellus oceanicus* | elongated | compressed | 3.30 | 0.13 | 0.03 | -3.28 | 0.09 | terminal | raised/top of head | absent | homocercal |
| *Gobionellus stomatus* | elongated | compressed | 2.63 | 0.16 | 0.03 | -3.05 | 0.11 | terminal | raised/top of head | absent | homocercal |
| *Hyporhamphus unifasciatus* | elongated | compressed | 3.40 | 0.31 | 0.02 | -1.61 | 0.11 | superior | lateral | absent | homocercal |

| | | | | | | | | | | | |
|---|---|---|---|---|---|---|---|---|---|---|---|
| *Lagocephalus laevigatus* | shortAndOrDeep | circular | 4.61 | 0.26 | 0.07 | -2.27 | 0.22 | terminal | lateral | absent | homocercal |
| *Lutjanus analis* | fusiform | oval | 4.54 | 0.26 | 0.04 | -2.35 | 0.30 | terminal | lateral | absent | homocercal |
| *Lutjanus jocu* | fusiform | oval | 4.85 | 0.29 | 0.05 | -2.20 | 0.31 | terminal | lateral | absent | homocercal |
| *Lutjanus synagris* | fusiform | oval | 4.09 | 0.29 | 0.06 | -2.47 | 0.35 | terminal | lateral | absent | homocercal |
| *Lycengraulis grossidens* | fusiform | compressed | 3.31 | 0.18 | 0.04 | -3.85 | 0.20 | sub-terminal/inferior | lateral | absent | homocercal |
| *Microgobius meeki* | elongated | compressed | 2.71 | 0.23 | 0.06 | -2.88 | 0.14 | superior | raised/top of head | absent | homocercal |
| *Mugil curema* | fusiform | circular | 4.51 | 0.21 | 0.06 | -3.17 | 0.21 | sub-terminal/inferior | lateral | absent | homocercal |
| *Oligoplites palometa* | fusiform | compressed | 3.91 | 0.18 | 0.04 | -3.13 | 0.24 | superior | lateral | absent | homocercal |
| *Oligoplites saliens* | fusiform | compressed | 3.91 | 0.16 | 0.05 | -3.62 | 0.26 | superior | lateral | absent | homocercal |
| *Oligoplites saurus* | fusiform | compressed | 3.56 | 0.19 | 0.04 | -2.85 | 0.22 | superior | lateral | absent | homocercal |
| *Opisthonema oglinum* | fusiform | compressed | 3.64 | 0.19 | 0.05 | -2.86 | 0.26 | terminal | lateral | absent | homocercal |
| *Paralichthys brasiliensis* | shortAndOrDeep | compressed and lies on side | 4.61 | 0.22 | 0.04 | -2.97 | 0.35 | superior | migrated to one side and raised | absent | homocercal |
| *Platanichthys platana* | fusiform | compressed | 3.12 | 0.18 | 0.04 | -3.04 | 0.26 | terminal | lateral | absent | homocercal |
| *Rhinosardinia bahiensis* | fusiform | compressed | 2.26 | 0.17 | 0.06 | -3.16 | 0.23 | terminal | lateral | absent | homocercal |
| *Sparisoma radians* | fusiform | compressed | 3.00 | 0.29 | 0.10 | -2.77 | 0.30 | sub-terminal/inferior | lateral | absent | homocercal |
| *Sphoeroides testudineus* | fusiform | circular | 3.66 | 0.30 | 0.07 | -2.23 | 0.29 | sub-terminal/inferior | raised/top of head | absent | homocercal |
| *Sphyraena barracuda* | elongated | compressed | 5.30 | 0.24 | 0.04 | -2.20 | 0.13 | superior | lateral | absent | homocercal |
| *Sphyraena guachancho* | elongated | compressed | 5.30 | 0.21 | 0.04 | -2.48 | 0.12 | superior | lateral | absent | homocercal |
| *Thalassophryne nattereri* | elongated | circular | 3.06 | 0.31 | 0.02 | -3.15 | 0.27 | superior | raised/top of head | absent | homocercal |
| *Lutjanus alexandrei* | fusiform | oval | 3.50 | 0.30 | 0.07 | -2.60 | 0.35 | terminal | lateral | absent | homocercal |
| *Sphoeroides greeleyi* | shortAndOrDeep | circular | 2.89 | 0.27 | 0.06 | -2.03 | 0.23 | sub-terminal/inferior | raised/top of head | absent | homocercal |
| *Archosargus probatocephalus* | shortAndOrDeep | compressed | 4.51 | 0.26 | 0.05 | -2.67 | 0.38 | terminal | lateral | absent | homocercal |
| *Atherinella brasiliensis* | elongated | oval | 2.87 | 0.19 | 0.05 | -2.83 | 0.17 | terminal | lateral | absent | homocercal |
| *Bairdiella ronchus* | shortAndOrDeep | compressed | 3.56 | 0.25 | 0.07 | -2.69 | 0.25 | sub-terminal/inferior | lateral | absent | homocercal |
| *Cathorops agassizii* | fusiform | circular | 3.11 | 0.28 | 0.03 | -2.59 | 0.19 | sub-terminal/inferior | lateral | absent | homocercal |
| *Chaetodipterus faber* | shortAndOrDeep | compressed | 4.51 | 0.20 | 0.05 | -2.74 | 0.52 | terminal | lateral | absent | homocercal |
| *Chaetodon ocellatus* | shortAndOrDeep | compressed | 3.00 | 0.33 | 0.10 | -2.18 | 0.64 | terminal | lateral | absent | homocercal |
| *Cynoscion virescens* | fusiform | oval | 4.74 | 0.21 | 0.03 | -3.09 | 0.20 | superior | lateral | absent | homocercal |
| *Elops saurus* | elongated | compressed | 4.61 | 0.17 | 0.03 | -3.16 | 0.17 | terminal | lateral | absent | homocercal |
| *Engraulis anchoita* | elongated | oval | 2.98 | 0.20 | 0.04 | -3.32 | 0.15 | sub-terminal/inferior | lateral | absent | homocercal |
| *Epinephelus adscensionis* | fusiform | compressed | 4.17 | 0.33 | 0.05 | -2.44 | 0.28 | terminal | lateral | absent | homocercal |
| *Epinephelus marginatus* | fusiform | compressed | 5.01 | 0.32 | 0.05 | -2.66 | 0.28 | superior | lateral | absent | homocercal |
| *Eucinostomus argenteus* | fusiform | compressed | 3.05 | 0.29 | 0.09 | -2.22 | 0.28 | terminal | lateral | absent | homocercal |
| *Eucinostomus gula* | fusiform | compressed | 3.24 | 0.27 | 0.09 | -2.51 | 0.32 | terminal | lateral | absent | homocercal |
| *Eucinostomus havana* | fusiform | compressed | 2.89 | 0.27 | 0.09 | -2.59 | 0.31 | terminal | lateral | absent | homocercal |
| *Eucinostomus melanopterus* | fusiform | compressed | 3.40 | 0.25 | 0.06 | -2.50 | 0.32 | terminal | lateral | absent | homocercal |
| *Eugerres brasilianus* | shortAndOrDeep | compressed | 4.08 | 0.28 | 0.07 | -2.27 | 0.36 | terminal | lateral | absent | homocercal |
| *Evorthodus lyricus* | fusiform | compressed | 2.71 | 0.17 | 0.04 | -3.18 | 0.14 | sub-terminal/inferior | raised/top of head | absent | homocercal |
| *Genyatremus luteus* | shortAndOrDeep | compressed | 3.61 | 0.23 | 0.08 | -3.28 | 0.38 | sub-terminal/inferior | lateral | absent | homocercal |

| | | | | | | | | | | | |
|---|---|---|---|---|---|---|---|---|---|---|---|
| *Guavina guavina* | fusiform | compressed | 3.57 | 0.27 | 0.05 | -2.94 | 0.17 | terminal | raised/top of head | absent | homocercal |
| *Harengula clupeola* | fusiform | compressed | 3.11 | 0.21 | 0.07 | -2.73 | 0.27 | terminal | lateral | absent | homocercal |
| *Hemiramphus brasiliensis* | elongated | compressed | 4.01 | 0.33 | 0.03 | -1.43 | 0.11 | superior | lateral | absent | homocercal |
| *Hypanus guttatus* | other | flattened | 6.11 | 0.24 | 0.01 | -2.43 | 0.03 | sub-terminal/inferior | raised/top of head | present | whip-like |
| *Mycteroperca bonaci* | fusiform | oval | 5.01 | 0.28 | 0.03 | -2.43 | 0.27 | superior | lateral | absent | homocercal |
| *Polydactylus virginicus* | fusiform | compressed | 3.50 | 0.19 | 0.07 | -3.62 | 0.20 | sub-terminal/inferior | lateral | absent | homocercal |
| *Prionotus punctatus* | fusiform | compressed | 3.81 | 0.29 | 0.07 | -2.22 | 0.21 | sub-terminal/inferior | raised/top of head | absent | homocercal |
| *Rhonciscus crocro* | fusiform | compressed | 3.64 | 0.30 | 0.07 | -2.46 | 0.33 | sub-terminal/inferior | lateral | absent | homocercal |
| *Sardinella brasiliensis* | fusiform | compressed | 3.30 | 0.19 | 0.05 | -2.88 | 0.19 | terminal | lateral | absent | homocercal |
| *Sciades herzbergii* | elongated | circular | 4.55 | 0.22 | 0.04 | -2.80 | 0.18 | sub-terminal/inferior | lateral | absent | homocercal |
| *Synodus foetens* | elongated | circular | 3.99 | 0.18 | 0.04 | -3.21 | 0.12 | terminal | raised/top of head | absent | homocercal |
| *Tylosurus acus* | elongated | circular | 5.03 | 0.29 | 0.03 | -1.71 | 0.07 | terminal | lateral | absent | homocercal |
| *Gymnothorax ocellatus* | eel-like | compressed | 4.50 | 0.14 | 0.01 | -3.75 | 0.08 | terminal | lateral | absent | isocercal |
| *Sparisoma axillare* | fusiform | compressed | 3.94 | 0.24 | 0.04 | -2.44 | 0.28 | sub-terminal/inferior | lateral | absent | homocercal |
| *Stellifer stellifer* | fusiform | compressed | 3.04 | 0.25 | 0.06 | -2.79 | 0.26 | sub-terminal/inferior | lateral | absent | homocercal |
| *Syngnathus scovelli* | eel-like | circular | 2.95 | 0.12 | 0.01 | -2.84 | 0.05 | terminal | lateral | absent | homocercal |
| *Acestrorhynchus falcatus* | elongated | compressed | 3.40 | 0.25 | 0.04 | -2.50 | 0.25 | terminal | lateral | absent | homocercal |
| *Acestrorhynchus lacustris* | elongated | compressed | 3.61 | 0.25 | 0.04 | -2.43 | 0.20 | terminal | lateral | absent | homocercal |
| *Acestrorhynchus pantaneiro* | elongated | compressed | 3.56 | 0.25 | 0.05 | -2.47 | 0.21 | sub-terminal/inferior | lateral | absent | homocercal |
| *Astyanax altiparanae* | shortAndOrDeep | compressed | 2.80 | 0.21 | 0.07 | -3.10 | 0.34 | superior | lateral | absent | homocercal |
| *Astyanax bimaculatus* | shortAndOrDeep | compressed | 2.86 | 0.23 | 0.09 | -3.06 | 0.27 | superior | lateral | absent | homocercal |
| *Auchenipterus nuchalis* | elongated | oval | 3.45 | 0.17 | 0.05 | -3.21 | 0.17 | terminal | lateral | absent | homocercal |
| *Bujurquina vittata* | fusiform | compressed | 2.53 | 0.26 | 0.06 | -2.36 | 0.33 | terminal | lateral | absent | homocercal |
| *Callichthys callichthys* | elongated | oval | 3.04 | 0.20 | 0.02 | -2.68 | 0.18 | sub-terminal/inferior | lateral | absent | homocercal |
| *Chaetobranchopsis australis* | shortAndOrDeep | compressed | 2.72 | 0.30 | 0.07 | -2.32 | 0.43 | superior | lateral | absent | homocercal |
| *Corydoras aeneus* | fusiform | oval | 2.30 | 0.24 | 0.05 | -2.18 | 0.29 | sub-terminal/inferior | lateral | absent | homocercal |
| *Corydoras britskii* | elongated | oval | 2.39 | 0.29 | 0.06 | -1.84 | 0.34 | sub-terminal/inferior | lateral | absent | homocercal |
| *Corydoras pantanalensis* | shortAndOrDeep | oval | 2.21 | 0.27 | 0.05 | -2.51 | 0.32 | sub-terminal/inferior | lateral | absent | homocercal |
| *Corydoras splendens* | shortAndOrDeep | oval | 2.62 | 0.22 | 0.05 | -2.14 | 0.28 | sub-terminal/inferior | lateral | absent | homocercal |
| *Curimatella alburnus* | fusiform | compressed | 3.07 | 0.18 | 0.05 | -2.81 | 0.29 | sub-terminal/inferior | lateral | absent | homocercal |
| *Curimatella dorsalis* | fusiform | oval | 2.70 | 0.20 | 0.08 | -3.11 | 0.21 | terminal | lateral | absent | homocercal |
| *Cyphocharax gillii* | fusiform | compressed | 2.58 | 0.21 | 0.07 | -2.93 | 0.28 | terminal | lateral | absent | homocercal |
| *Eigenmannia trilineata* | elongated | compressed | 3.22 | 0.12 | 0.02 | -3.43 | 0.17 | sub-terminal/inferior | lateral | absent | reduced or absent |
| *Entomocorus benjamini* | elongated | compressed | 2.21 | 0.17 | 0.05 | -3.00 | 0.22 | terminal | lateral | absent | homocercal |
| *Gymnogeophagus balzanii* | shortAndOrDeep | compressed | 2.87 | 0.25 | 0.05 | -2.41 | 0.37 | terminal | lateral | absent | homocercal |
| *Hemiodus microlepis* | fusiform | compressed | 3.37 | 0.17 | 0.05 | -3.11 | 0.23 | terminal | lateral | absent | homocercal |
| *Hemiodus orthonops* | fusiform | oval | 3.43 | 0.17 | 0.05 | -2.99 | 0.18 | terminal | lateral | absent | homocercal |
| *Hemisorubim platyrhynchos* | elongated | circular | 4.11 | 0.26 | 0.03 | -2.24 | 0.14 | superior | raised/top of head | absent | homocercal |
| *Hoplias malabaricus* | elongated | oval | 4.17 | 0.26 | 0.04 | -2.94 | 0.19 | terminal | lateral | absent | homocercal |

| | | | | | | | | | | | |
|---|---|---|---|---|---|---|---|---|---|---|---|
| *Hypoptopoma inexspectatum* | elongated | oval | 2.19 | 0.24 | 0.04 | -2.32 | 0.12 | sub-terminal/inferior | lateral | absent | homocercal |
| *Hypostomus boulengeri* | elongated | angular | 3.54 | 0.27 | 0.04 | -1.93 | 0.18 | sub-terminal/inferior | raised/top of head | absent | homocercal |
| *Hypostomus cochliodon* | elongated | angular | 3.42 | 0.16 | 0.03 | -2.47 | 0.18 | sub-terminal/inferior | raised/top of head | absent | homocercal |
| *Hypostomus plecostomus* | elongated | angular | 4.16 | 0.31 | 0.04 | -1.96 | 0.19 | sub-terminal/inferior | raised/top of head | absent | homocercal |
| *Leporinus friderici* | fusiform | compressed | 3.87 | 0.19 | 0.04 | -2.86 | 0.28 | terminal | lateral | absent | homocercal |
| *Leporinus lacustris* | fusiform | oval | 3.18 | 0.19 | 0.04 | -2.79 | 0.28 | terminal | lateral | absent | homocercal |
| *Loricaria coximensis* | elongated | angular | 2.41 | 0.18 | 0.02 | -2.44 | 0.10 | sub-terminal/inferior | raised/top of head | absent | homocercal |
| *Loricaria luciae* | elongated | angular | 3.08 | 0.17 | 0.03 | -2.47 | 0.06 | sub-terminal/inferior | raised/top of head | absent | homocercal |
| *Loricariichthys labialis* | elongated | angular | 3.19 | 0.19 | 0.02 | -3.20 | 0.09 | sub-terminal/inferior | raised/top of head | absent | homocercal |
| *Loricariichthys platymetopon* | elongated | angular | 3.65 | 0.14 | 0.03 | -2.74 | 0.11 | sub-terminal/inferior | raised/top of head | absent | homocercal |
| *Megaleporinus elongatus* | elongated | compressed | 3.91 | 0.18 | 0.04 | -2.68 | 0.23 | sub-terminal/inferior | lateral | absent | homocercal |
| *Moenkhausia oligolepis* | shortAndOrDeep | compressed | 2.57 | 0.19 | 0.08 | -3.01 | 0.29 | superior | lateral | absent | homocercal |
| *Moenkhausia bonita* | fusiform | compressed | 2.40 | 0.18 | 0.07 | -2.98 | 0.23 | superior | lateral | absent | homocercal |
| *Moenkhausia dichroura* | fusiform | compressed | 2.49 | 0.20 | 0.08 | -3.11 | 0.24 | superior | lateral | absent | homocercal |
| *Moenkhausia forestii* | shortAndOrDeep | compressed | 1.61 | 0.22 | 0.08 | -3.31 | 0.34 | superior | lateral | absent | homocercal |
| *Moenkhausia intermedia* | fusiform | oval | 2.34 | 0.18 | 0.10 | -3.60 | 0.28 | superior | lateral | absent | homocercal |
| *Moenkhausia sanctaefilomenae* | shortAndOrDeep | compressed | 2.19 | 0.19 | 0.08 | -3.07 | 0.31 | superior | lateral | absent | homocercal |
| *Pimelodella gracilis* | elongated | compressed | 3.05 | 0.15 | 0.04 | -3.18 | 0.11 | sub-terminal/inferior | lateral | absent | homocercal |
| *Pimelodella mucosa* | elongated | compressed | 2.84 | 0.20 | 0.05 | -2.82 | 0.19 | sub-terminal/inferior | lateral | absent | homocercal |
| *Pimelodella taenioptera* | elongated | compressed | 2.77 | 0.14 | 0.03 | -2.87 | 0.13 | sub-terminal/inferior | lateral | absent | homocercal |
| *Pimelodus argenteus* | elongated | compressed | 3.39 | 0.22 | 0.05 | -2.40 | 0.21 | sub-terminal/inferior | lateral | absent | homocercal |
| *Pimelodus maculatus* | fusiform | compressed | 3.93 | 0.21 | 0.05 | -2.46 | 0.25 | sub-terminal/inferior | lateral | absent | homocercal |
| *Plagioscion ternetzi* | elongated | oval | 3.82 | 0.28 | 0.04 | -2.86 | 0.26 | terminal | lateral | absent | homocercal |
| *Poptella paraguayensis* | shortAndOrDeep | compressed | 2.57 | 0.23 | 0.09 | -2.60 | 0.46 | superior | lateral | absent | homocercal |
| *Potamorhina squamoralevis* | fusiform | compressed | 3.23 | 0.28 | 0.06 | -2.45 | 0.32 | terminal | lateral | absent | homocercal |
| *Prochilodus lineatus* | fusiform | compressed | 4.38 | 0.20 | 0.04 | -2.81 | 0.29 | terminal | lateral | absent | homocercal |
| *Psectrogaster curviventris* | fusiform | compressed | 3.04 | 0.23 | 0.07 | -3.07 | 0.39 | superior | lateral | absent | homocercal |
| *Pseudoplatystoma corruscans* | fusiform | circular | 5.11 | 0.30 | 0.02 | -2.02 | 0.17 | sub-terminal/inferior | raised/top of head | absent | homocercal |
| *Pseudoplatystoma fasciatum* | fusiform | circular | 4.83 | 0.29 | 0.02 | -1.96 | 0.15 | sub-terminal/inferior | raised/top of head | absent | homocercal |
| *Pseudoplatystoma reticulatum* | elongated | circular | 4.20 | 0.31 | 0.03 | -1.99 | 0.16 | sub-terminal/inferior | raised/top of head | absent | homocercal |
| *Pterygoplichthys anisitsi* | elongated | angular | 4.01 | 0.28 | 0.03 | -2.13 | 0.19 | sub-terminal/inferior | raised/top of head | absent | homocercal |
| *Pygocentrus nattereri* | shortAndOrDeep | compressed | 4.07 | 0.28 | 0.06 | -2.94 | 0.45 | superior | lateral | absent | homocercal |
| *Rhamphichthys hahni* | elongated | compressed | 3.28 | 0.12 | 0.01 | -2.86 | 0.10 | sub-terminal/inferior | lateral | absent | reduced or absent |
| *Rineloricaria nigricauda* | elongated | angular | 2.08 | 0.17 | 0.02 | -2.56 | 0.07 | sub-terminal/inferior | raised/top of head | absent | homocercal |
| *Rineloricaria parva* | elongated | angular | 2.57 | 0.15 | 0.02 | -2.63 | 0.07 | sub-terminal/inferior | raised/top of head | absent | homocercal |
| *Roeboides descalvadensis* | fusiform | compressed | 2.37 | 0.19 | 0.08 | -3.28 | 0.30 | superior | lateral | absent | homocercal |
| *Salminus brasiliensis* | fusiform | oval | 4.74 | 0.24 | 0.03 | -2.95 | 0.24 | terminal | lateral | absent | homocercal |

| | | | | | | | | | | | |
|---|---|---|---|---|---|---|---|---|---|---|---|
| *Satanoperca pappaterra* | shortAndOrDeep | compressed | 3.30 | 0.23 | 0.09 | -2.56 | 0.30 | terminal | lateral | absent | homocercal |
| *Schizodon borellii* | fusiform | oval | 3.58 | 0.19 | 0.05 | -2.86 | 0.23 | terminal | lateral | absent | homocercal |
| *Serrasalmus marginatus* | shortAndOrDeep | compressed | 3.30 | 0.28 | 0.07 | -3.17 | 0.41 | superior | lateral | absent | homocercal |
| *Serrasalmus spilopleura* | shortAndOrDeep | compressed | 3.26 | 0.28 | 0.09 | -2.99 | 0.47 | superior | lateral | absent | homocercal |
| *Steindachnerina brevipinna* | fusiform | oval | 3.12 | 0.19 | 0.06 | -3.32 | 0.26 | sub-terminal/inferior | lateral | absent | homocercal |
| *Steindachnerina conspersa* | elongated | compressed | 2.78 | 0.19 | 0.07 | -3.19 | 0.29 | sub-terminal/inferior | lateral | absent | homocercal |
| *Sturisoma robustum* | elongated | angular | 3.47 | 0.19 | 0.02 | -2.15 | 0.10 | sub-terminal/inferior | raised/top of head | absent | homocercal |
| *Tetragonopterus argenteus* | shortAndOrDeep | compressed | 2.62 | 0.22 | 0.09 | -2.74 | 0.44 | superior | lateral | absent | homocercal |
| *Thoracocharax stellatus* | shortAndOrDeep | compressed | 2.16 | 0.23 | 0.08 | -3.39 | 0.45 | superior | lateral | absent | homocercal |
| *Trachelyopterus striatulus* | elongated | compressed | 3.32 | 0.21 | 0.02 | -3.11 | 0.22 | terminal | lateral | absent | homocercal |
| *Trachydoras paraguayensis* | elongated | compressed | 3.18 | 0.21 | 0.07 | -2.72 | 0.27 | sub-terminal/inferior | lateral | absent | homocercal |
| *Triportheus nematurus* | shortAndOrDeep | compressed | 3.06 | 0.24 | 0.06 | -2.62 | 0.33 | superior | lateral | absent | homocercal |
| *Triportheus pantanensis* | shortAndOrDeep | compressed | 2.94 | 0.18 | 0.04 | -3.26 | 0.32 | superior | lateral | absent | homocercal |
| *Astyanax abramis* | shortAndOrDeep | compressed | 2.86 | 0.22 | 0.07 | -2.63 | 0.39 | superior | lateral | absent | homocercal |
| *Astyanax asuncionensis* | shortAndOrDeep | compressed | 2.91 | 0.20 | 0.07 | -3.36 | 0.32 | superior | lateral | absent | homocercal |
| *Corydoras hastatus* | fusiform | compressed | 1.13 | 0.19 | 0.07 | -2.74 | 0.24 | sub-terminal/inferior | lateral | absent | homocercal |
| *Eigenmannia virescens* | elongated | compressed | 3.78 | 0.10 | 0.02 | -3.49 | 0.13 | sub-terminal/inferior | lateral | absent | reduced or absent |
| *Leporinus striatus* | fusiform | oval | 3.22 | 0.19 | 0.04 | -2.51 | 0.21 | terminal | lateral | absent | homocercal |
| *Trachelyopterus galeatus* | fusiform | compressed | 3.40 | 0.22 | 0.03 | -3.15 | 0.25 | terminal | lateral | absent | homocercal |
| *Abudefduf saxatilis* | shortAndOrDeep | compressed | 3.13 | 0.21 | 0.07 | -2.95 | 0.42 | terminal | lateral | absent | homocercal |
| *Abudefduf taurus* | shortAndOrDeep | compressed | 3.22 | 0.27 | 0.06 | -2.25 | 0.40 | terminal | lateral | absent | homocercal |
| *Acanthostracion polygonium* | shortAndOrDeep | angular | 3.91 | 0.18 | 0.08 | -2.52 | 0.31 | sub-terminal/inferior | raised/top of head | absent | homocercal |
| *Acanthostracion quadricornis* | shortAndOrDeep | angular | 4.01 | 0.14 | 0.05 | -2.52 | 0.31 | sub-terminal/inferior | raised/top of head | absent | homocercal |
| *Acanthurus bahianus* | shortAndOrDeep | compressed | 3.61 | 0.24 | 0.09 | -2.38 | 0.42 | terminal | lateral | absent | homocercal |
| *Acanthurus chirurgus* | shortAndOrDeep | compressed | 3.66 | 0.22 | 0.07 | -2.14 | 0.41 | terminal | lateral | absent | homocercal |
| *Acanthurus coeruleus* | shortAndOrDeep | compressed | 3.66 | 0.22 | 0.06 | -2.15 | 0.50 | terminal | lateral | absent | homocercal |
| *Aetobatus narinari* | other | flattened | 6.21 | 0.17 | 0.01 | -2.85 | 0.09 | sub-terminal/inferior | raised/top of head | present | whip-like |
| *Alphestes afer* | fusiform | oval | 3.50 | 0.30 | 0.05 | -3.18 | 0.28 | terminal | lateral | absent | homocercal |
| *Aluterus schoepfii* | shortAndOrDeep | compressed | 4.11 | 0.22 | 0.04 | -1.64 | 0.33 | superior | lateral | absent | homocercal |
| *Aluterus scriptus* | shortAndOrDeep | compressed | 4.70 | 0.17 | 0.02 | -1.73 | 0.25 | superior | lateral | absent | homocercal |
| *Amblycirrhitus pinos* | fusiform | compressed | 2.48 | 0.30 | 0.08 | -2.80 | 0.34 | terminal | raised/top of head | absent | homocercal |
| *Anchoa hepsetus* | elongated | oval | 2.73 | 0.20 | 0.06 | -3.45 | 0.18 | sub-terminal/inferior | lateral | absent | homocercal |
| *Anisotremus surinamensis* | shortAndOrDeep | compressed | 4.33 | 0.27 | 0.08 | -3.02 | 0.38 | terminal | lateral | absent | homocercal |
| *Anisotremus virginicus* | shortAndOrDeep | compressed | 3.70 | 0.24 | 0.06 | -2.85 | 0.38 | terminal | lateral | absent | homocercal |
| *Antennarius multiocellatus* | shortAndOrDeep | compressed | 3.00 | 0.21 | 0.02 | -2.38 | 0.47 | superior | lateral | absent | homocercal |
| *Antennarius striatus* | shortAndOrDeep | compressed | 3.22 | 0.37 | 0.06 | -2.75 | 0.44 | superior | raised/top of head | absent | homocercal |
| *Apogon maculatus* | fusiform | oval | 2.41 | 0.27 | 0.10 | -2.88 | 0.29 | terminal | lateral | absent | homocercal |
| *Atherinomorus stipes* | elongated | compressed | 2.30 | 0.22 | 0.09 | -3.15 | 0.18 | terminal | lateral | absent | homocercal |
| *Aulostomus maculatus* | elongated | compressed | 4.61 | 0.28 | 0.02 | -1.62 | 0.07 | terminal | lateral | absent | homocercal |

| | | | | | | | | | | | |
|---|---|---|---|---|---|---|---|---|---|---|---|
| *Azurina cyanea* | fusiform | oval | 2.71 | 0.28 | 0.07 | -2.57 | 0.31 | terminal | lateral | absent | homocercal |
| *Azurina multilineata* | fusiform | compressed | 3.00 | 0.22 | 0.07 | -3.17 | 0.31 | terminal | lateral | absent | homocercal |
| *Balistes vetula* | shortAndOrDeep | compressed | 4.09 | 0.23 | 0.05 | -1.78 | 0.38 | terminal | lateral | absent | homocercal |
| *Bodianus rufus* | fusiform | compressed | 3.69 | 0.25 | 0.04 | -2.58 | 0.26 | terminal | lateral | absent | homocercal |
| *Bothus lunatus* | shortAndOrDeep | compressed and lies on side | 3.83 | 0.22 | 0.03 | -3.01 | 0.46 | terminal | migrated to one side and raised | absent | homocercal |
| *Bothus ocellatus* | shortAndOrDeep | compressed and lies on side | 2.89 | 0.20 | 0.06 | -3.36 | 0.52 | sub-terminal/inferior | migrated to one side and raised | absent | homocercal |
| *Brachygenys chrysargyreum* | fusiform | compressed | 3.14 | 0.23 | 0.09 | -3.16 | 0.27 | terminal | lateral | absent | homocercal |
| *Calamus bajonado* | shortAndOrDeep | compressed | 4.41 | 0.25 | 0.08 | -2.10 | 0.35 | terminal | lateral | absent | homocercal |
| *Calamus calamus* | shortAndOrDeep | compressed | 4.03 | 0.22 | 0.08 | -2.32 | 0.42 | terminal | lateral | absent | homocercal |
| *Calamus pennatula* | shortAndOrDeep | compressed | 3.61 | 0.27 | 0.07 | -2.01 | 0.38 | terminal | lateral | absent | homocercal |
| *Cantherhines macrocerus* | shortAndOrDeep | compressed | 3.83 | 0.24 | 0.05 | -1.64 | 0.47 | superior | lateral | absent | homocercal |
| *Cantherhines pullus* | shortAndOrDeep | compressed | 3.00 | 0.26 | 0.08 | -1.58 | 0.42 | terminal | lateral | absent | homocercal |
| *Canthidermis sufflamen* | shortAndOrDeep | compressed | 4.17 | 0.24 | 0.05 | -1.85 | 0.42 | terminal | lateral | absent | homocercal |
| *Canthigaster rostrata* | shortAndOrDeep | angular | 2.48 | 0.27 | 0.06 | -1.81 | 0.41 | terminal | raised/top of head | absent | homocercal |
| *Caranx bartholomaei* | fusiform | compressed | 4.61 | 0.26 | 0.06 | -2.34 | 0.30 | terminal | lateral | absent | homocercal |
| *Caranx ruber* | fusiform | compressed | 4.40 | 0.23 | 0.05 | -2.56 | 0.27 | terminal | lateral | absent | homocercal |
| *Carcharhinus acronotus* | fusiform | circular | 5.30 | 0.23 | 0.02 | -2.72 | 0.16 | sub-terminal/inferior | lateral | absent | heterocercal |
| *Carcharhinus falciformis* | fusiform | circular | 5.86 | 0.23 | 0.02 | -2.72 | 0.13 | sub-terminal/inferior | lateral | absent | heterocercal |
| *Carcharhinus leucas* | fusiform | circular | 5.89 | 0.22 | 0.02 | -3.12 | 0.15 | sub-terminal/inferior | lateral | absent | heterocercal |
| *Carcharhinus limbatus* | fusiform | circular | 5.66 | 0.25 | 0.02 | -2.69 | 0.16 | sub-terminal/inferior | lateral | absent | heterocercal |
| *Carcharhinus longimanus* | fusiform | circular | 5.99 | 0.22 | 0.01 | -2.85 | 0.13 | sub-terminal/inferior | lateral | absent | heterocercal |
| *Carcharhinus perezii* | fusiform | circular | 5.70 | 0.23 | 0.03 | -2.64 | 0.15 | sub-terminal/inferior | lateral | absent | heterocercal |
| *Centropyge argi* | shortAndOrDeep | compressed | 2.08 | 0.21 | 0.10 | -3.43 | 0.45 | terminal | lateral | absent | homocercal |
| *Cephalopholis cruentata* | fusiform | oval | 3.75 | 0.34 | 0.06 | -2.42 | 0.29 | superior | lateral | absent | homocercal |
| *Cephalopholis fulva* | fusiform | oval | 3.78 | 0.36 | 0.06 | -2.11 | 0.29 | superior | lateral | absent | homocercal |
| *Chaetodon capistratus* | shortAndOrDeep | compressed | 2.71 | 0.24 | 0.08 | -2.46 | 0.53 | terminal | lateral | absent | homocercal |
| *Chaetodon sedentarius* | shortAndOrDeep | compressed | 2.71 | 0.24 | 0.09 | -2.68 | 0.51 | terminal | lateral | absent | homocercal |
| *Chaetodon striatus* | shortAndOrDeep | compressed | 2.77 | 0.23 | 0.07 | -2.55 | 0.56 | terminal | lateral | absent | homocercal |
| *Chilomycterus antennatus* | shortAndOrDeep | angular | 3.64 | 0.37 | 0.10 | -2.22 | 0.31 | terminal | lateral | absent | homocercal |
| *Clepticus parrae* | fusiform | oval | 3.40 | 0.26 | 0.05 | -2.67 | 0.26 | terminal | lateral | absent | homocercal |
| *Coryphopterus glaucofraenum* | elongated | oval | 2.08 | 0.21 | 0.06 | -2.87 | 0.16 | terminal | raised/top of head | absent | homocercal |
| *Dactylopterus volitans* | fusiform | angular | 4.09 | 0.17 | 0.05 | -2.99 | 0.15 | sub-terminal/inferior | raised/top of head | absent | homocercal |
| *Decapterus punctatus* | fusiform | compressed | 3.55 | 0.23 | 0.07 | -2.74 | 0.18 | terminal | lateral | absent | homocercal |
| *Diodon holocanthus* | shortAndOrDeep | angular | 3.91 | 0.27 | 0.07 | -3.31 | 0.44 | terminal | lateral | absent | homocercal |
| *Diodon hystrix* | shortAndOrDeep | angular | 4.51 | 0.26 | 0.07 | -3.19 | 0.35 | sub-terminal/inferior | lateral | absent | homocercal |
| *Diplodus caudimacula* | shortAndOrDeep | compressed | 3.40 | 0.21 | 0.07 | -2.90 | 0.45 | terminal | lateral | absent | homocercal |
| *Echidna catenata* | eel-like | compressed | 5.11 | 0.09 | 0.01 | -3.79 | 0.07 | terminal | lateral | absent | isocercal |
| *Elacatinus evelynae* | elongated | oval | 1.39 | 0.29 | 0.08 | -2.57 | 0.16 | sub-terminal/inferior | raised/top of head | absent | homocercal |

| | | | | | | | | | | | |
|---|---|---|---|---|---|---|---|---|---|---|---|
| *Entomacrodus nigricans* | elongated | oval | 2.30 | 0.17 | 0.05 | -3.29 | 0.20 | sub-terminal/inferior | raised/top of head | absent | homocercal |
| *Epinephelus guttatus* | fusiform | oval | 4.33 | 0.32 | 0.05 | -2.72 | 0.27 | superior | lateral | absent | homocercal |
| *Epinephelus itajara* | fusiform | compressed | 5.52 | 0.30 | 0.03 | -2.67 | 0.28 | superior | lateral | absent | homocercal |
| *Epinephelus striatus* | fusiform | compressed | 4.80 | 0.34 | 0.05 | -2.51 | 0.29 | superior | lateral | absent | homocercal |
| *Eques lanceolatus* | fusiform | compressed | 3.22 | 0.21 | 0.05 | -3.21 | 0.31 | sub-terminal/inferior | lateral | absent | homocercal |
| *Equetus punctatus* | fusiform | compressed | 3.30 | 0.21 | 0.06 | -3.12 | 0.29 | terminal | lateral | absent | homocercal |
| *Euthynnus alletteratus* | fusiform | oval | 4.80 | 0.22 | 0.05 | -2.95 | 0.23 | terminal | lateral | absent | homocercal |
| *Fistularia tabacaria* | elongated | circular | 5.30 | 0.25 | 0.02 | -1.75 | 0.03 | terminal | lateral | absent | homocercal |
| *Galeocerdo cuvier* | fusiform | angular | 6.62 | 0.17 | 0.01 | -3.19 | 0.14 | sub-terminal/inferior | lateral | present | heterocercal |
| *Gerres cinereus* | fusiform | compressed | 3.71 | 0.23 | 0.07 | -2.60 | 0.32 | terminal | lateral | absent | homocercal |
| *Ginglymostoma cirratum* | elongated | circular | 6.06 | 0.23 | 0.01 | -2.80 | 0.13 | sub-terminal/inferior | lateral | present | heterocercal |
| *Gnatholepis thompsoni* | elongated | compressed | 2.10 | 0.23 | 0.06 | -2.51 | 0.19 | terminal | raised/top of head | absent | homocercal |
| *Gobioclinus guppyi* | elongated | oval | 2.44 | 0.27 | 0.07 | -2.89 | 0.21 | terminal | raised/top of head | absent | homocercal |
| *Gramma loreto* | fusiform | compressed | 2.08 | 0.26 | 0.07 | -2.99 | 0.25 | terminal | raised/top of head | absent | homocercal |
| *Gramma melacara* | elongated | oval | 2.30 | 0.25 | 0.08 | -3.51 | 0.22 | terminal | lateral | absent | homocercal |
| *Gymnothorax moringa* | eel-like | compressed | 5.30 | 0.18 | 0.01 | -3.67 | 0.10 | terminal | lateral | absent | isocercal |
| *Gymnothorax vicinus* | eel-like | compressed | 4.88 | 0.14 | 0.01 | -3.76 | 0.06 | terminal | lateral | absent | isocercal |
| *Haemulon album* | fusiform | compressed | 4.37 | 0.29 | 0.05 | -2.26 | 0.31 | terminal | lateral | absent | homocercal |
| *Haemulon aurolineatum* | fusiform | compressed | 3.22 | 0.28 | 0.06 | -2.35 | 0.29 | terminal | lateral | absent | homocercal |
| *Haemulon carbonarium* | fusiform | compressed | 3.58 | 0.27 | 0.08 | -2.48 | 0.30 | terminal | lateral | absent | homocercal |
| *Haemulon flavolineatum* | shortAndOrDeep | compressed | 3.40 | 0.27 | 0.08 | -2.40 | 0.32 | terminal | lateral | absent | homocercal |
| *Haemulon macrostomum* | shortAndOrDeep | compressed | 3.76 | 0.32 | 0.06 | -1.98 | 0.35 | terminal | lateral | absent | homocercal |
| *Haemulon parra* | shortAndOrDeep | compressed | 3.72 | 0.28 | 0.07 | -2.38 | 0.32 | terminal | lateral | absent | homocercal |
| *Haemulon plumierii* | fusiform | compressed | 3.97 | 0.31 | 0.07 | -2.08 | 0.33 | terminal | lateral | absent | homocercal |
| *Haemulon sciurus* | fusiform | compressed | 3.83 | 0.28 | 0.06 | -2.17 | 0.32 | terminal | lateral | absent | homocercal |
| *Haemulon vittatum* | elongated | oval | 3.14 | 0.22 | 0.06 | -2.88 | 0.17 | terminal | lateral | absent | homocercal |
| *Halichoeres bivittatus* | fusiform | oval | 3.56 | 0.20 | 0.03 | -2.80 | 0.24 | terminal | lateral | absent | homocercal |
| *Halichoeres garnoti* | fusiform | oval | 2.96 | 0.25 | 0.06 | -2.63 | 0.22 | terminal | lateral | absent | homocercal |
| *Halichoeres maculipinna* | fusiform | compressed | 2.89 | 0.26 | 0.05 | -2.55 | 0.22 | sub-terminal/inferior | lateral | absent | homocercal |
| *Halichoeres poeyi* | fusiform | compressed | 3.00 | 0.23 | 0.06 | -2.62 | 0.23 | terminal | lateral | absent | homocercal |
| *Halichoeres radiatus* | fusiform | oval | 3.93 | 0.25 | 0.04 | -2.46 | 0.30 | terminal | lateral | absent | homocercal |
| *Harengula humeralis* | fusiform | compressed | 3.09 | 0.22 | 0.08 | -2.68 | 0.27 | terminal | lateral | absent | homocercal |
| *Hemiramphus balao* | elongated | compressed | 3.69 | 0.36 | 0.04 | -1.33 | 0.10 | superior | lateral | absent | hypocercal |
| *Heteroconger longissimus* | eel-like | circular | 3.93 | 0.08 | 0.01 | -4.91 | 0.03 | superior | raised/top of head | absent | reduced or absent |
| *Heteropriacanthus cruentatus* | fusiform | compressed | 3.93 | 0.26 | 0.11 | -2.76 | 0.33 | superior | lateral | absent | homocercal |
| *Holacanthus ciliaris* | shortAndOrDeep | compressed | 3.81 | 0.22 | 0.07 | -2.72 | 0.50 | terminal | lateral | absent | homocercal |
| *Holacanthus tricolor* | shortAndOrDeep | compressed | 3.56 | 0.20 | 0.05 | -2.62 | 0.45 | terminal | lateral | absent | homocercal |
| *Holocentrus adscensionis* | fusiform | compressed | 4.11 | 0.22 | 0.07 | -2.89 | 0.26 | terminal | lateral | absent | homocercal |
| *Holocentrus rufus* | fusiform | oval | 3.56 | 0.21 | 0.07 | -2.95 | 0.23 | terminal | lateral | absent | homocercal |

| | | | | | | | | | | | |
|---|---|---|---|---|---|---|---|---|---|---|---|
| *Hypanus americanus* | other | flattened | 6.03 | 0.24 | 0.03 | -2.22 | 0.06 | sub-terminal/inferior | raised/top of head | present | whip-like |
| *Hypoatherina harringtonensis* | elongated | oval | 2.30 | 0.20 | 0.06 | -2.84 | 0.15 | terminal | lateral | absent | homocercal |
| *Hypoplectrus aberrans* | shortAndOrDeep | compressed | 2.56 | 0.30 | 0.08 | -2.58 | 0.37 | terminal | lateral | absent | homocercal |
| *Hypoplectrus chlorurus* | shortAndOrDeep | compressed | 2.54 | 0.31 | 0.06 | -2.75 | 0.40 | terminal | lateral | absent | homocercal |
| *Hypoplectrus nigricans* | shortAndOrDeep | compressed | 2.72 | 0.32 | 0.08 | -2.06 | 0.32 | terminal | lateral | absent | homocercal |
| *Hypoplectrus puella* | shortAndOrDeep | compressed | 2.72 | 0.36 | 0.12 | -2.14 | 0.39 | terminal | lateral | absent | homocercal |
| *Kyphosus incisor* | fusiform | compressed | 4.65 | 0.21 | 0.05 | -2.87 | 0.32 | terminal | lateral | absent | homocercal |
| *Kyphosus sectatrix* | fusiform | compressed | 4.33 | 0.21 | 0.05 | -2.72 | 0.35 | terminal | lateral | absent | homocercal |
| *Labrisomus nuchipinnis* | fusiform | compressed | 3.14 | 0.25 | 0.04 | -2.65 | 0.26 | terminal | raised/top of head | absent | homocercal |
| *Lachnolaimus maximus* | shortAndOrDeep | compressed | 4.51 | 0.33 | 0.02 | -1.68 | 0.33 | sub-terminal/inferior | lateral | absent | homocercal |
| *Lactophrys bicaudalis* | shortAndOrDeep | compressed | 3.87 | 0.23 | 0.09 | -2.12 | 0.37 | sub-terminal/inferior | lateral | absent | homocercal |
| *Lactophrys trigonus* | shortAndOrDeep | angular | 4.01 | 0.28 | 0.07 | -2.05 | 0.42 | sub-terminal/inferior | raised/top of head | absent | homocercal |
| *Lactophrys triqueter* | shortAndOrDeep | angular | 3.85 | 0.23 | 0.08 | -2.02 | 0.44 | sub-terminal/inferior | raised/top of head | absent | homocercal |
| *Lutjanus apodus* | fusiform | oval | 4.43 | 0.31 | 0.07 | -2.15 | 0.30 | terminal | lateral | absent | homocercal |
| *Lutjanus cyanopterus* | fusiform | oval | 5.08 | 0.30 | 0.05 | -2.26 | 0.32 | terminal | lateral | absent | homocercal |
| *Lutjanus griseus* | fusiform | oval | 4.49 | 0.30 | 0.07 | -2.20 | 0.30 | terminal | lateral | absent | homocercal |
| *Lutjanus mahogoni* | fusiform | oval | 3.87 | 0.31 | 0.08 | -2.21 | 0.28 | terminal | lateral | absent | homocercal |
| *Malacanthus plumieri* | elongated | oval | 4.38 | 0.21 | 0.04 | -2.31 | 0.15 | terminal | lateral | absent | homocercal |
| *Megalops atlanticus* | fusiform | compressed | 5.52 | 0.17 | 0.03 | -3.55 | 0.24 | superior | lateral | absent | homocercal |
| *Melichthys niger* | shortAndOrDeep | compressed | 3.91 | 0.22 | 0.04 | -2.00 | 0.41 | terminal | lateral | absent | homocercal |
| *Microspathodon chrysurus* | shortAndOrDeep | compressed | 3.04 | 0.26 | 0.08 | -2.84 | 0.41 | terminal | lateral | absent | homocercal |
| *Monacanthus ciliatus* | shortAndOrDeep | compressed | 3.00 | 0.24 | 0.07 | -1.72 | 0.54 | superior | lateral | absent | homocercal |
| *Mulloidichthys martinicus* | fusiform | oval | 3.80 | 0.21 | 0.06 | -2.54 | 0.22 | sub-terminal/inferior | lateral | absent | homocercal |
| *Mustelus canis* | elongated | circular | 5.01 | 0.19 | 0.03 | -2.73 | 0.13 | sub-terminal/inferior | lateral | present | heterocercal |
| *Mycteroperca tigris* | fusiform | oval | 4.62 | 0.32 | 0.04 | -2.55 | 0.25 | superior | lateral | absent | homocercal |
| *Mycteroperca venenosa* | fusiform | oval | 4.61 | 0.29 | 0.04 | -2.56 | 0.27 | superior | lateral | absent | homocercal |
| *Myrichthys breviceps* | eel-like | circular | 4.62 | 0.06 | 0.01 | -4.76 | 0.03 | sub-terminal/inferior | lateral | absent | isocercal |
| *Myrichthys ocellatus* | eel-like | circular | 4.70 | 0.06 | 0.01 | -4.76 | 0.03 | sub-terminal/inferior | lateral | absent | isocercal |
| *Myripristis jacobus* | fusiform | oval | 3.22 | 0.27 | 0.12 | -3.66 | 0.36 | superior | lateral | absent | homocercal |
| *Negaprion brevirostris* | fusiform | circular | 5.83 | 0.22 | 0.02 | -2.88 | 0.14 | sub-terminal/inferior | lateral | absent | heterocercal |
| *Neoniphon marianus* | fusiform | compressed | 2.89 | 0.32 | 0.11 | -2.31 | 0.25 | terminal | lateral | absent | homocercal |
| *Neoniphon vexillarium* | fusiform | oval | 2.89 | 0.24 | 0.10 | -3.17 | 0.29 | terminal | lateral | absent | homocercal |
| *Ocyurus chrysurus* | fusiform | oval | 4.46 | 0.27 | 0.06 | -2.50 | 0.30 | terminal | lateral | absent | homocercal |
| *Odontoscion dentex* | fusiform | compressed | 3.40 | 0.26 | 0.07 | -3.08 | 0.28 | terminal | lateral | absent | homocercal |
| *Ogcocephalus nasutus* | other | angular | 3.64 | 0.23 | 0.02 | -3.90 | 0.24 | terminal | lateral | absent | homocercal |
| *Ophichthus ophis* | eel-like | oval | 5.35 | 0.11 | 0.01 | -3.81 | 0.05 | sub-terminal/inferior | lateral | absent | isocercal |
| *Ophioblennius atlanticus* | fusiform | oval | 2.94 | 0.18 | 0.03 | -2.89 | 0.19 | sub-terminal/inferior | raised/top of head | absent | homocercal |
| *Opistognathus aurifrons* | elongated | oval | 2.30 | 0.23 | 0.05 | -3.33 | 0.14 | terminal | raised/top of head | absent | homocercal |
| *Opistognathus maxillosus* | elongated | oval | 2.56 | 0.29 | 0.05 | -3.30 | 0.20 | terminal | raised/top of head | absent | homocercal |

| | | | | | | | | | | | |
|---|---|---|---|---|---|---|---|---|---|---|---|
| *Opistognathus whitehursti* | elongated | oval | 2.64 | 0.30 | 0.10 | -3.21 | 0.17 | terminal | raised/top of head | absent | homocercal |
| *Parablennius marmoreus* | fusiform | oval | 2.14 | 0.24 | 0.06 | -3.69 | 0.20 | sub-terminal/inferior | raised/top of head | absent | homocercal |
| *Paranthias furcifer* | fusiform | oval | 3.65 | 0.23 | 0.04 | -2.73 | 0.25 | superior | lateral | absent | homocercal |
| *Pareques acuminatus* | fusiform | compressed | 3.14 | 0.25 | 0.06 | -3.19 | 0.30 | sub-terminal/inferior | lateral | absent | homocercal |
| *Pempheris schomburgkii* | shortAndOrDeep | compressed | 2.71 | 0.24 | 0.11 | -3.51 | 0.35 | superior | lateral | absent | homocercal |
| *Phaeoptyx conklini* | fusiform | oval | 2.48 | 0.28 | 0.11 | -3.41 | 0.29 | terminal | lateral | absent | homocercal |
| *Platybelone argalus* | eel-like | circular | 3.91 | 0.36 | 0.04 | -1.38 | 0.05 | terminal | lateral | absent | homocercal |
| *Pomacanthus arcuatus* | shortAndOrDeep | compressed | 4.09 | 0.23 | 0.08 | -2.78 | 0.59 | terminal | lateral | absent | homocercal |
| *Pomacanthus paru* | shortAndOrDeep | compressed | 3.72 | 0.27 | 0.04 | -2.08 | 0.64 | terminal | lateral | absent | homocercal |
| *Priacanthus arenatus* | fusiform | compressed | 3.91 | 0.25 | 0.11 | -2.94 | 0.29 | superior | lateral | absent | homocercal |
| *Prognathodes aculeatus* | shortAndOrDeep | compressed | 2.30 | 0.33 | 0.08 | -1.90 | 0.49 | terminal | lateral | absent | homocercal |
| *Pseudupeneus maculatus* | fusiform | oval | 3.40 | 0.22 | 0.05 | -2.30 | 0.24 | terminal | lateral | absent | homocercal |
| *Rhizoprionodon porosus* | fusiform | circular | 4.73 | 0.22 | 0.02 | -2.66 | 0.13 | sub-terminal/inferior | lateral | absent | heterocercal |
| *Rypticus saponaceus* | fusiform | compressed | 3.56 | 0.29 | 0.05 | -2.73 | 0.27 | superior | lateral | absent | homocercal |
| *Sargocentron coruscum* | fusiform | oval | 2.71 | 0.27 | 0.11 | -2.92 | 0.27 | terminal | lateral | absent | homocercal |
| *Scartella cristata* | fusiform | circular | 2.48 | 0.19 | 0.03 | -3.04 | 0.21 | sub-terminal/inferior | raised/top of head | absent | homocercal |
| *Scarus coelestinus* | fusiform | compressed | 4.34 | 0.25 | 0.03 | -2.24 | 0.33 | terminal | lateral | absent | homocercal |
| *Scarus guacamaia* | fusiform | oval | 4.79 | 0.27 | 0.04 | -2.33 | 0.32 | sub-terminal/inferior | lateral | absent | homocercal |
| *Scarus iseri* | fusiform | oval | 3.56 | 0.25 | 0.03 | -2.48 | 0.32 | sub-terminal/inferior | lateral | absent | homocercal |
| *Scarus taeniopterus* | fusiform | oval | 3.56 | 0.26 | 0.05 | -2.22 | 0.30 | sub-terminal/inferior | lateral | absent | homocercal |
| *Scarus vetula* | fusiform | compressed | 4.11 | 0.25 | 0.03 | -2.34 | 0.31 | sub-terminal/inferior | lateral | absent | homocercal |
| *Scomberomorus cavalla* | fusiform | oval | 5.21 | 0.19 | 0.03 | -2.71 | 0.18 | terminal | lateral | absent | homocercal |
| *Scomberomorus regalis* | fusiform | oval | 5.21 | 0.19 | 0.04 | -2.57 | 0.15 | terminal | lateral | absent | homocercal |
| *Scorpaena brasiliensis* | fusiform | compressed | 3.56 | 0.30 | 0.08 | -2.81 | 0.26 | terminal | raised/top of head | absent | homocercal |
| *Scorpaena grandicornis* | fusiform | oval | 3.40 | 0.33 | 0.07 | -2.56 | 0.28 | terminal | raised/top of head | absent | homocercal |
| *Scorpaena inermis* | fusiform | oval | 2.60 | 0.36 | 0.08 | -2.19 | 0.24 | terminal | raised/top of head | absent | homocercal |
| *Scorpaena plumieri* | fusiform | compressed | 3.81 | 0.32 | 0.06 | -2.56 | 0.30 | terminal | raised/top of head | absent | homocercal |
| *Scorpaenodes caribbaeus* | fusiform | oval | 2.48 | 0.35 | 0.09 | -2.36 | 0.26 | terminal | raised/top of head | absent | homocercal |
| *Selar crumenophthalmus* | fusiform | compressed | 4.25 | 0.24 | 0.08 | -2.78 | 0.24 | terminal | lateral | absent | homocercal |
| *Seriola dumerili* | fusiform | oval | 5.25 | 0.23 | 0.05 | -2.67 | 0.26 | terminal | lateral | absent | homocercal |
| *Serranus tigrinus* | fusiform | oval | 3.40 | 0.32 | 0.07 | -2.50 | 0.21 | terminal | lateral | absent | homocercal |
| *Sparisoma aurofrenatum* | fusiform | compressed | 3.33 | 0.24 | 0.05 | -2.39 | 0.34 | terminal | lateral | absent | homocercal |
| *Sparisoma chrysopterum* | fusiform | compressed | 3.83 | 0.25 | 0.05 | -2.51 | 0.31 | sub-terminal/inferior | lateral | absent | homocercal |
| *Sparisoma rubripinne* | fusiform | compressed | 3.87 | 0.26 | 0.04 | -2.48 | 0.32 | terminal | lateral | absent | homocercal |
| *Sparisoma viride* | fusiform | oval | 4.16 | 0.26 | 0.04 | -2.21 | 0.33 | terminal | lateral | absent | homocercal |
| *Sphoeroides spengleri* | fusiform | angular | 3.40 | 0.24 | 0.06 | -2.14 | 0.28 | terminal | raised/top of head | absent | homocercal |
| *Sphyraena picudilla* | elongated | circular | 4.18 | 0.28 | 0.05 | -2.22 | 0.11 | terminal | lateral | absent | homocercal |
| *Sphyrna lewini* | elongated | circular | 6.06 | 0.23 | 0.02 | -3.61 | 0.12 | sub-terminal/inferior | lateral | absent | heterocercal |
| *Sphyrna tiburo* | elongated | circular | 5.01 | 0.22 | 0.02 | -2.60 | 0.14 | sub-terminal/inferior | lateral | absent | heterocercal |

| | | | | | | | | | | | |
|---|---|---|---|---|---|---|---|---|---|---|---|
| *Stegastes fuscus* | fusiform | compressed | 2.53 | 0.22 | 0.06 | -2.90 | 0.41 | terminal | lateral | absent | homocercal |
| *Stegastes leucostictus* | shortAndOrDeep | compressed | 2.30 | 0.23 | 0.08 | -3.21 | 0.43 | terminal | lateral | absent | homocercal |
| *Stegastes planifrons* | shortAndOrDeep | compressed | 2.56 | 0.24 | 0.08 | -2.85 | 0.44 | terminal | lateral | absent | homocercal |
| *Stegastes variabilis* | shortAndOrDeep | compressed | 2.53 | 0.27 | 0.07 | -2.46 | 0.39 | terminal | lateral | absent | homocercal |
| *Strongylura timucu* | elongated | circular | 4.11 | 0.34 | 0.03 | -1.50 | 0.05 | terminal | lateral | absent | homocercal |
| *Synodus intermedius* | elongated | circular | 3.83 | 0.23 | 0.03 | -2.78 | 0.14 | terminal | raised/top of head | absent | homocercal |
| *Synodus synodus* | elongated | circular | 3.76 | 0.26 | 0.04 | -2.70 | 0.13 | terminal | raised/top of head | absent | homocercal |
| *Thalassoma bifasciatum* | fusiform | oval | 3.22 | 0.22 | 0.03 | -2.60 | 0.19 | terminal | lateral | absent | homocercal |
| *Trachinotus falcatus* | shortAndOrDeep | compressed | 5.01 | 0.20 | 0.05 | -3.06 | 0.41 | terminal | lateral | absent | homocercal |
| *Trachinotus goodei* | shortAndOrDeep | compressed | 3.91 | 0.17 | 0.04 | -3.47 | 0.40 | terminal | lateral | absent | homocercal |
| *Tylosurus crocodilus* | elongated | circular | 5.01 | 0.25 | 0.02 | -1.90 | 0.08 | terminal | lateral | absent | homocercal |
| *Xyrichtys novacula* | elongated | compressed | 3.64 | 0.21 | 0.03 | -2.63 | 0.31 | terminal | lateral | absent | homocercal |
| *Xyrichtys splendens* | elongated | oval | 2.86 | 0.26 | 0.05 | -2.39 | 0.29 | terminal | lateral | absent | homocercal |
| *Acanthistius pictus* | shortAndOrDeep | compressed | 3.85 | 0.29 | 0.04 | -2.64 | 0.36 | terminal | lateral | absent | homocercal |
| *Aplodactylus punctatus* | elongated | oval | 3.78 | 0.21 | 0.03 | -2.71 | 0.27 | sub-terminal/inferior | lateral | absent | homocercal |
| *Auchenionchus microcirrhis* | elongated | compressed | 3.10 | 0.22 | 0.05 | -2.90 | 0.20 | terminal | raised/top of head | absent | homocercal |
| *Bovichtus chilensis* | elongated | compressed | 2.24 | 0.30 | 0.06 | -2.80 | 0.17 | terminal | raised/top of head | absent | homocercal |
| *Calliclinus geniguttatus* | elongated | compressed | 2.50 | 0.28 | 0.06 | -2.74 | 0.19 | terminal | raised/top of head | absent | homocercal |
| *Cheilodactylus variegatus* | fusiform | oval | 3.78 | 0.27 | 0.05 | -2.40 | 0.28 | terminal | lateral | absent | homocercal |
| *Chromis crusma* | shortAndOrDeep | oval | 2.64 | 0.23 | 0.06 | -3.22 | 0.36 | superior | lateral | absent | homocercal |
| *Eptatretus polytrema* | eel-like | circular | 4.53 | 0.30 | 0.00 | -2.82 | 0.09 | sub-terminal/inferior | lateral | absent | protocercal |
| *Genypterus chilensis* | elongated | oval | 5.01 | 0.23 | 0.02 | -2.92 | 0.11 | terminal | lateral | absent | isocercal |
| *Girella laevifrons* | shortAndOrDeep | oval | 3.65 | 0.27 | 0.08 | -2.70 | 0.31 | terminal | lateral | absent | homocercal |
| *Gobiesox marmoratus* | elongated | circular | 2.19 | 0.25 | 0.06 | -2.67 | 0.18 | superior | raised/top of head | absent | homocercal |
| *Graus nigra* | shortAndOrDeep | compressed | 4.17 | 0.31 | 0.04 | -2.26 | 0.31 | terminal | lateral | absent | homocercal |
| *Helcogrammoides cunninghami* | elongated | compressed | 1.82 | 0.23 | 0.06 | -2.97 | 0.18 | terminal | raised/top of head | absent | homocercal |
| *Hemilutjanus macrophthalmos* | fusiform | oval | 3.91 | 0.30 | 0.07 | -2.77 | 0.34 | terminal | lateral | absent | homocercal |
| *Hypsoblennius sordidus* | fusiform | compressed | 2.94 | 0.18 | 0.05 | -3.33 | 0.14 | terminal | raised/top of head | absent | homocercal |
| *Isacia conceptionis* | fusiform | oval | 4.09 | 0.24 | 0.05 | -2.85 | 0.26 | terminal | lateral | absent | homocercal |
| *Labrisomus philippii* | fusiform | compressed | 3.61 | 0.26 | 0.05 | -2.69 | 0.27 | terminal | raised/top of head | absent | homocercal |
| *Myxodes viridis* | elongated | compressed | 2.60 | 0.17 | 0.04 | -3.39 | 0.19 | terminal | lateral | absent | homocercal |
| *Paralabrax humeralis* | fusiform | oval | 4.00 | 0.30 | 0.06 | -2.35 | 0.26 | terminal | lateral | absent | homocercal |
| *Paralichthys adspersus* | shortAndOrDeep | compressed and lies on side | 4.25 | 0.25 | 0.03 | -3.27 | 0.44 | superior | migrated to one side and raised | absent | homocercal |
| *Paralichthys microps* | shortAndOrDeep | compressed and lies on side | 3.71 | 0.26 | 0.02 | -2.99 | 0.38 | superior | migrated to one side and raised | absent | homocercal |
| *Pinguipes chilensis* | elongated | oval | 3.93 | 0.23 | 0.05 | -2.45 | 0.17 | terminal | lateral | absent | homocercal |
| *Scartichthys viridis* | elongated | oval | 2.98 | 0.17 | 0.05 | -2.99 | 0.25 | sub-terminal/inferior | raised/top of head | absent | homocercal |
| *Schroederichthys chilensis* | elongated | circular | 4.13 | 0.19 | 0.02 | -3.17 | 0.12 | sub-terminal/inferior | lateral | present | heterocercal |

| | | | | | | | | | | | |
|---|---|---|---|---|---|---|---|---|---|---|---|
| *Sebastes oculatus* | fusiform | compressed | 3.71 | 0.33 | 0.06 | -2.29 | 0.27 | terminal | raised/top of head | absent | homocercal |
| *Semicossyphus darwini* | fusiform | compressed | 4.25 | 0.25 | 0.04 | -2.54 | 0.26 | terminal | lateral | absent | homocercal |
| *Sicyases sanguineus* | elongated | circular | 2.13 | 0.31 | 0.03 | -2.76 | 0.22 | sub-terminal/inferior | lateral | absent | homocercal |
| *Ambassis agassizii* | fusiform | compressed | 2.30 | 0.23 | 0.09 | -2.90 | 0.29 | terminal | lateral | absent | homocercal |
| *Anguilla australis* | eel-like | circular | 4.87 | 0.14 | 0.01 | -3.61 | 0.07 | terminal | lateral | absent | isocercal |
| *Anguilla reinhardtii* | eel-like | circular | 5.11 | 0.14 | 0.01 | -3.48 | 0.07 | terminal | lateral | absent | isocercal |
| *Galaxias maculatus* | elongated | compressed | 3.07 | 0.17 | 0.04 | -3.04 | 0.15 | terminal | lateral | absent | homocercal |
| *Gobiomorphus australis* | elongated | compressed | 2.89 | 0.21 | 0.05 | -2.95 | 0.18 | superior | lateral | absent | homocercal |
| *Gobiomorphus coxii* | elongated | compressed | 2.94 | 0.24 | 0.05 | -2.78 | 0.18 | superior | lateral | absent | homocercal |
| *Hypseleotris compressa* | fusiform | compressed | 2.48 | 0.22 | 0.04 | -2.73 | 0.29 | superior | lateral | absent | homocercal |
| *Hypseleotris galii* | fusiform | compressed | 1.70 | 0.25 | 0.07 | -2.75 | 0.21 | superior | lateral | absent | homocercal |
| *Leiopotherapon unicolor* | fusiform | compressed | 3.50 | 0.24 | 0.06 | -3.00 | 0.25 | terminal | lateral | absent | homocercal |
| *Mugil cephalus* | fusiform | circular | 4.78 | 0.18 | 0.04 | -3.37 | 0.20 | terminal | lateral | absent | homocercal |
| *Notesthes robusta* | shortAndOrDeep | compressed | 3.56 | 0.29 | 0.05 | -2.56 | 0.28 | superior | raised/top of head | absent | homocercal |
| *Percalates novemaculeatus* | fusiform | compressed | 4.09 | 0.24 | 0.07 | -3.08 | 0.26 | terminal | lateral | absent | homocercal |
| *Philypnodon grandiceps* | elongated | compressed | 2.48 | 0.26 | 0.04 | -2.63 | 0.14 | superior | raised/top of head | absent | homocercal |
| *Philypnodon macrostomus* | fusiform | compressed | 1.87 | 0.29 | 0.06 | -2.68 | 0.14 | superior | raised/top of head | absent | homocercal |
| *Potamalosa richmondia* | fusiform | compressed | 3.63 | 0.17 | 0.05 | -3.38 | 0.20 | terminal | lateral | absent | homocercal |
| *Pseudomugil signifer* | fusiform | compressed | 2.22 | 0.19 | 0.07 | -2.90 | 0.22 | superior | lateral | absent | homocercal |
| *Retropinna semoni* | elongated | compressed | 2.30 | 0.17 | 0.06 | -3.17 | 0.14 | terminal | lateral | absent | homocercal |
| *Tandanus tandanus* | fusiform | circular | 4.50 | 0.19 | 0.04 | -2.78 | 0.17 | sub-terminal/inferior | lateral | absent | isocercal |
| *Trachystoma petardi* | fusiform | oval | 4.39 | 0.22 | 0.05 | -2.85 | 0.19 | terminal | lateral | absent | homocercal |
| *Agonus cataphractus* | fusiform | angular | 3.04 | 0.22 | 0.05 | -2.77 | 0.14 | sub-terminal/inferior | raised/top of head | absent | homocercal |
| *Ammodytes tobianus* | elongated | circular | 3.09 | 0.16 | 0.02 | -2.94 | 0.10 | superior | lateral | absent | homocercal |
| *Anguilla anguilla* | eel-like | circular | 4.80 | 0.13 | 0.02 | -3.44 | 0.06 | terminal | lateral | absent | isocercal |
| *Ciliata mustela* | elongated | oval | 3.22 | 0.19 | 0.02 | -3.14 | 0.13 | sub-terminal/inferior | lateral | absent | homocercal |
| *Clupea harengus* | fusiform | oval | 3.97 | 0.18 | 0.05 | -3.00 | 0.19 | superior | lateral | absent | homocercal |
| *Gasterosteus aculeatus* | fusiform | compressed | 2.40 | 0.25 | 0.06 | -2.48 | 0.23 | terminal | lateral | absent | homocercal |
| *Myoxocephalus scorpius* | fusiform | oval | 4.09 | 0.33 | 0.06 | -2.41 | 0.23 | sub-terminal/inferior | raised/top of head | absent | homocercal |
| *Pholis gunnellus* | eel-like | compressed | 3.29 | 0.10 | 0.02 | -3.94 | 0.12 | terminal | lateral | absent | homocercal |
| *Platichthys flesus* | shortAndOrDeep | compressed and lies on side | 4.09 | 0.21 | 0.04 | -3.81 | 0.38 | superior | migrated to one side and raised | absent | homocercal |
| *Pleuronectes platessa* | shortAndOrDeep | compressed and lies on side | 4.82 | 0.23 | 0.02 | -2.87 | 0.42 | superior | migrated to one side and raised | absent | homocercal |
| *Pollachius virens* | fusiform | circular | 4.87 | 0.22 | 0.04 | -2.84 | 0.20 | terminal | lateral | absent | homocercal |
| *Pomatoschistus microps* | elongated | compressed | 2.20 | 0.25 | 0.05 | -3.04 | 0.17 | superior | raised/top of head | absent | homocercal |
| *Pomatoschistus minutus* | elongated | compressed | 2.40 | 0.23 | 0.05 | -2.91 | 0.13 | superior | raised/top of head | absent | homocercal |
| *Salmo trutta* | fusiform | oval | 5.06 | 0.21 | 0.05 | -3.01 | 0.21 | terminal | lateral | absent | homocercal |
| *Sprattus sprattus* | fusiform | compressed | 2.93 | 0.19 | 0.04 | -2.73 | 0.19 | superior | lateral | absent | homocercal |
| *Syngnathus rostellatus* | eel-like | angular | 2.92 | 0.14 | 0.02 | -2.64 | 0.04 | terminal | lateral | absent | homocercal |
| *Zoarces viviparus* | elongated | oval | 3.95 | 0.19 | 0.03 | -3.27 | 0.16 | sub-terminal/inferior | raised/top of head | absent | isocercal |

| | | | | | | | | | | | |
|---|---|---|---|---|---|---|---|---|---|---|---|
| *Adololopas moyasmithae* | shortAndOrDeep | circular | 3.39 | 0.24 | 0.03 | -2.74 | 0.19 | sub-terminal/inferior | lateral | absent | heterocercal |
| *Asthenorhynchus meemannae* | fusiform | compressed | 3.91 | 0.26 | 0.03 | -2.41 | 0.18 | sub-terminal/inferior | lateral | absent | heterocercal |
| *Austroptyctodus gardineri* | elongated | compressed | 3.00 | 0.10 | 0.06 | -2.38 | 0.25 | terminal | lateral | present | whip-like |
| *Bothriolepis canadensis* | shortAndOrDeep | angular | 4.51 | 0.12 | 0.03 | -3.48 | 0.16 | sub-terminal/inferior | raised/top of head | absent | heterocercal |
| *Bothriolepis sp.* | shortAndOrDeep | angular | 3.50 | 0.12 | 0.03 | -2.80 | 0.15 | sub-terminal/inferior | raised/top of head | absent | heterocercal |
| *Bothriolepis yeungae* | fusiform | angular | 3.91 | 0.10 | 0.01 | -3.22 | 0.09 | sub-terminal/inferior | raised/top of head | absent | heterocercal |
| *Bruntonichthys multidens* | fusiform | circular | 3.78 | 0.18 | 0.05 | -3.78 | 0.19 | terminal | lateral | present | heterocercal |
| *Bullerichthys fascidens* | fusiform | circular | 3.90 | 0.22 | 0.03 | -3.43 | 0.19 | terminal | lateral | present | heterocercal |
| *Cabonnichthys burnsi* | elongated | circular | 4.32 | 0.16 | 0.01 | -3.31 | 0.12 | terminal | lateral | present | isocercal |
| *Callistiopterus clappi* | fusiform | circular | 2.05 | 0.24 | 0.03 | -2.75 | 0.14 | terminal | lateral | present | isocercal |
| *Campbellodus decipiens* | elongated | compressed | 3.91 | 0.22 | 0.03 | -3.02 | 0.18 | terminal | lateral | present | whip-like |
| *Camuropiscis concinnus* | fusiform | circular | 3.37 | 0.21 | 0.04 | -2.84 | 0.13 | sub-terminal/inferior | lateral | absent | heterocercal |
| *Camuropiscis laidlawi* | shortAndOrDeep | angular | 3.09 | 0.25 | 0.04 | -2.17 | 0.19 | sub-terminal/inferior | lateral | absent | heterocercal |
| *Canowindra grossi* | elongated | circular | 3.91 | 0.21 | 0.01 | -2.66 | 0.09 | terminal | lateral | present | isocercal |
| *Cheirolepis canadensis* | fusiform | oval | 3.97 | 0.24 | 0.03 | -3.56 | 0.18 | terminal | lateral | present | heterocercal |
| *Chirodipterus australis* | shortAndOrDeep | circular | 3.36 | 0.21 | 0.03 | -2.76 | 0.21 | sub-terminal/inferior | lateral | absent | heterocercal |
| *Compagopiscis croucheri* | fusiform | circular | 3.40 | 0.21 | 0.05 | -3.91 | 0.20 | sub-terminal/inferior | lateral | absent | heterocercal |
| *Diplacanthus ellsi* | fusiform | oval | 2.56 | 0.17 | 0.03 | -4.17 | 0.17 | terminal | lateral | absent | heterocercal |
| *Diplacanthus horridus* | fusiform | oval | 3.01 | 0.22 | 0.05 | -3.52 | 0.23 | terminal | lateral | absent | heterocercal |
| *Eastmanosteus calliaspis* | fusiform | angular | 4.32 | 0.18 | 0.04 | -3.53 | 0.20 | terminal | lateral | present | heterocercal |
| *Eastmanosteus sp. A* | fusiform | angular | 4.38 | 0.18 | 0.03 | -3.51 | 0.22 | terminal | lateral | present | heterocercal |
| *Eastmanosteus sp. B* | fusiform | angular | 4.94 | 0.19 | 0.02 | -3.91 | 0.12 | terminal | lateral | present | heterocercal |
| *Elpistostege watsoni* | eel-like | flattened | 5.06 | 0.22 | 0.01 | -2.49 | 0.08 | terminal | raised/top of head | present | isocercal |
| *Escuminaspis laticeps* | other | flattened | 3.77 | 0.39 | 0.03 | -1.98 | 0.12 | sub-terminal/inferior | raised/top of head | absent | heterocercal |
| *Euphanerops longaevus* | eel-like | circular | 3.64 | 0.17 | 0.04 | -3.23 | 0.14 | sub-terminal/inferior | lateral | absent | hypocercal |
| *Eusthenopteron foordi* | fusiform | circular | 4.77 | 0.22 | 0.02 | -3.31 | 0.18 | terminal | lateral | present | isocercal |
| *Fallacosteus turnerae* | fusiform | circular | 3.40 | 0.22 | 0.05 | -2.61 | 0.13 | sub-terminal/inferior | lateral | absent | heterocercal |
| *Fleurantia denticulata* | fusiform | compressed | 4.08 | 0.26 | 0.05 | -2.19 | 0.22 | terminal | lateral | present | heterocercal |
| *Glyptolepis sp.* | fusiform | compressed | 4.07 | 0.20 | 0.01 | -3.60 | 0.18 | terminal | lateral | present | heterocercal |
| *Gogodipterus paddyensis* | shortAndOrDeep | circular | 3.89 | 0.22 | 0.03 | -2.77 | 0.18 | sub-terminal/inferior | lateral | absent | heterocercal |
| *Gogonasus andrewsae* | fusiform | circular | 3.59 | 0.17 | 0.01 | -3.90 | 0.19 | sub-terminal/inferior | lateral | present | heterocercal |
| *Gogosardina coatesi* | fusiform | oval | 2.60 | 0.22 | 0.07 | -3.52 | 0.17 | sub-terminal/inferior | lateral | present | heterocercal |
| *Gogoselachus lynnbeazleyae* | elongated | circular | 3.40 | 0.23 | 0.05 | -2.84 | 0.21 | terminal | lateral | present | heterocercal |
| *Gooloogongia loomesi* | eel-like | flattened | 4.38 | 0.18 | 0.02 | -2.86 | 0.18 | superior | raised/top of head | present | isocercal |
| *Griphognathus whitei* | fusiform | compressed | 4.20 | 0.29 | 0.03 | -1.91 | 0.29 | sub-terminal/inferior | lateral | absent | heterocercal |
| *Groenlandaspis sp.* | shortAndOrDeep | angular | 3.83 | 0.22 | 0.02 | -3.14 | 0.33 | sub-terminal/inferior | raised/top of head | present | heterocercal |
| *Halimacanthodes ahlbergi* | elongated | compressed | 3.18 | 0.25 | 0.04 | -2.08 | 0.22 | terminal | lateral | present | heterocercal |
| *Harrytoombsia elegans* | fusiform | circular | 3.33 | 0.21 | 0.06 | -3.07 | 0.26 | terminal | lateral | present | heterocercal |

| | | | | | | | | | | | |
|---|---|---|---|---|---|---|---|---|---|---|---|
| *Holodipterus elderae* | elongated | compressed | 3.91 | 0.24 | 0.03 | -2.58 | 0.18 | sub-terminal/inferior | lateral | absent | heterocercal |
| *Holodipterus gogoensis* | fusiform | compressed | 4.27 | 0.25 | 0.03 | -2.26 | 0.24 | sub-terminal/inferior | lateral | absent | heterocercal |
| *Holonema westolli* | fusiform | compressed | 4.27 | 0.25 | 0.02 | -2.83 | 0.22 | sub-terminal/inferior | raised/top of head | present | heterocercal |
| *Holoptychius jarviki* | elongated | oval | 3.85 | 0.20 | 0.02 | -3.59 | 0.18 | terminal | lateral | present | heterocercal |
| *Homalacanthus concinnus* | elongated | oval | 3.43 | 0.15 | 0.03 | -4.35 | 0.20 | terminal | lateral | absent | heterocercal |
| *Incisoscutum ritchiei* | fusiform | circular | 3.37 | 0.20 | 0.06 | -3.27 | 0.17 | sub-terminal/inferior | lateral | absent | heterocercal |
| *Incisoscutum sarahae* | fusiform | circular | 3.91 | 0.18 | 0.04 | -3.38 | 0.16 | sub-terminal/inferior | lateral | present | heterocercal |
| *Kapitany sarcopt* | fusiform | compressed | 3.90 | 0.23 | 0.03 | -2.99 | 0.18 | terminal | lateral | present | heterocercal |
| *Kendrickichthys cavernosus* | fusiform | circular | 4.64 | 0.17 | 0.03 | -3.55 | 0.16 | terminal | lateral | present | heterocercal |
| *Kimberleyichthys bispicatus* | fusiform | circular | 4.38 | 0.24 | 0.03 | -3.73 | 0.21 | terminal | lateral | present | heterocercal |
| *Kimberleyichthys whybrowi* | fusiform | circular | 4.29 | 0.17 | 0.03 | -4.19 | 0.21 | terminal | lateral | present | heterocercal |
| *Latocamurus coulthardi* | fusiform | circular | 3.14 | 0.20 | 0.06 | -3.49 | 0.14 | sub-terminal/inferior | lateral | absent | heterocercal |
| *Levesquaspis patteni* | other | flattened | 2.83 | 0.40 | 0.06 | -2.13 | 0.12 | sub-terminal/inferior | raised/top of head | absent | heterocercal |
| *Mandageria fairfaxi* | elongated | circular | 5.08 | 0.17 | 0.01 | -3.24 | 0.12 | sub-terminal/inferior | lateral | present | isocercal |
| *Materpiscis attenboroughi* | elongated | compressed | 3.74 | 0.22 | 0.03 | -2.90 | 0.16 | terminal | lateral | present | whip-like |
| *Mcnamaraspis kaprios* | fusiform | circular | 3.57 | 0.16 | 0.04 | -3.79 | 0.18 | sub-terminal/inferior | lateral | present | heterocercal |
| *Miguashaia bureaui* | fusiform | compressed | 3.88 | 0.27 | 0.04 | -3.23 | 0.22 | terminal | lateral | present | heterocercal |
| *Mimipiscis bartrami* | fusiform | oval | 2.53 | 0.24 | 0.07 | -3.44 | 0.26 | sub-terminal/inferior | lateral | present | homocercal |
| *Mimipiscis toombsi (normal)* | fusiform | oval | 2.08 | 0.31 | 0.08 | -3.39 | 0.25 | sub-terminal/inferior | lateral | present | homocercal |
| *Mimipiscis toombsi (supersized)* | fusiform | oval | 3.00 | 0.30 | 0.05 | -3.10 | 0.33 | sub-terminal/inferior | lateral | present | homocercal |
| *Moythomasia durgaringa* | fusiform | oval | 2.83 | 0.24 | 0.04 | -3.19 | 0.28 | sub-terminal/inferior | lateral | present | heterocercal |
| *Namugawi wirngarri* | fusiform | oval | 3.11 | 0.14 | 0.02 | -3.62 | 0.11 | terminal | lateral | present | leptocercal |
| *Onychodus jandemarrai* | fusiform | compressed | 5.08 | 0.20 | 0.03 | -3.47 | 0.10 | terminal | lateral | present | leptocercal |
| *Pickeringius acanthophorus* | fusiform | oval | 2.64 | 0.24 | 0.05 | -3.44 | 0.21 | sub-terminal/inferior | lateral | present | heterocercal |
| *Pillararhynhcus longi* | shortAndOrDeep | circular | 3.32 | 0.22 | 0.04 | -2.61 | 0.19 | sub-terminal/inferior | lateral | absent | heterocercal |
| *Pinguosteus thulborni* | shortAndOrDeep | circular | 3.00 | 0.29 | 0.06 | -2.40 | 0.22 | sub-terminal/inferior | lateral | absent | heterocercal |
| *Plourdosteus canadensis* | elongated | circular | 4.57 | 0.19 | 0.02 | -3.35 | 0.18 | sub-terminal/inferior | lateral | absent | heterocercal |
| *Quebecius quebecensis* | fusiform | compressed | 4.09 | 0.21 | 0.03 | -3.40 | 0.18 | terminal | lateral | present | heterocercal |
| *Remigolepis walkeri* | fusiform | angular | 3.61 | 0.09 | 0.02 | -3.72 | 0.24 | sub-terminal/inferior | raised/top of head | absent | heterocercal |
| *Rhinodipterus kimberleyensis* | shortAndOrDeep | circular | 3.40 | 0.25 | 0.05 | -2.30 | 0.18 | sub-terminal/inferior | lateral | absent | heterocercal |
| *Robinsondipterus longi* | fusiform | compressed | 4.26 | 0.24 | 0.03 | -2.12 | 0.23 | sub-terminal/inferior | lateral | absent | heterocercal |
| *Rolfosteus canningensis* | fusiform | circular | 3.78 | 0.26 | 0.04 | -2.08 | 0.08 | sub-terminal/inferior | lateral | absent | heterocercal |
| *Scaumenacia curta* | fusiform | compressed | 4.23 | 0.18 | 0.04 | -2.87 | 0.20 | terminal | lateral | present | heterocercal |
| *Simosteus tuberculatus* | fusiform | oval | 4.17 | 0.19 | 0.03 | -3.39 | 0.23 | terminal | lateral | present | heterocercal |
| *Soederberghia simpsoni* | elongated | compressed | 2.98 | 0.45 | 0.04 | -2.00 | 0.25 | terminal | lateral | absent | heterocercal |
| *Torosteus pulchellus* | fusiform | circular | 3.40 | 0.23 | 0.05 | -3.91 | 0.27 | terminal | lateral | present | heterocercal |
| *Torosteus sp. ?* | fusiform | circular | 3.00 | 0.28 | 0.07 | -2.74 | 0.22 | sub-terminal/inferior | lateral | absent | heterocercal |
| *Torosteus tuberculatus* | fusiform | circular | 3.74 | 0.20 | 0.04 | -3.96 | 0.22 | terminal | lateral | present | heterocercal |

| | | | | | | | | | | | |
|---|---|---|---|---|---|---|---|---|---|---|---|
| *Triazeugacanthus affinis* | elongated | oval | 1.72 | 0.18 | 0.05 | -4.03 | 0.20 | terminal | lateral | absent | heterocercal |
| *Tubonasus lennardensis* | elongated | angular | 3.04 | 0.33 | 0.07 | -2.01 | 0.19 | sub-terminal/inferior | lateral | absent | heterocercal |
| *undescribed chondrich.* | elongated | circular | 2.71 | 0.41 | 0.09 | -2.05 | 0.28 | terminal | lateral | present | heterocercal |
| *Xeradipterus hatcheri* | shortAndOrDeep | circular | 3.86 | 0.19 | 0.03 | -2.91 | 0.11 | sub-terminal/inferior | lateral | absent | heterocercal |

**Table S5.** Hypervolume centroid positions of three Devonian (Gogo, Miguasha, Canowindra) and six modern fish communities on the first seven principal coordinate analysis axes obtained by applying principal coordinates analysis to a Gower dissimilarity matrix of species' traits. Boldface indicates instances where Devonian communities fall outside the range observed in modern communities.

| site | habitat | group | PC1 | PC2 | PC3 | PC4 | PC5 | PC6 | PC7 |
|---|---|---|---|---|---|---|---|---|---|
| Canowindra Billabong | fresh water | temperate/subtropical | **-0.167** | -0.002 | -0.009 | -0.029 | 0.027 | -0.027 | **-0.069** |
| Gogo Reefs | reef | Devonian | **-0.079** | **0.051** | -0.017 | -0.009 | **-0.056** | 0.023 | **-0.06** |
| Miguasha Estuary | estuary | Devonian | **-0.086** | **0.063** | -0.021 | **-0.048** | **-0.001** | -0.008 | **-0.026** |
| Santa Cruz Channel Estuary | estuary | tropical | 0.007 | -0.016 | 0.005 | 0.005 | 0.009 | -0.007 | 0.008 |
| Braço Morto Acima and Abaixo | fresh water | tropical | 0.004 | -0.05 | -0.02 | 0.015 | 0.006 | 0.023 | 0.006 |
| Caribbean Reefs | reef | tropical | 0.014 | -0.005 | 0.029 | -0.002 | 0.008 | -0.005 | 0.011 |
| Chile Reefs | reef | temperate/subtropical | 0.016 | -0.022 | 0.005 | -0.033 | 0.027 | -0.012 | 0.004 |
| Nepean River | fresh water | temperate/subtropical | 0.008 | 0.007 | -0.036 | 0.007 | 0.021 | -0.013 | 0.033 |
| Ythan Estuary | estuary | temperate/subtropical | -0.043 | -0.027 | -0.023 | 0.014 | 0.031 | -0.03 | 0.035 |

**Table S6.** Correlation between species traits and seven principal coordinate analysis axes obtained by applying principal coordinates analysis to a Gower dissimilarity matrix of species' traits.

| trait | axis | test | stat | value | p.value |
|---|---|---|---|---|---|
| eye diameter | PC1 | Linear Model | r2 | 0.622 | 0.000 |
| body depth | PC1 | Linear Model | r2 | 0.523 | 0.000 |
| BodyShapeII (transverse) | PC1 | Kruskal-Wallis | eta2 | 0.359 | 0.000 |
| caudal fin shape | PC1 | Kruskal-Wallis | eta2 | 0.349 | 0.000 |
| BodyShapeI (sagittal) | PC1 | Kruskal-Wallis | eta2 | 0.298 | 0.000 |
| head length | PC1 | Linear Model | r2 | 0.292 | 0.000 |
| mouth position | PC1 | Kruskal-Wallis | eta2 | 0.278 | 0.000 |
| log(total length) | PC1 | Linear Model | r2 | 0.227 | 0.000 |
| log(pre-orbital length) | PC1 | Linear Model | r2 | 0.166 | 0.000 |
| spiracle | PC1 | Kruskal-Wallis | eta2 | 0.159 | 0.000 |
| eye position | PC1 | Kruskal-Wallis | eta2 | 0.022 | 0.002 |
| mouth position | PC2 | Kruskal-Wallis | eta2 | 0.479 | 0.000 |
| eye position | PC2 | Kruskal-Wallis | eta2 | 0.376 | 0.000 |
| BodyShapeI (sagittal) | PC2 | Kruskal-Wallis | eta2 | 0.214 | 0.000 |
| log(total length) | PC2 | Linear Model | r2 | 0.163 | 0.000 |
| spiracle | PC2 | Kruskal-Wallis | eta2 | 0.122 | 0.000 |
| BodyShapeII (transverse) | PC2 | Kruskal-Wallis | eta2 | 0.082 | 0.000 |
| caudal fin shape | PC2 | Kruskal-Wallis | eta2 | 0.06 | 0.000 |
| log(pre-orbital length) | PC2 | Linear Model | r2 | 0.042 | 0.000 |
| head length | PC2 | Linear Model | r2 | 0.005 | 0.104 |
| body depth | PC2 | Linear Model | r2 | 0.005 | 0.119 |
| eye diameter | PC2 | Linear Model | r2 | 0.002 | 0.367 |
| log(pre-orbital length) | PC3 | Linear Model | r2 | 0.499 | 0.000 |
| log(total length) | PC3 | Linear Model | r2 | 0.409 | 0.000 |
| head length | PC3 | Linear Model | r2 | 0.316 | 0.000 |
| BodyShapeII (transverse) | PC3 | Kruskal-Wallis | eta2 | 0.049 | 0.000 |
| BodyShapeI (sagittal) | PC3 | Kruskal-Wallis | eta2 | 0.041 | 0.000 |
| eye diameter | PC3 | Linear Model | r2 | 0.038 | 0.000 |
| spiracle | PC3 | Kruskal-Wallis | eta2 | 0.015 | 0.004 |
| caudal fin shape | PC3 | Kruskal-Wallis | eta2 | 0.007 | 0.170 |
| body depth | PC3 | Linear Model | r2 | 0.002 | 0.316 |
| mouth position | PC3 | Kruskal-Wallis | eta2 | 0 | 0.392 |
| eye position | PC3 | Kruskal-Wallis | eta2 | -0.003 | 0.757 |
| mouth position | PC4 | Kruskal-Wallis | eta2 | 0.34 | 0.000 |
| BodyShapeI (sagittal) | PC4 | Kruskal-Wallis | eta2 | 0.271 | 0.000 |
| eye position | PC4 | Kruskal-Wallis | eta2 | 0.254 | 0.000 |
| BodyShapeII (transverse) | PC4 | Kruskal-Wallis | eta2 | 0.133 | 0.000 |

| | | | | | |
|---|---|---|---|---|---|
| body depth | PC4 | Linear Model | r2 | 0.112 | 0.000 |
| head length | PC4 | Linear Model | r2 | 0.069 | 0.000 |
| spiracle | PC4 | Kruskal-Wallis | eta2 | 0.055 | 0.000 |
| log(total length) | PC4 | Linear Model | r2 | 0.049 | 0.000 |
| caudal fin shape | PC4 | Kruskal-Wallis | eta2 | 0.028 | 0.004 |
| log(pre-orbital length) | PC4 | Linear Model | r2 | 0.027 | 0.000 |
| eye diameter | PC4 | Linear Model | r2 | 0.001 | 0.402 |
| BodyShapeI (sagittal) | PC5 | Kruskal-Wallis | eta2 | 0.571 | 0.000 |
| BodyShapeII (transverse) | PC5 | Kruskal-Wallis | eta2 | 0.151 | 0.000 |
| mouth position | PC5 | Kruskal-Wallis | eta2 | 0.141 | 0.000 |
| caudal fin shape | PC5 | Kruskal-Wallis | eta2 | 0.106 | 0.000 |
| head length | PC5 | Linear Model | r2 | 0.086 | 0.000 |
| eye diameter | PC5 | Linear Model | r2 | 0.082 | 0.000 |
| eye position | PC5 | Kruskal-Wallis | eta2 | 0.029 | 0.000 |
| spiracle | PC5 | Kruskal-Wallis | eta2 | 0.029 | 0.000 |
| log(total length) | PC5 | Linear Model | r2 | 0.011 | 0.019 |
| body depth | PC5 | Linear Model | r2 | 0.006 | 0.087 |
| log(pre-orbital length) | PC5 | Linear Model | r2 | 0 | 0.742 |
| eye position | PC6 | Kruskal-Wallis | eta2 | 0.245 | 0.000 |
| BodyShapeI | PC6 | Kruskal-Wallis | eta2 | 0.23 | 0.000 |
| mouth position | PC6 | Kruskal-Wallis | eta2 | 0.12 | 0.000 |
| BodyShapeII (transverse) | PC6 | Kruskal-Wallis | eta2 | 0.112 | 0.000 |
| log(pre-orbital length) | PC6 | Linear Model | r2 | 0.064 | 0.000 |
| body depth | PC6 | Linear Model | r2 | 0.031 | 0.000 |
| log(total length) | PC6 | Linear Model | r2 | 0.026 | 0.000 |
| caudal fin shape | PC6 | Kruskal-Wallis | eta2 | 0.016 | 0.040 |
| head length | PC6 | Linear Model | r2 | 0.015 | 0.007 |
| eye diameter | PC6 | Linear Model | r2 | 0 | 0.776 |
| spiracle | PC6 | Kruskal-Wallis | eta2 | -0.001 | 0.580 |
| BodyShapeI (sagittal) | PC7 | Kruskal-Wallis | eta2 | 0.348 | 0.000 |
| BodyShapeII (transverse) | PC7 | Kruskal-Wallis | eta2 | 0.262 | 0.000 |
| spiracle | PC7 | Kruskal-Wallis | eta2 | 0.255 | 0.000 |
| body depth | PC7 | Linear Model | r2 | 0.101 | 0.000 |
| caudal fin shape | PC7 | Kruskal-Wallis | eta2 | 0.083 | 0.000 |
| eye position | PC7 | Kruskal-Wallis | eta2 | 0.022 | 0.002 |
| mouth position | PC7 | Kruskal-Wallis | eta2 | 0.011 | 0.027 |
| log(pre-orbital length) | PC7 | Linear Model | r2 | 0.01 | 0.025 |
| log(total length) | PC7 | Linear Model | r2 | 0.004 | 0.185 |
| eye diameter | PC7 | Linear Model | r2 | 0.002 | 0.284 |
| head length | PC7 | Linear Model | r2 | 0.001 | 0.404 |
